# Factors that Influence Career Choice Among Different Populations of Neuroscience Trainees

**DOI:** 10.1101/2021.04.12.439464

**Authors:** Lauren E. Ullrich, John Ogawa, Michelle D. Jones-London

## Abstract

Enhancing the diversity of the scientific workforce is critical to achieving the mission of NIH: “To seek fundamental knowledge about the nature and behavior of living systems and the application of that knowledge to enhance health, lengthen life, and reduce the burdens of illness and disability.” However, specific groups have historically been, and continue to be, underrepresented in the biomedical research workforce, especially academia. Career choice is a multi-factorial process that evolves over time; among all trainees, expressed interest in faculty research careers decreases over time in graduate school, but that trend is amplified in women and members of historically underrepresented racial and ethnic groups (Fuhrmann, Halme, O’Sullivan, & Lindstaedt, 2011; Gibbs, McGready, Bennett, & Griffin, 2014; C. Golde & Dore, 2004; Roach & Sauermann, 2017; Sauermann & Roach, 2012). Neuroscience as a discipline has characteristics that may exacerbate the overall trends seen in the life sciences, such as a greater growth in the number of awarded neuroscience PhDs than in other life sciences fields (US National Science Foundation, 2016b). This work was designed to investigate how career interest changes over time among recent neuroscience PhD graduates, and whether differences in career interests are associated with social identity (i.e. gender and race/ethnicity), experiences in graduate school and postdoctoral training (e.g. relationship with advisor; feelings of belonging), and personal characteristics (e.g. confidence in one’s potential to be an independent researcher). We report results from a survey of 1,479 PhD neuroscientists (including 16% underrepresented (UR) and 54% female scientists). We saw repeated evidence that individual preferences about careers in general, and academic careers specifically, predict current career interest. These statistically significant preferences mostly had medium to low effect size that varied by career type. These findings were mediated by social identity and experiences in graduate school and postdoctoral training. Our findings highlight the important influence of the advisor in shaping a trainee’s career path, and the ways in which academic culture is perceived as unwelcoming or incongruent with the values or priorities of certain groups. For women, issues of work/life balance and structural issues of academia, and for UR women in particular, lower confidence in their ability to be an independent researcher, affected their interest in academia. Both women and UR men in our study report a lower importance of autonomy in their careers. UR respondents report feeling less like they were a part of the social and intellectual community. However, they have formed beneficial relationships with faculty outside their PhD institutions that, particularly for UR women, are associated with increased interest in academia. Our findings suggest several areas for positive growth, ways to change how we think about the impact of mentorship, and policy and programmatic interventions that extend beyond trying to change or “fix” the individual and instead recognize the systemic structures that influence career choices.

## INTRODUCTION

During the past few decades, the biomedical sciences career track has undergone significant changes. The number of doctorate recipients in the life sciences has nearly doubled in the past 30 years, while the number of tenure-track faculty appointments 3-5 years after graduation has remained flat (Larson, Ghaffarzadegan, & Xue, 2014; Roach & Sauermann, 2017). Consistent with this trend, the share of life science PhDs holding faculty positions has declined: in 1993, 17.3% of life science PhD graduates held a tenure-track position 3-5 years after graduation; by 2013, that percentage was 10.6% (National Science Board, 2016). This sea change in career prospects and outcomes has sparked a national conversation within the scientific community about how trainees make career choices and how to best prepare them for their future careers (Gibbs & Griffin, 2013; Gibbs et al., 2014; National Institutes of Health, 2012; M. V. Sinche, 2016).

As a discipline, Neuroscience has characteristics that may exacerbate the overall trends seen in the life sciences. For example, the number of trainees in neuroscience has been expanding much more rapidly than other fields. Between 1990 and 2013, the number of neuroscience PhDs awarded increased more than 5-fold—in comparison, the number of biological PhDs awarded increased 2-fold (US National Science Foundation, 2016b). Consequently, the magnitude of the difference between the number of trainees and faculty positions in neuroscience is likely greater than biology and biomedicine overall. This is supported by the fact that in 2017, 5% of neurobiology and neuroscience PhD holders who were 3-7 years after graduation held a tenured or tenure-track position, compared to 10% of comparable biological, agricultural, and environmental life sciences PhDs as a group (calculated from National Science Foundation, 2020). This may lead students and/or faculty to place more emphasis on preparation for careers other than research faculty positions. An example of a program created to respond to this trend is the NIH Broadening Experiences in Scientific Training (BEST) initiative, which prepares individuals for a broader range of careers in the biomedical research enterprise (Lenzi, Korn, Wallace, Desmond, & Labosky, 2020). This kind of program presents alternatives to trainees who are currently spending longer time periods as postdoctoral fellows competing for limited faculty positions.

Historically, specific groups have been, and continue to be, underrepresented in science and technology. These underrepresented groups include American Indians/Alaska Natives; Blacks/African Americans; Hispanics/Latinos; Native Hawaiians/Other Pacific Islanders and persons with disabilities (US National Science Foundation, 2016b). These groups are underrepresented at every level of postsecondary education, and underrepresentation is progressively greater at every rung on the academic ladder (National Academy of Sciences (US), 2011). For example, in 2016, scientists that belong to underrepresented racial and ethnic groups received 14% of life science doctoral degrees, but made up only 10% of tenured and tenure-track life science faculty in the United States (US National Science Foundation, 2016a, 2016b).

Enhancing the diversity of the scientific workforce is critical to “fostering scientific innovation, enhancing global competitiveness, contributing to robust learning environments, improving the quality of the research, advancing the likelihood that underserved or health disparity populations participate in, and benefit from health research, and enhancing public trust” (National Institutes of Health, 2019b).

However, decades of efforts by the National Institutes of Health (NIH) and others to “enhance the pipeline” by increasing entry of women and underrepresented scientists into undergraduate and PhD science programs has not had an appreciable effect on the relative proportion of underrepresented tenure-track faculty (Gibbs, Basson, Xierali, & Broniatowski, 2016). This has commonly been described as “the leaky pipeline” (Miller & Wai, 2015), a metaphor that assumes that at PhD entry all trainees aspire to a faculty research position, but some “leak” from the faculty pipeline into a different career. The framing has evolved over time from a discussion about the “pipeline” to a recognition of different “pathways” to science (K. A. Griffin, 2016). It is possible that some trainees either never wanted a faculty research position or were interested in a variety of professions. Alternatively, trainees may be or feel forced out of the faculty track by a lack of opportunities, an unwelcoming academic culture, or other circumstances beyond their control.

Moreover, compared to well-represented students, women and people from underrepresented groups (including and perhaps especially those with multiple underrepresented or marginalized identities) face additional or unique challenges in training, and may make different career choices based on their experiences and values (Fisher et al., 2019; Gibbs & Griffin, 2013; Sauermann & Roach, 2012). Career choice is a multi-factorial process that evolves over time; among all trainees, expressed interest in faculty research careers decreases over time in graduate school, but that trend is amplified in women and members of U.S-based historically underrepresented racial and ethnic groups (Fuhrmann et al., 2011; Gibbs et al., 2014; C. Golde & Dore, 2004; Roach & Sauermann, 2017; Sauermann & Roach, 2012). Previous research has shown that factors such as research self-efficacy (confidence in one’s ability as a researcher); social and intellectual feeling of belonging; and interactions with one’s advisor can all affect career interests, particularly among underrepresented scientists (Estrada, Woodcock, Hernandez, & Schultz, 2011; Estrada, Zhi, Nwankwo, & Gershon, 2019; Gazley et al., 2014; Gibbs et al., 2014; Hayter & Parker, 2018).

In order to create and administer effective training programs for a diverse research workforce, in 2017, the National Institute of Neurological Disorders and Stroke (NINDS) sought information about the factors influencing career choice among different populations, particularly those underrepresented in the neuroscience workforce. This work was designed to investigate how career interest changes over time among recent neuroscience PhD graduates, and whether differences in career interests are associated with social identity (i.e. gender and race/ethnicity), experiences in graduate school and postdoctoral training (e.g. relationship with advisor; feelings of belonging), and personal characteristics (e.g. confidence in one’s potential to be an independent researcher). NINDS sought input from current or recent trainees in the neuroscience field to help inform future training programs and initiatives to better serve the neuroscience community. While the COVID-19 pandemic has created unprecedented challenges worldwide and provided opportunities for long-overdue public conversations about structural racism that have affected neuroscientists at all career stages, this survey is a snapshot in time and does not capture the current conditions. However, the academic culture and systemic training environments highlighted in the survey remain relevant. NINDS is committed to the development of a biomedical research workforce that is representative of the diversity in American society and the information collected from this study was aimed to help give NINDS and the entire neuroscience community a clearer picture of the environment and experiences of our trainee and potential trainee community.

## RESULTS

### Sample Demographics

Table S1 in the *Supporting Information Appendix* presents basic demographic information about the sample of 1,479 PhD neuroscientists who responded to the survey. We solicited responses from all recent doctoral recipients (CY2008 or later) who were US citizens or permanent residents and had applied for NINDS funding or have been appointed to NINDS training (T32) or research education grants (R25) between 2003 and 2017. Respondent information included gender (54% female, n=793), underrepresented (UR) status (16% UR, n=233), and social identity (9% female UR, n=133; 7% male UR, n=100; 45% female well-represented (WR), n=660; and 40% male WR, n=586). 41% of participants were in a postdoctoral position, 27% in research-focused academic positions, 10% in science-related, non-research positions, 9% in non-academic research positions, 7% in teaching-focused academic positions, 4% in non-science positions, and 2% were unemployed. 48% held a PhD in neuroscience, the rest were in biological or health-related fields. 251 PhD institutions were represented in 47 states, the District of Columbia, and Puerto Rico. Additional descriptive statistics for all dependent and independent variables in the study are presented in Table S2 in the *Supporting Information Appendix*.

### Approach

After characterizing basic demographic information, we first investigated whether there were differences by social identity in factors likely to influence career interest, such as experiences in graduate school and postdoctoral training, personal characteristics, and objective measures. Next, we looked at whether there were changes in interest in the four career types (research-focused academic faculty positions, teaching-focused academic faculty positions, non-academic research positions, and science-related, non-research positions) over time across the whole sample and by social identity. We then performed two different sets of follow-up analyses on career interests. The first set of analyses investigated which factors predicted changes in interest in the four career types over the course of graduate school. The second set of analyses investigated which factors predicted *current* interest in the four career types.

### Differences by Gender and UR in Explanatory Variables

First, we asked whether experiences in graduate school and postdoctoral training, personal characteristics, and objective measures differed by social identity in our sample (full results of the analyses to investigate differences by Gender and UR status are presented in Table S3 in the *Supporting Information Appendix)*. Significant findings were followed-up by examining differences either in means or slopes for subsamples defined by whichever was significant of gender, UR status, or their interaction (Table S4 in the *Supporting Information Appendix*). Definitions for small, medium and large effect sizes for this article are generally taken from Cohen (1988): mean differences use d (small (s) ≥ 0.2, medium (m) ≥ 0.5, large (l) ≥ 0.8), and correlation coefficients and individual regression coefficients use r (s ≥ 0.1, m ≥ 0.3, l ≥ 0.6). Finally, for odds ratios we used (rounded) Cohen’s cutoffs (s ≥ 1.5, m ≥ 2.5, l ≥ 4.5).

The follow-up analyses revealed several differences by Gender in Current Position (Figure 1 and Table S4a). A higher proportion of males than females were in research-focused academic faculty positions. Conversely, a higher proportion of females than males were in teaching-focused academic faculty positions and science-related, non-research positions.

**Figure 1.**
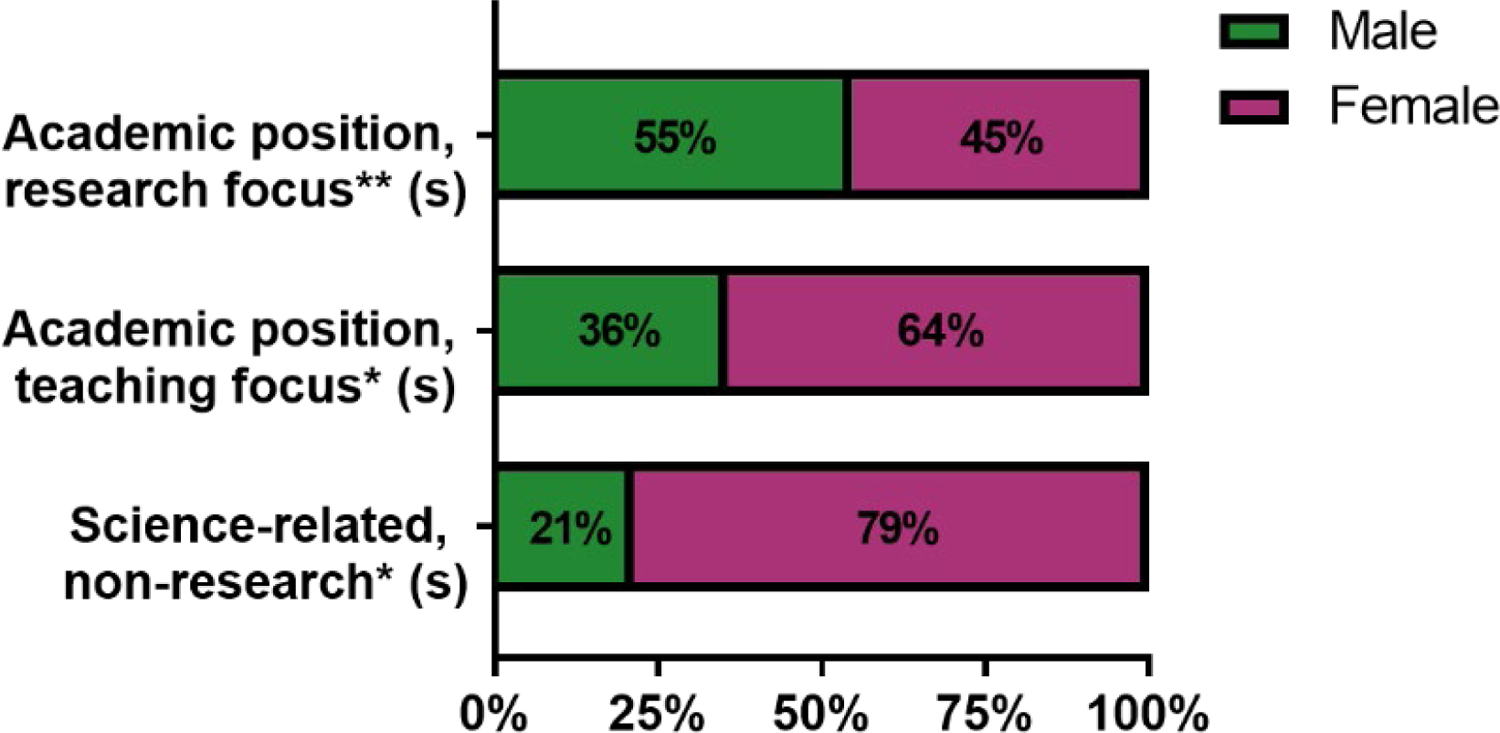
Gender differences in current position among PhD neuroscientists. Proportion of females and males in the sample by job sector of their current position. Significance levels from Chi-squared statistics (Table S4a). (s) = small effect size, * = p < 0.05, ** = p < 0.01.

We also found differences in experiences and personal characteristics between males and females (Figure 2, Table S4b). Females’ assessments of their relationships with their PhD program advisors were significantly lower than those from males, as reflected by the negative z-scores. Females also report a significantly lower publication rate than males. In addition, females’ current ratings of their confidence in their potential to be independent researchers were significantly lower than males. For the factors assessing the importance of different aspects of careers, females rated the “Autonomy” factor less important and the “Work/Life Balance” factor more important than males. Finally, for the factors assessing whether different “features of academia” increased or decreased interest in academia, females reported that the “Funding/Job Market/Promotion” factor and the “Research/Autonomy” factor decreased their interest in academia more than males.

**Figure 2.**
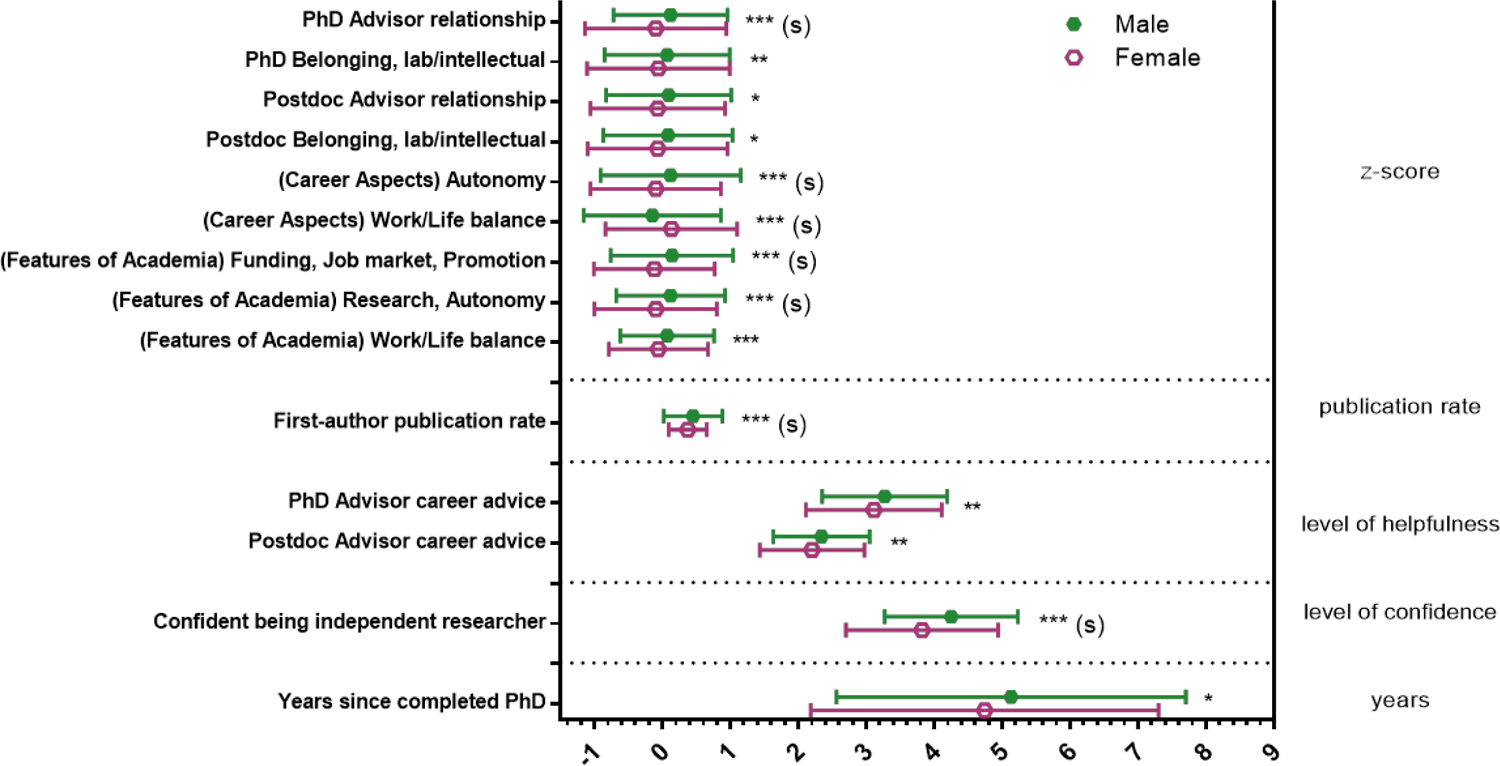
Gender differences in experiences and characteristics among PhD neuroscientists. Mean responses to variables capturing experiences, personal characteristics, and objective measures. Significance levels from F statistics (ANOVA, Table S4b) comparing the means for females and males for each variable were all significant at p < 0.05, at least. Responses on the X-axis were z-scores for the top variables; total publications/years of research for publication rate; level of helpfulness (1-4, 4 being very helpful) for the career advice variables; level of confidence (1-5, 5 being most confident) for confidence in being an independent researcher; and years for years since completed PhD. Effect sizes are labeled when they reach at least “small” size. (s) = small effect size, * = p < 0.05, ** = p < 0.01, *** = p < 0.001

WR and UR respondents also differed on several variables. We found that WR respondents were far less likely than UR respondents to have been the first person or in the first generation of their family to graduate from a 4-year college or university (Figure 3, Table S4c). We also found differences between UR and WR respondents in experiences and personal characteristics (Figure 4, Table S4d). UR respondents reported feeling more support from faculty outside their institutions during their PhD programs than WR respondents. Conversely, UR respondents had lower scores on the factor that captured feelings of belonging intellectually/socially to their postdoc research group than WR respondents. UR respondents also reported lower publication rates than WR respondents. For the factors assessing the importance of different aspects of careers, UR respondents reported lower importance of the “Autonomy” factor than WR respondents. Finally, for the factors assessing the influence of different features of academia, UR respondents reported that the “Work/Life Balance” factor increased their interest in academia more than WR respondents.<colcnt=1>

**Figure 3.**
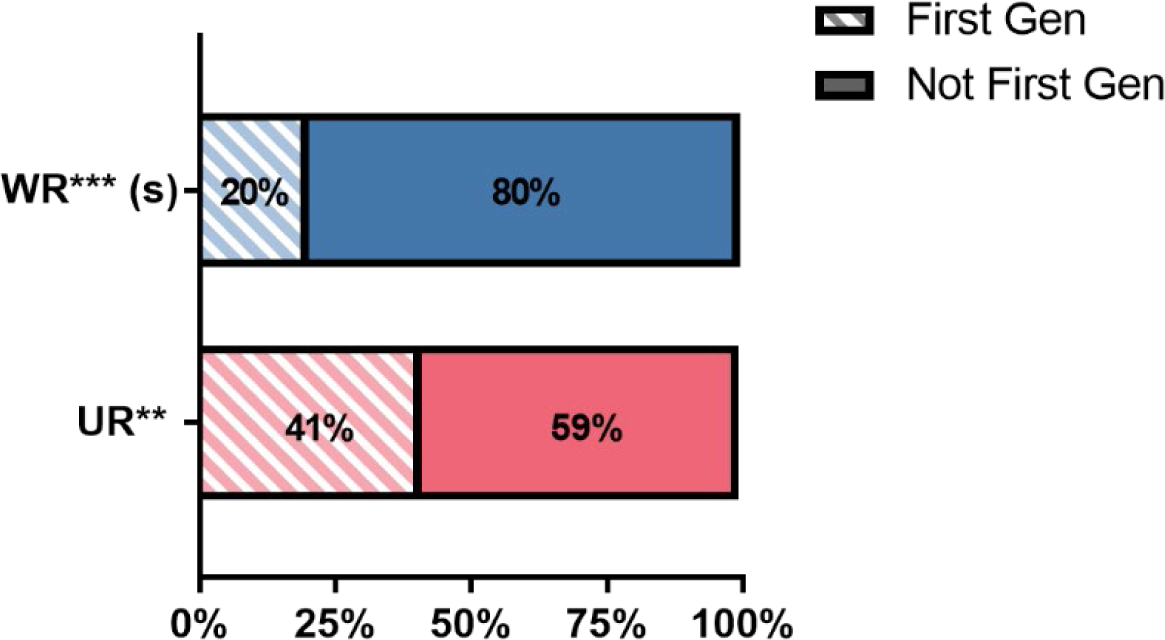
WR respondents were less likely to be first generation college students than UR respondents. Proportion of respondents who were the first person or in the first generation in their family to graduate from a 4-year college by UR status. Significance levels from Chi-squared statistics. (Table S4c). Effect sizes are labeled when they reach at least “small” size. (s) = small effect size; ** = p < 0.01, *** = p < 0.001

**Figure 4.**
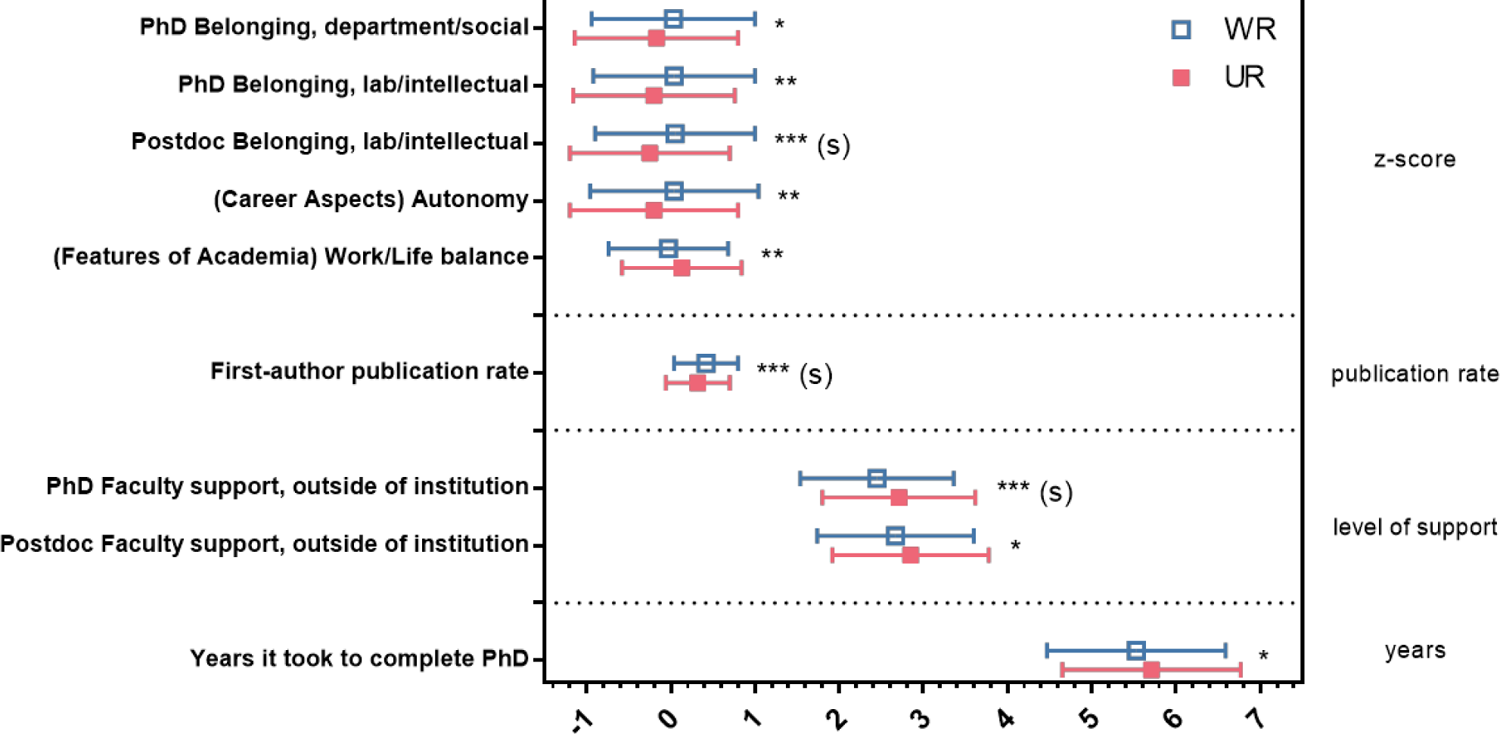
UR status differences in experiences and characteristics among PhD neuroscientists. Mean responses to variables capturing experiences, personal characteristics, and objective measures. Significance levels from F statistics (ANOVA, Table S4d) comparing the means for UR respondents and WR respondents for each variable were all significant at p < 0.05. Responses on the X-axis were z-scores for the top variables; total publications/years of research for publication rate; level of helpfulness (1-4, 4 being very helpful) for the outside faculty support variable; and years for years it took to complete PhD. Effect sizes are labeled when they reach at least “small” size. (s) = small effect size; * = p < 0.05, ** = p < 0.01, *** = p < 0.001

At the intersection of social identity (Gender and UR status), we found a 2-way interaction of Gender and UR status for feelings of belonging in both the PhD research group and PhD department (Figure 5, Table S4e). Although there was no difference between WR and UR males, UR females reported lower feelings of belonging than WR females on both the factor that captured feelings of intellectual belonging to their PhD lab/research group and the factor that captured feelings of social belonging to their PhD department.

**Figure 5.**
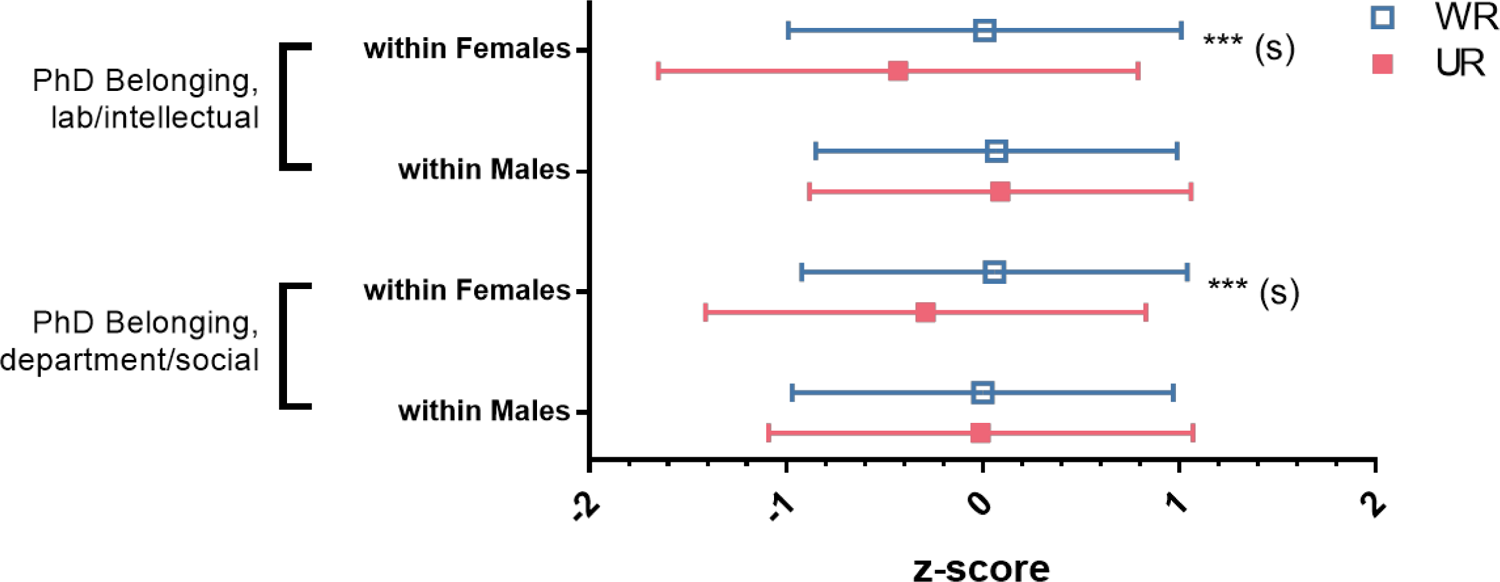
UR females feel a lower sense of belonging than WR females, with no difference for males. Mean responses split by gender and UR status on their feelings of belonging to their PhD lab/research group or department (reduced to 2 single factors with factor analysis). Significance levels from F statistics (ANOVA, Table S4e). Effect sizes are labeled when they reach at least “small” size. (s) = small effect size; * = p < 0.05, ** = p < 0.01, *** = p < 0.001

### Changes in Career Interest Over Time

We were interested in whether there were changes in the four career interest ratings over time across the entire sample. Using repeated measures MANOVAs, we found significant main effects for time for all four career types (see Table S5a in the *Supporting Information Appendix* and Figure 6): decreases over time for interest in research-focused academic faculty positions and teaching-focused academic faculty positions, and increases over time for non-academic research positions and science-related, non-research positions. This reflects similar trends as other studies: interest in academia, both research- and teaching-focused positions, goes down over time while interest in non-academic careers goes up over time (Fuhrmann et al., 2011; Gibbs et al., 2014; Gibbs, McGready, & Griffin, 2015; C. M. Golde & Dore, 2001; Goulden, Mason, & Frasch, 2011; Roach & Sauermann, 2017; Sauermann & Roach, 2012; but see: Wood et al., 2020).

**Figure 6.**
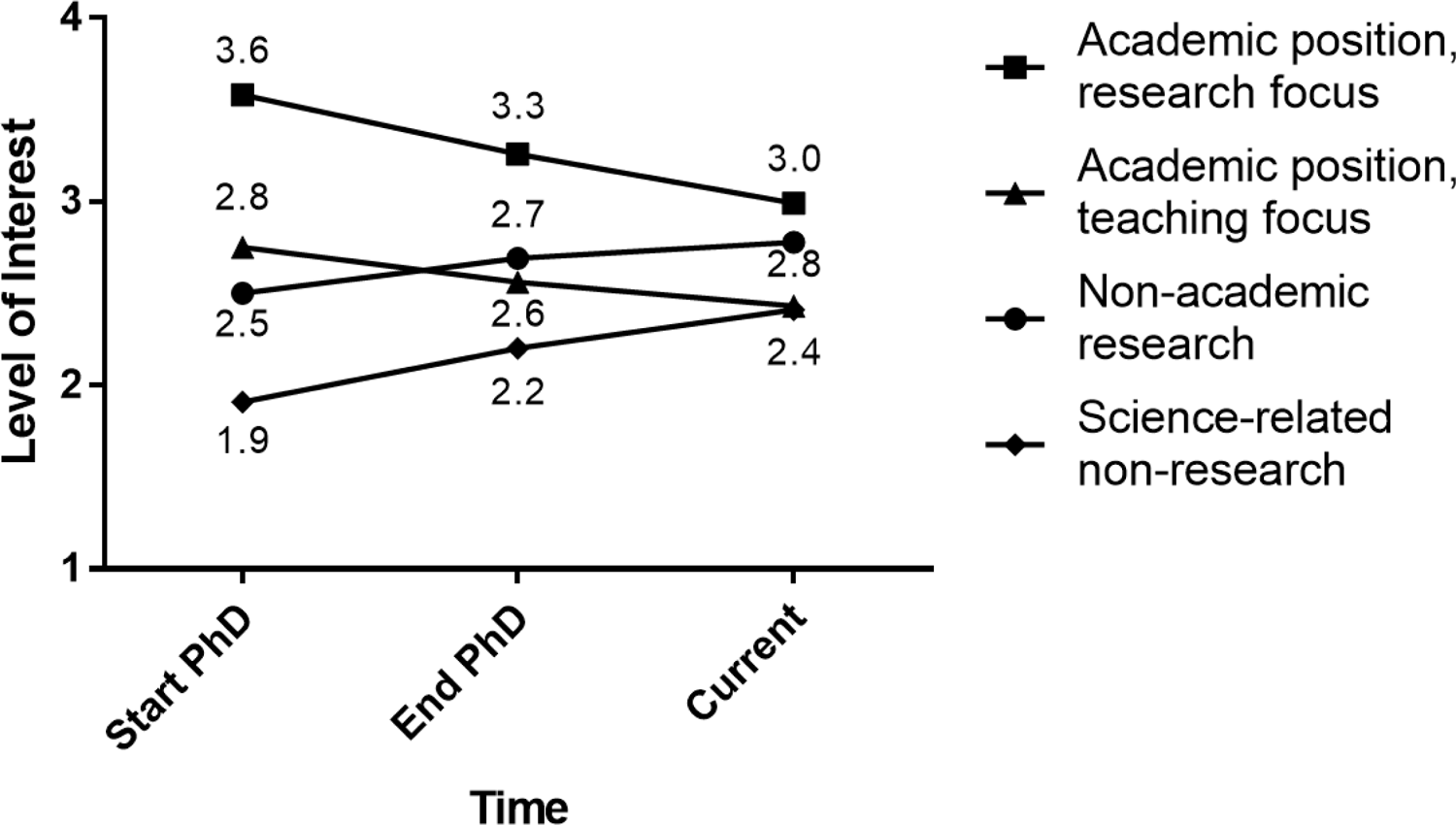
Change in career interest ratings over time among PhD neuroscientists. Mean responses of 1,479 PhD neuroscientists who were asked to rate their level of interest in four different career paths at three times: Start of PhD (T1), End of PhD (T2), and Current (T3), on a 4-point scale (where 1 represents “no interest” and 4 represents “strong interest”). Repeated measures MANOVAs found T1 v T2 was significant at p < 0.001 for all careers; T2 v T3 was significant at p < 0.001 for Academic Faculty, research focus and Science-related non-research, and p < 0.01 for academic faculty, teaching focus and non-academic research (Table S5a).

To determine whether there were differences in trajectory of interest over time by social identity, follow-up analyses were conducted for significant interactions with Gender and UR status (see Table S5b in the *Supporting Information Appendix)*. For interest in research-focused academic faculty positions, the “Time by Gender” interaction indicated that females’ interest in research-focused academic faculty positions was lower at the start of training and decreased over time at a higher rate than males’ interest (Figure 7). For interest in teaching-focused academic faculty positions, the “Time by UR Status” interaction indicated that well-represented participants’ interest in teaching-focused academic faculty positions decreased over time at a higher rate than UR participants’ interest (Figure 8). Finally, for interest in science-related, non-research positions, the “Time by Gender” interaction indicated that females’ interest in science-related, non-research positions increased over time at a higher rate than males’ interest (Figure 9).<colcnt=1>

**Figure 7.**
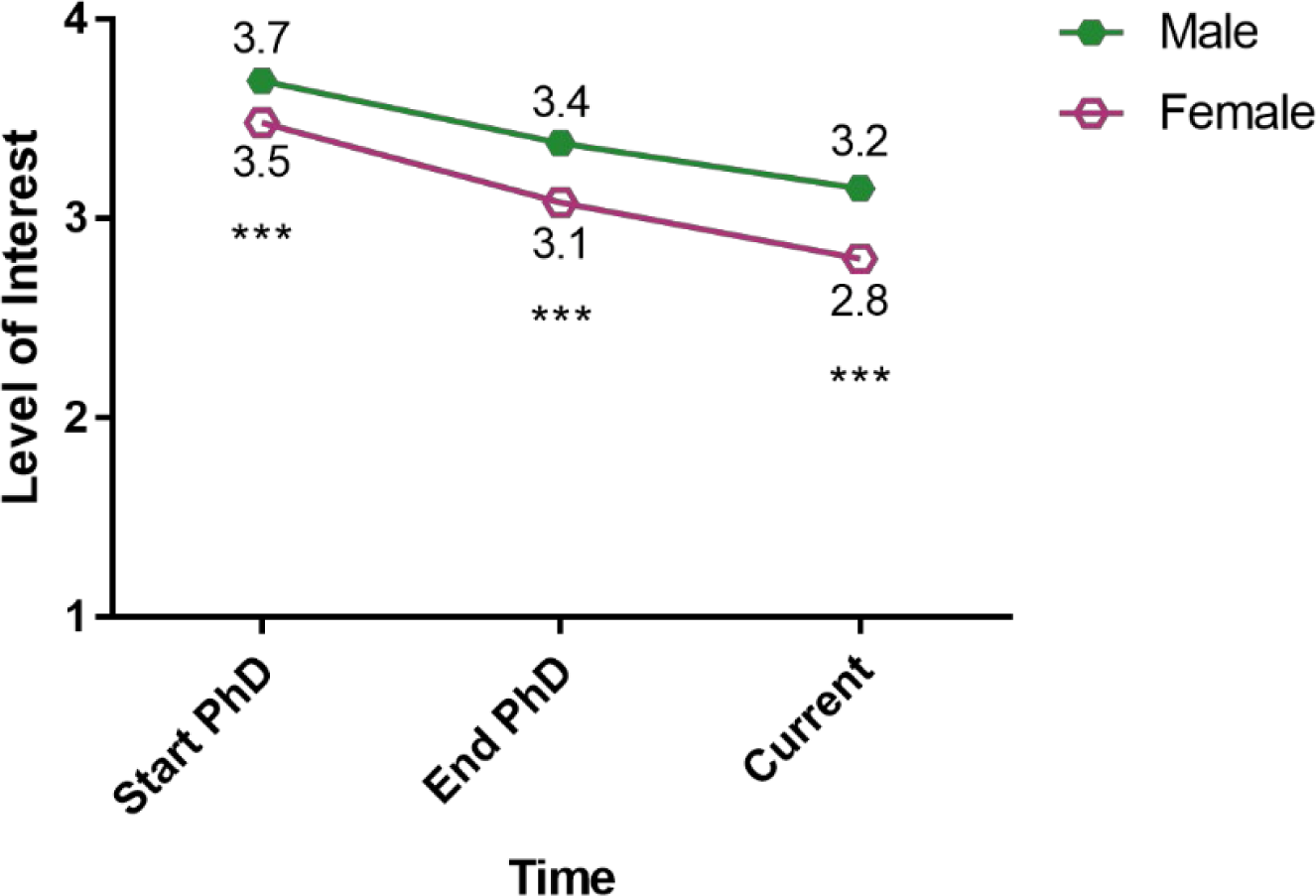
Females less interested in academic research positions than males at PhD start and over time, and interest decreased at a higher rate. Mean responses split by gender on level of interest in research-focused academic faculty positions at Start of PhD (T1), End of PhD (T2),and Current (T3) on a 4-point scale (where 1 represents “no interest” and 4 represents “strong interest”). Omnibus repeated measures MANOVA found an interaction between time and gender. Significance levels from follow-up ANOVAs (Table S5b). *** = p < 0.001

**Figure 8.**
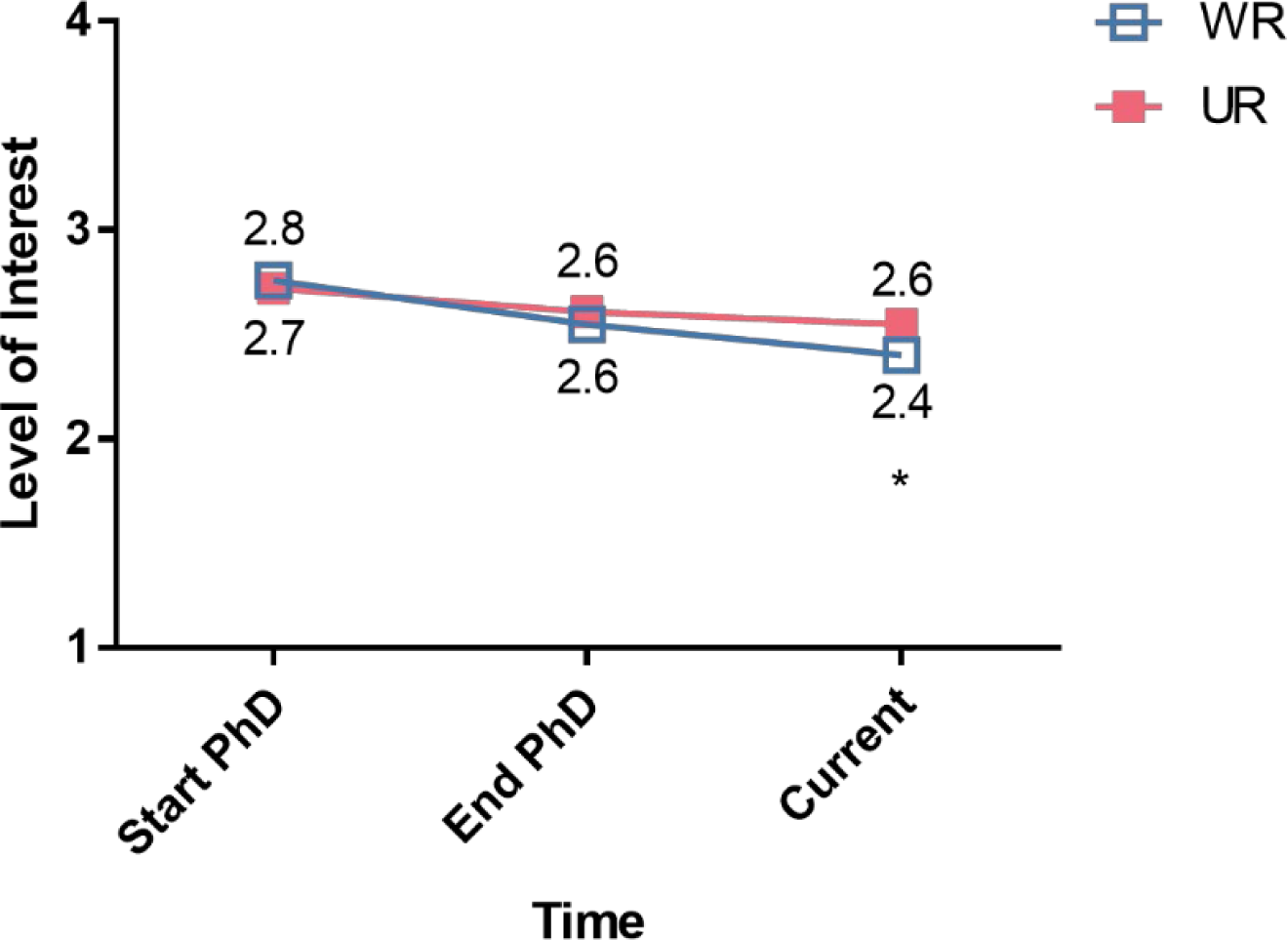
WR respondents less interested in academic teaching positions following PhD completion. Mean responses split by UR status on level of interest in teaching-focused academic faculty positions at Start of PhD (T1), End of PhD (T2), and Current (T3) on a 4-point scale (where 1 represents “no interest” and 4 represents “strong interest”). Omnibus repeated measures MANOVA found an interaction between time and UR status. Significance levels from follow-up ANOVAs (Table S5b). * = p < 0.05

**Figure 9.**
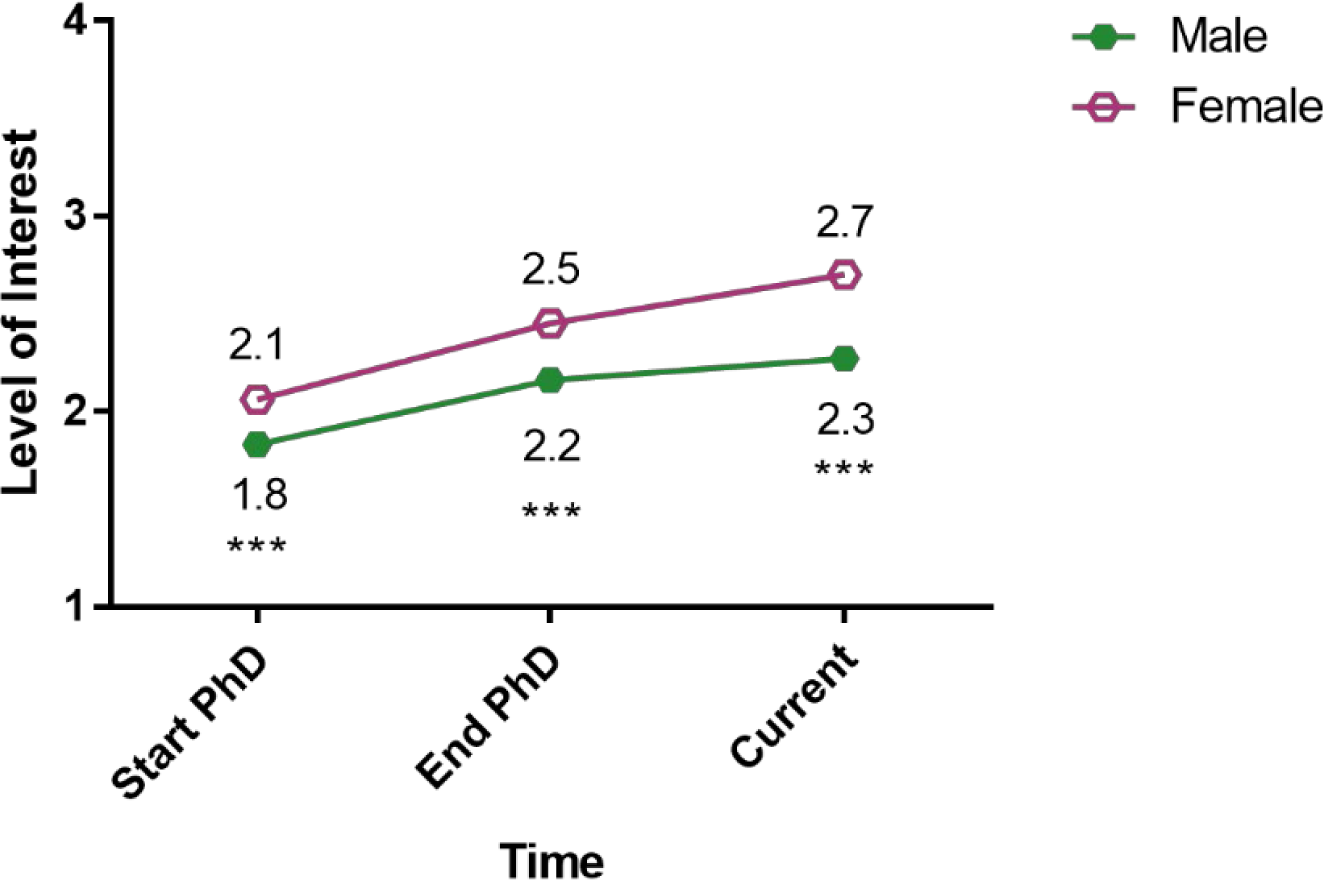
Females more interested in science-related, non-research positions than males at PhD start and over time, and their interest increased at a higher rate. Mean responses split by gender on level of interest in science-related, non-research positions at Start of PhD (T1), End of PhD (T2), and Current (T3) on a 4-point scale (where 1 represents “no interest” and 4 represents “strong interest”). Omnibus repeated measures MANOVA found an interaction between time and gender. Significance levels from follow-up ANOVAs (Table S5b). *** = p < 0.001

### Predicting Change in Interest

Next, we were interested in determining which, if any, factors predicted changes in career interest over the course of a PhD program. For this analysis, we chose predictor variables that captured issues that were contemporaneous with participants’ time in PhD training. Descriptions of the procedures can be found in the Methods section, and preliminary steps and results for construction of the regression are reported in Tables S6 and S7 in the *Supporting Information Appendix*.

The final step in this analysis was to regress each of the four career interest ratings at end of graduate school (T2) on: 1) the interest rating for the same career at start of graduate school (T1), 2) the variables that had significant correlations with it in the correlation step (Table S6), and 3) the interactions that were significant predictors of it in the interaction test step (Table S7). Because the T1 interest rating is in the equation simultaneously with the other variables, it is interpreted as predicting change in interest during graduate school. The results of these analyses are presented in Table 1. We discuss the regression for each career type in the following sub-sections. Discussion of each regression includes only coefficients/variables that were significant and had at least a small effect size. Significant main effects in the context of interactions are not discussed, and significant lower-level interactions in the context of higher-level interactions are not discussed fully, because their meaning is difficult to determine in that context.

**Table 1.** Regressions Predicting Career Interest Ratings at T2 (End of PhD) from Graduate School Era Explanatory Variables. Results of regressions for each of the four career interest T2 ratings (end of PhD) on T1 (start of PhD) interest rating, significantly correlated variables (Table S6), and significant predictor interactions (Table S7). Unstd. Coeff. = Unstandardized coefficient; sig. = significance. Effect sizes are labeled when they reach at least “small” size. (S) = small effect size, (M) = medium effect size, (L) = large effect size. * = p < 0.05; ** = p < 0.01; *** = p < 0.001

| Independent Variables (Graduate School Era Explanatory) |  | Dependent Variables<br>T2 (End of PhD) Career Interest Ratings |  |  |  |  |  |  |  |  |  |  |  |
| --- | --- | --- | --- | --- | --- | --- | --- | --- | --- | --- | --- | --- | --- |
|  |  | Academic Faculty/Research |  |  | Academic Faculty/Teaching |  |  | Non-academic Research |  |  | Science/Non-research |  |  |
|  |  | Unstd. Coeff. | Sig. | Effect size | Unstd. Coeff. | Sig. | Effect size | Unstd. Coeff. | Sig. | Effect size | Unstd. Coeff. | Sig. | Effect size |
| Variables in the Equation | (Intercept) | 2.682 | *** |  | 2.359 | *** |  | 2.662 | *** |  | 2.442 | *** |  |
|  | T1 (Start of PhD) Interest Rating | 0.597 | *** | M (0.174) | 0.696 | *** | L (0.421) | 0.536 | *** | M (0.148) | 0.766 | *** | L (0.323) |
|  | T1 Interest Rating -BY- Gender |  |  |  |  |  |  | 0.132 | ** | - |  |  |  |
|  | T1 Interest Rating -BY- UR Status |  |  |  |  |  |  |  |  |  | -0.135 | * | - |
|  | PhD Belonging, lab | 0.091 | ** | - |  |  |  |  |  |  |  |  |  |
|  | PhD Belonging, lab -BY- Gender |  |  |  |  |  |  |  |  |  |  |  |  |
|  | PhD Belonging, lab -BY- UR Status |  |  |  |  |  |  |  |  |  |  |  |  |
|  | PhD Belonging, lab -BY- Gender -BY- UR Status |  |  |  |  |  |  |  |  |  |  |  |  |
|  | PhD Advisor career advice | 0.235 | *** | S (0.045) | 0.113 | *** | - |  |  |  | -0.124 | *** | S (0.014) |
|  | PhD Advisor career advice -BY- UR Status |  |  |  | -0.115 |  |  |  |  |  |  |  |  |
|  | PhD Faculty support, at institution |  |  |  | -0.010 |  |  |  |  |  |  |  |  |
|  | PhD Faculty support, at institution -BY- Gender |  |  |  | 0.022 |  |  |  |  |  |  |  |  |
|  | PhD Faculty support, at institution -BY- UR Status |  |  |  | 0.089 |  |  |  |  |  |  |  |  |
|  | PhD Faculty support, at institution -BY- Gender -BY- UR Status |  |  |  | -0.311 | * | - |  |  |  |  |  |  |
|  | PhD Faculty support, outside institution | 0.008 |  |  |  |  |  |  |  |  |  |  |  |
|  | PhD Faculty support, outside institution -BY- Gender | 0.029 |  |  |  |  |  |  |  |  |  |  |  |
|  | PhD Faculty support, outside institution -BY- UR Status | 0.124 |  |  |  |  |  |  |  |  |  |  |  |
|  | PhD Faculty support, outside institution -BY- Gender -BY- UR | -0.304 | * | - |  |  |  |  |  |  |  |  |  |
|  | Gender | 0.097 |  |  | -0.121 |  |  | 0.049 |  |  |  |  |  |
|  | UR | -0.266 |  |  | 0.085 |  |  |  |  |  | 0.236 | *** | - |
|  | Gender -BY- UR | 0.446 |  |  | 0.792 | * | - |  |  |  |  |  |  |
| Model Statistics | R<sup>2</sup> | 0.309 |  |  | 0.436 |  |  | 0.339 |  |  | 0.419 |  |  |
|  | Adjusted R<sup>2</sup> | 0.304 |  |  | 0.432 |  |  | 0.337 |  |  | 0.417 |  |  |
|  | F | 65.68 |  |  | 113.29 |  |  | 251.81 |  |  | 265.55 |  |  |
|  | df | 11 |  |  | 11 |  |  | 4 |  |  | 5 |  |  |
|  | df.residual | 1468 |  |  | 1468 |  |  | 1475 |  |  | 1474 |  |  |
|  | p | 0.0000 |  |  | 0.0000 |  |  | 0.0000 |  |  | 0.0000 |  |  |
|  | Non-T1-Interest Variance | 0.14 |  |  | 0.01 |  |  | 0.19 |  |  | 0.10 |  |  |

### Factors that Predict Changes in Interest in Research-focused Academic Positions

First, we predicted change in respondents’ ratings of interest in research-focused academic faculty positions from the start of graduate school (T1) to the end of graduate school (T2). The full regression equation (including all predictors in Figure 10) was itself significant, accounting for 30.4% of the variance in interest in research academia at T2 (Adjusted R^2^, Table 1). Note that interest at the start of graduate school predicted a modest 17.4% of the variance. Helpfulness of career advice from PhD advisor (4.5% variance) was the only other significant predictor with sufficient effect size to report. Higher ratings of the helpfulness of career advice from PhD advisors was also related to greater interest in academic research careers during graduate school. Although other single variables were significant predictors, their effect sizes did not meet the threshold for reporting.

**Figure 10.**
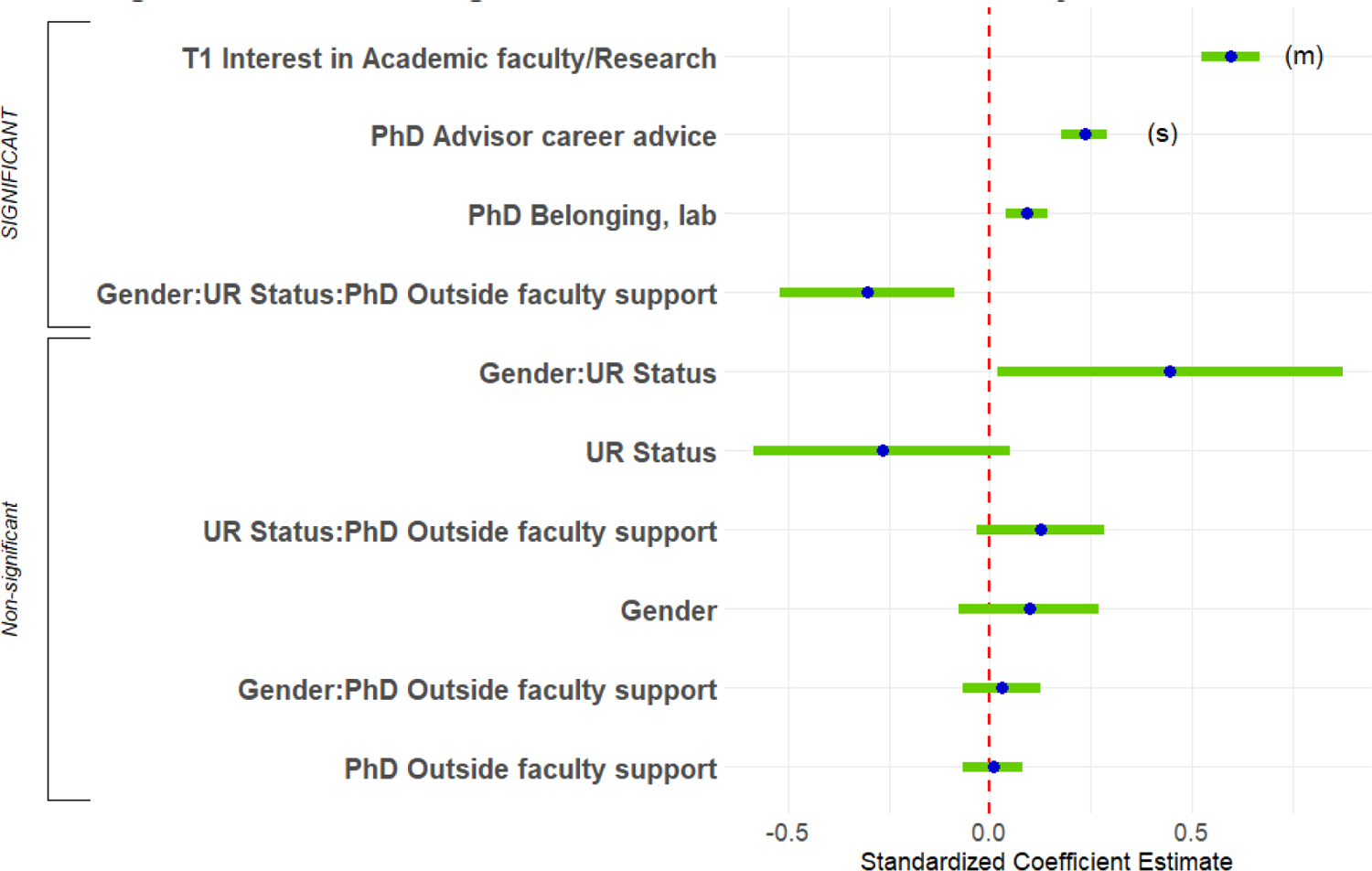
Predicting end of PhD interest in research-focused academic faculty positions. Standardized regression coefficients and error bars for linear regression predicting interest at the end of PhD training (T2) in research-focused academic faculty positions. Dependent variables were level of interest at the end of PhD training on a 4-point scale (where 1 represents “no interest” and 4 represents “strong interest”). Independent variables captured level of interest at the start of PhD training (T1), experiences during PhD training, personal characteristics, objective measures, and interactions with gender and UR status. The entire equation was significant at p < 0.001 and captured 30.4% of the variance (adjusted; Table 1). Effect sizes are labeled when they reach at least “small” size. (s) = small effect size, (m) = medium effect size

The regression analysis identified a significant 3-way interaction among Gender, UR status, and level of support from faculty outside respondents’ institutions during their PhD program (hereafter, “PhD outside faculty support”). The interaction, and all other 3-way interactions in this paper, was followed-up by testing gender differences in prediction of the dependent variable by the explanatory variable within levels of UR status. Although higher support from outside faculty was related to an increase in interest in academic research over the course of graduate school for UR females, it was related to a decrease in interest for UR males (Figure 11; there was no relation for WRs).

**Figure 11.**
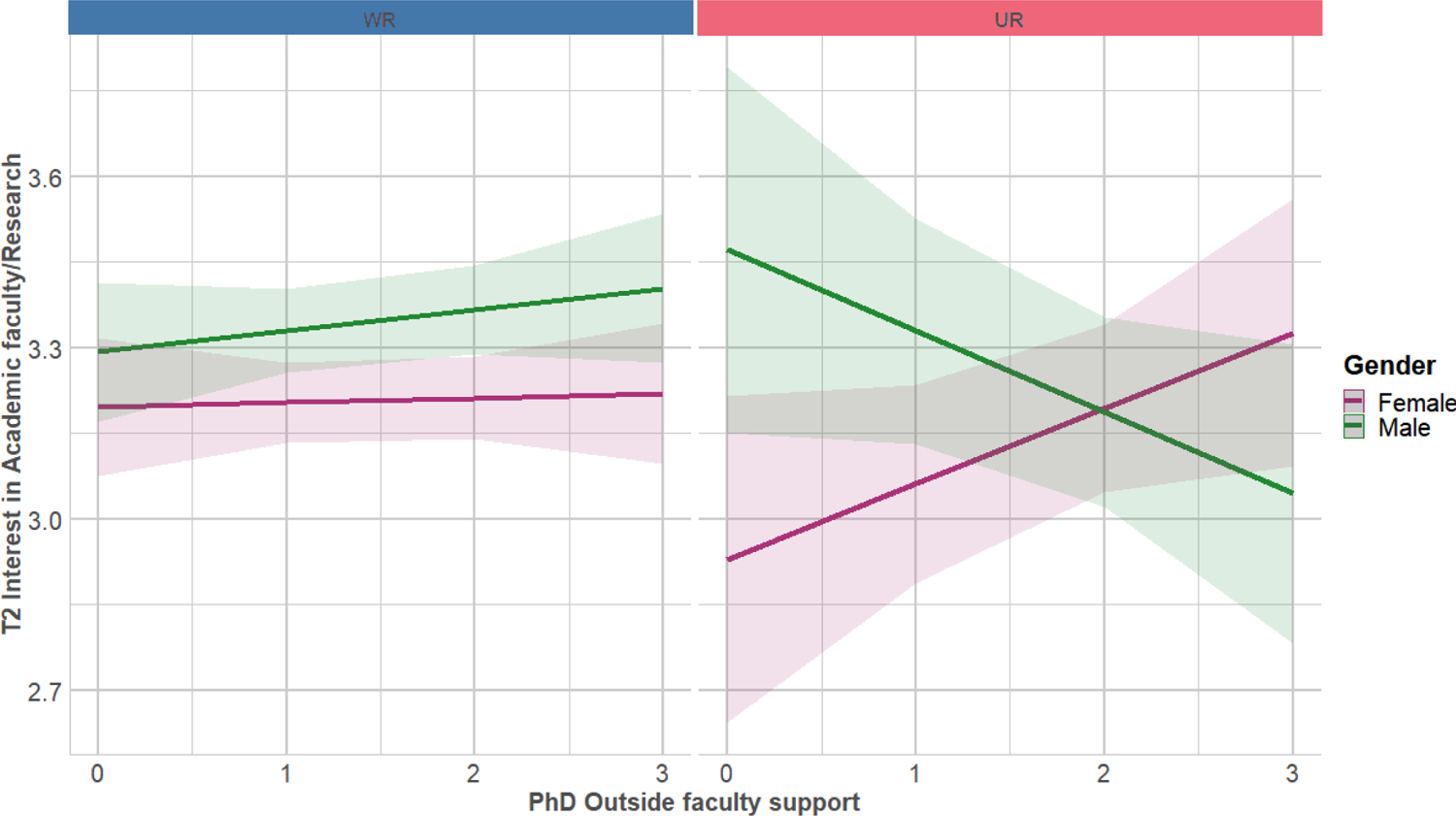
Outside faculty support during PhD associated with increased interest in research-focused academic faculty positions for UR females, but decreased interest in UR males, no gender difference in WR respondents. Regression lines predicting interest at the end of PhD training (T2) in research-focused academic faculty positions from level of outside faculty support during PhD training, split by gender and UR status.

Dependent variable was level of interest at the end of PhD training on a 4-point scale (where 1 represents “no interest” and 4 represents “strong interest”). Independent variable was level of helpfulness of outside faculty (0-3, 3 being very helpful). Interaction was significant at p < 0.05 (Table 1; Table S10a).

### Factors that Predict Changes in Interest in Teaching-focused Academic Positions

Second, we predicted change in respondents’ ratings of interest in teaching-focused academic faculty positions. The full regression equation was significant, accounting for 43.2% of the variance in T2 interest (Figure 12; Adjusted R^2^, Table 1). Interest in academic teaching at the start of graduate school predicted interest in academic teaching at the end of graduate school to a high degree (42.1% of the variance). Again, although other single variables were significant predictors, their effect sizes did not meet the threshold for reporting.

**Figure 12.**
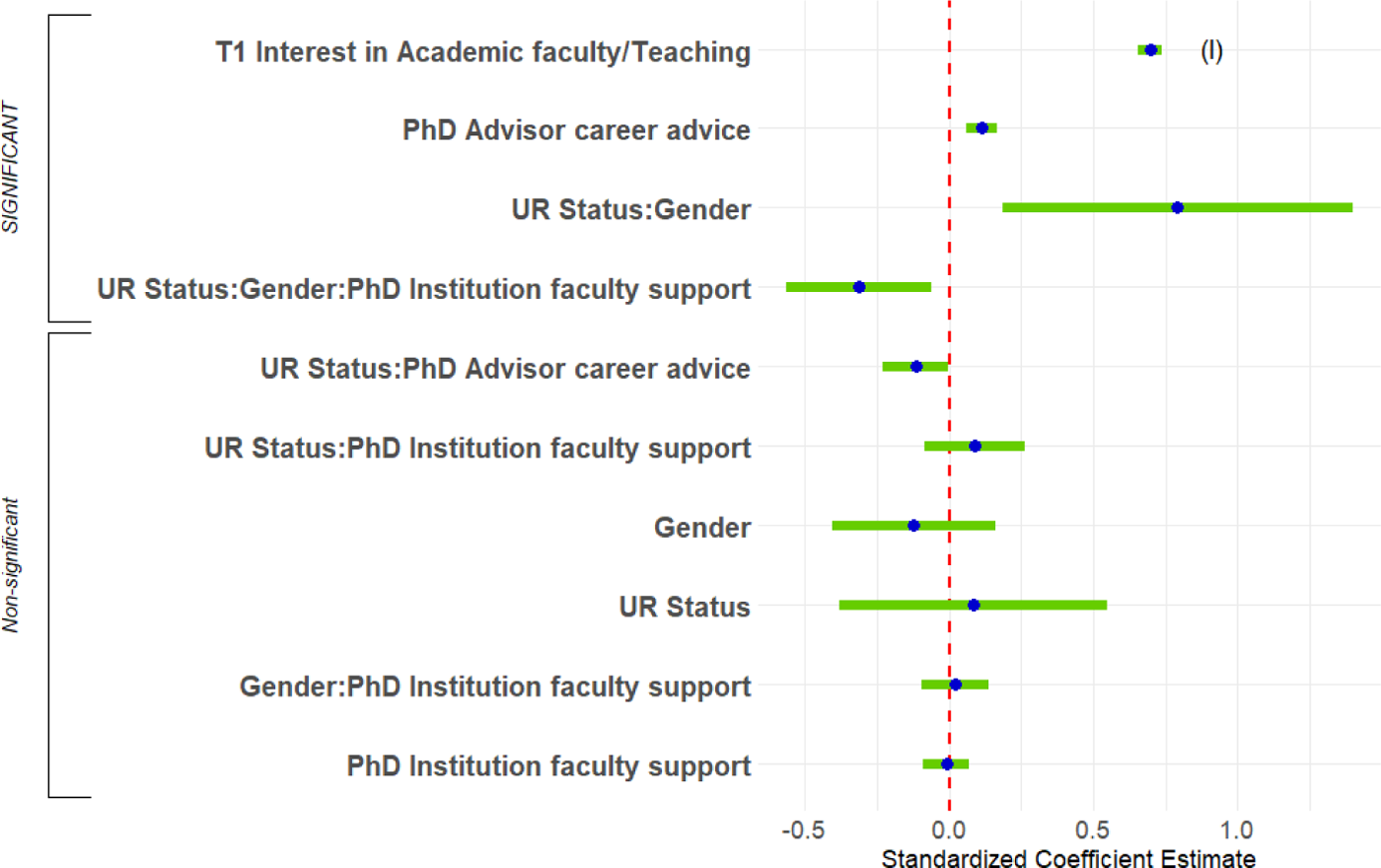
Predicting T2 interest in teaching-focused academic faculty positions. Standardized regression coefficients and error bars for linear regression predicting interest at the end of PhD training (T2) in teaching-focused academic faculty positions. Dependent variables were level of interest at the end of PhD training on a 4-point scale (where 1 represents “no interest” and 4 represents “strong interest”). Independent variables captured level of interest at the start of their PhD training (T1), experiences during PhD training, personal characteristics, objective measures, and interactions with gender and UR status. The entire equation was significant at p < 0.001 and captured 43.2% of the variance (adjusted; Table 1). Effect sizes are labeled when they reach at least “small” size. (l) = large effect size

**Figure 13.**
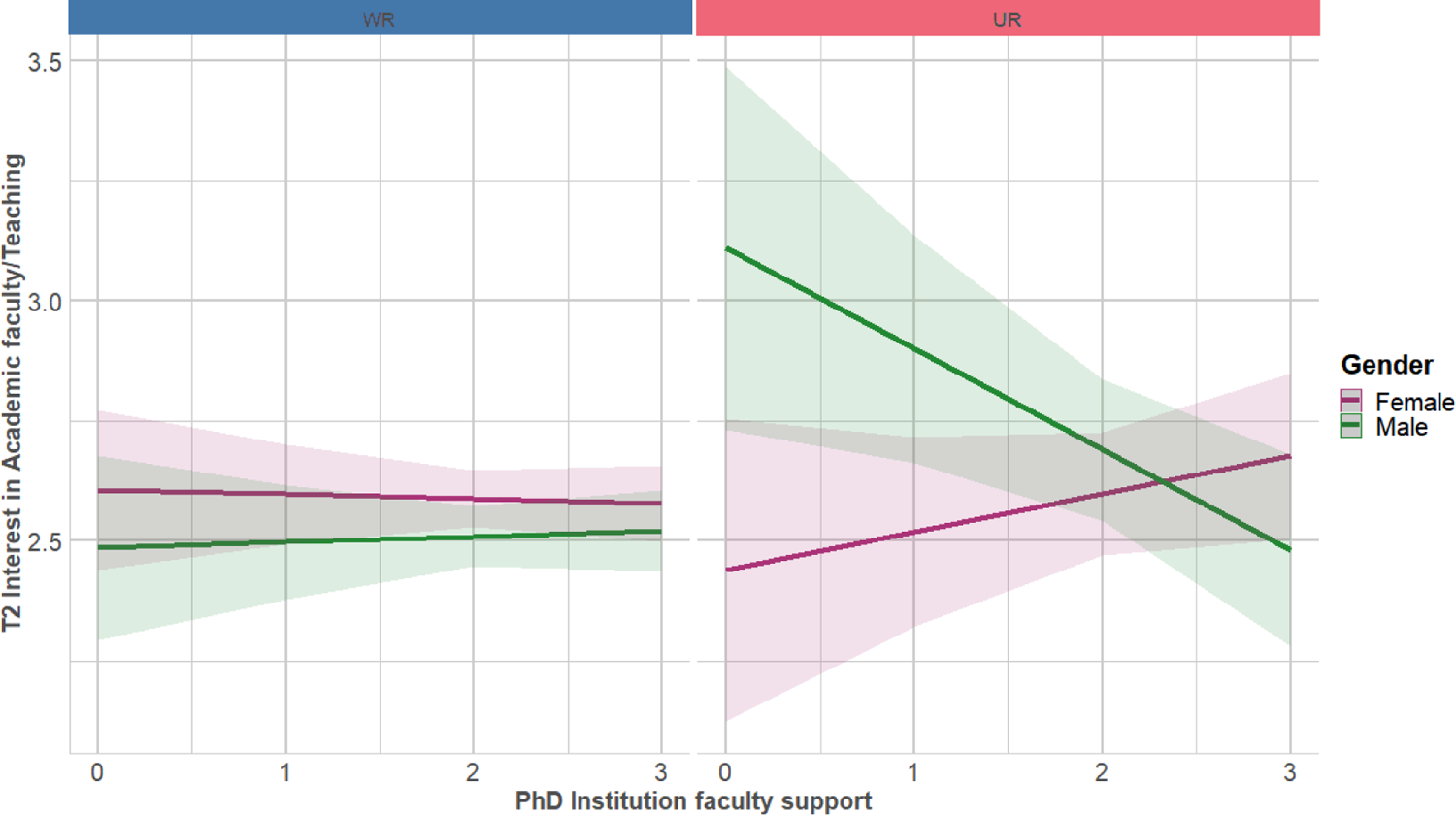
PhD institution faculty support was associated with increased interest in teaching-focused academic faculty positions for UR females, but was associated with decreased interest in UR males, no gender difference in WR respondents. Regression lines predicting interest at the end of PhD training (T2) in teaching-focused academic faculty positions from level of institution faculty support during PhD training, split by gender and UR status. Dependent variable was level of interest at the end of their PhD training (T2) on a 4-point scale (where 1 represents “no interest” and 4 represents “strong interest”). Independent variable was level of helpfulness of PhD institution faculty (0-3, 3 being very helpful). Interaction was significant at p < 0.05 (Table 1; Table S10a).

**Figure 14.**
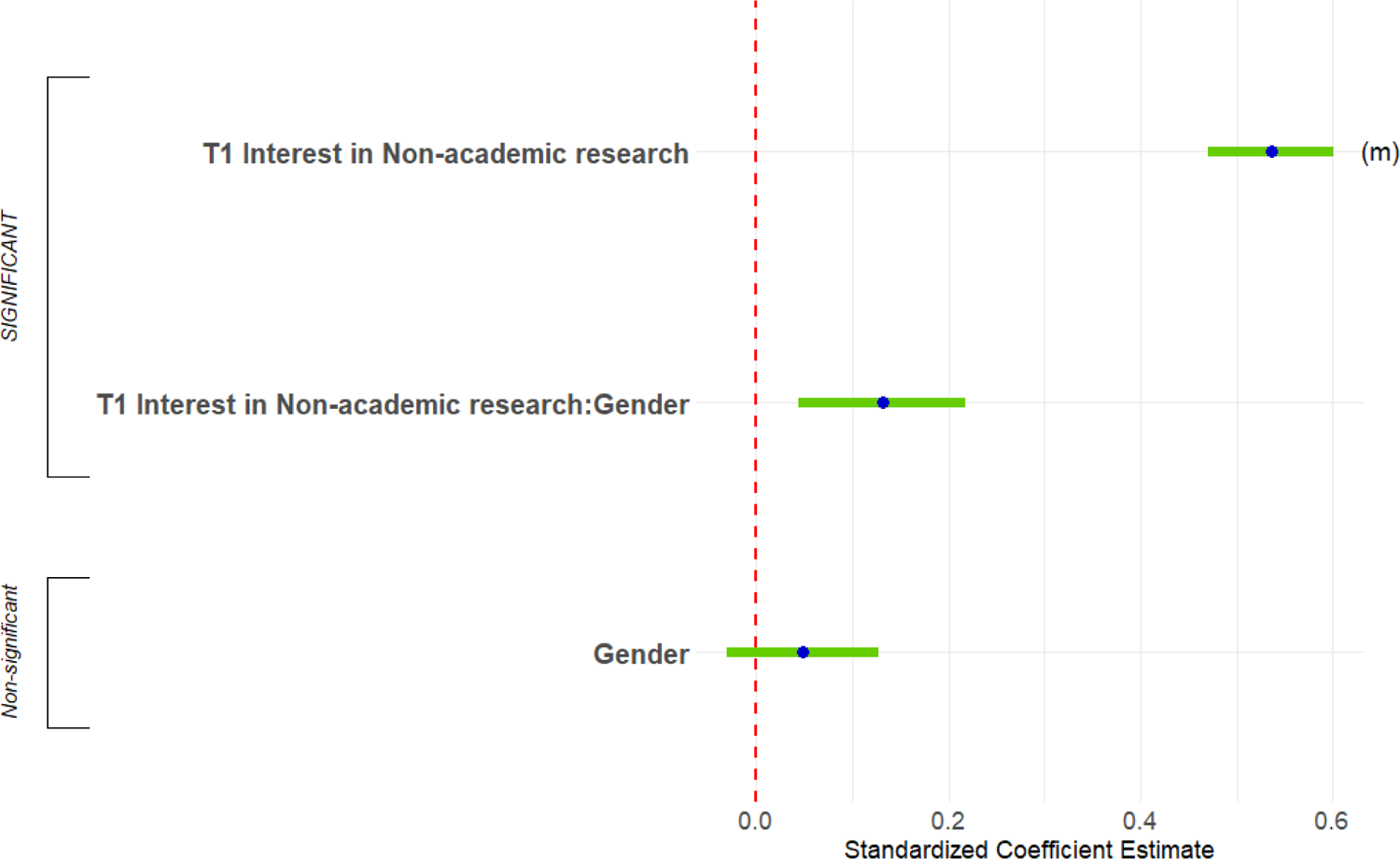
Predicting T2 interest in non-academic research positions. Standardized regression coefficients and error bars for linear regression predicting interest at the end of PhD training (T2) in research/non-academic positions. Dependent variable was level of interest at the end of their PhD training (T2) on a 4-point scale (where 1 represents “no interest” and 4 represents “strong interest”). Independent variables captured level of interest at the start of their PhD training (T1), experiences during PhD training, personal characteristics, objective measures, and interactions with gender and UR status. The entire equation was significant at p < 0.001 and captured 33.7% of the variance (adjusted; Table 1). Effect sizes are labeled when they reach at least “small” size. (m) = medium effect size

A 3-way interaction very similar to the interaction discussed in the last section, among gender, UR status, and level of support from faculty at respondents’ institutions during their PhD program (hereafter, “PhD institution faculty support”) was significant. Although there was no relation for WRs, PhD institution faculty support was associated with increased interest in teaching-focused academic faculty positions for UR females, but greatly decreased interest in UR males (Figure 13).<colcnt=1>

### Factors that Predict Changes in Interest in Non-academic Research Positions

Third, we predicted change in respondents’ ratings of interest in non-academic research positions. The full regression equation was significant, accounting for 33.7% of the variance (Figure 14; Adjusted R^2^, Table 1). Interest in non-academic research positions at the start of graduate school predicted interest in non-academic research positions at the end of graduate school to a moderate degree (14.8% of the variance). This finding was in the context, however, of a significant 2-way interaction between gender and T1 rating of interest in non-academic research positions. The association between interest at start of graduate school and interest at end of graduate school was somewhat stronger for males than it was for females (Figure 15).

**Figure 15.**
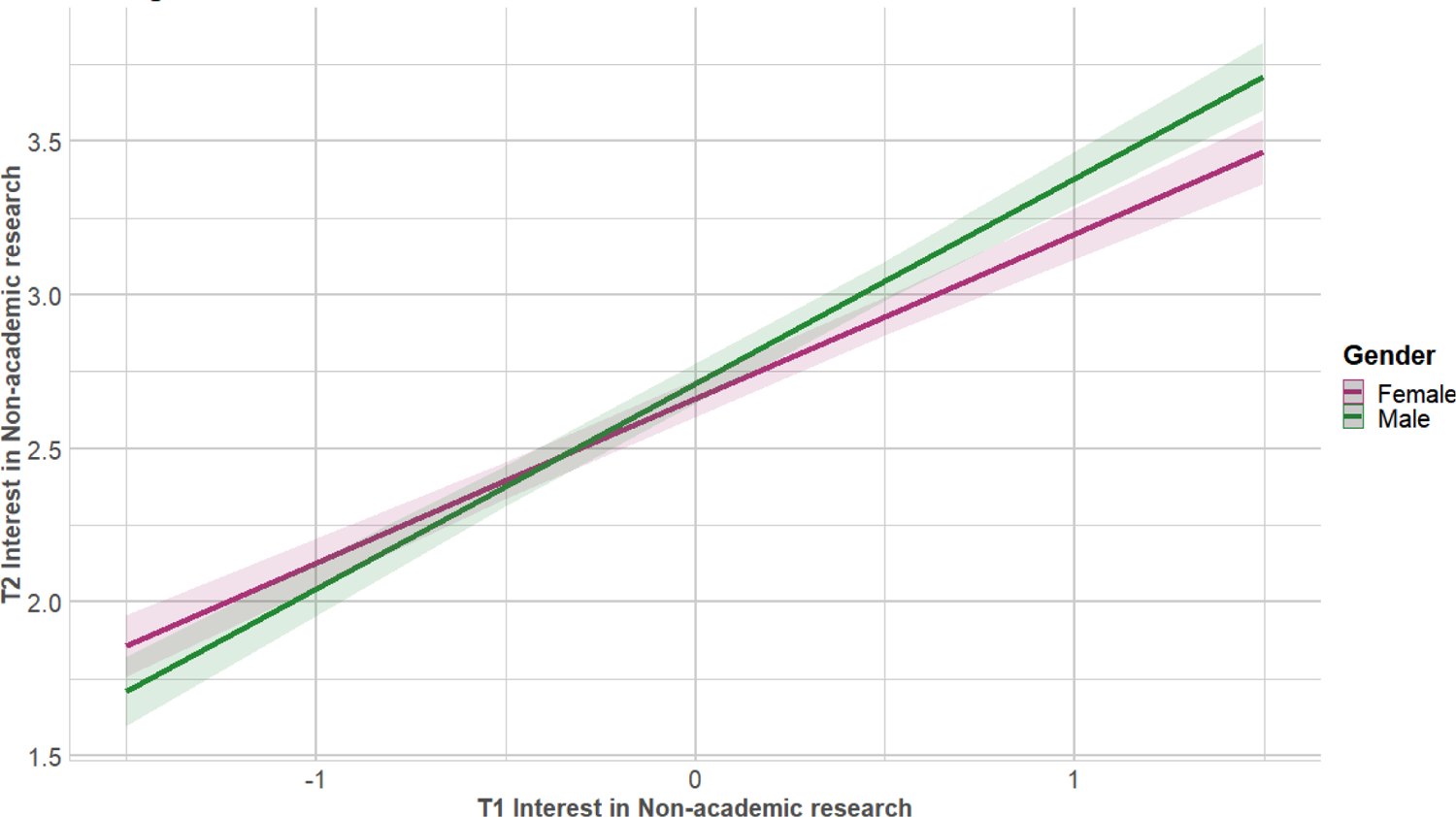
T1 interest in non-academic research positions was a stronger predictor for T2 interest for males than females. Graph shows regression lines predicting interest at the end of PhD training (T2) in research/non-academic positions from T1 interest, split by gender. Dependent variable from 1,479 PhD neuroscientists who were asked to rate their level of interest at the end of their PhD training (T2) on a 4-point scale (where 1 represents “no interest” and 4 represents “strong interest”). Independent variable was interest at T1, centered for interaction (−2 to 2). Interaction was significant at p < 0.01 (Table 1; Table S10a).

### Factors that Predict Changes in Interest in Science-related, Non-research Positions

Finally, we predicted change in respondents’ ratings of interest in science-related, non-research positions. The full regression equation was significant, accounting for 41.7% of the variance (Figure 16; Adjusted R^2^, Table 1). Interest in science-related, non-research positions at the start of graduate school predicted interest in science/non-research at the end of graduate school to a high degree (32.3% of the variance; this finding was in the context, however, of a significant 2-way interaction, below). Helpfulness of career advice from PhD advisor was also significant. This finding was in the context, however, of a significant 2-way interaction with a moderate effect size. Lower ratings of the helpfulness of career advice from PhD advisors was related to becoming more interested in science-related, non-research positions across graduate school (1.4% of the variance).

**Figure 16.**
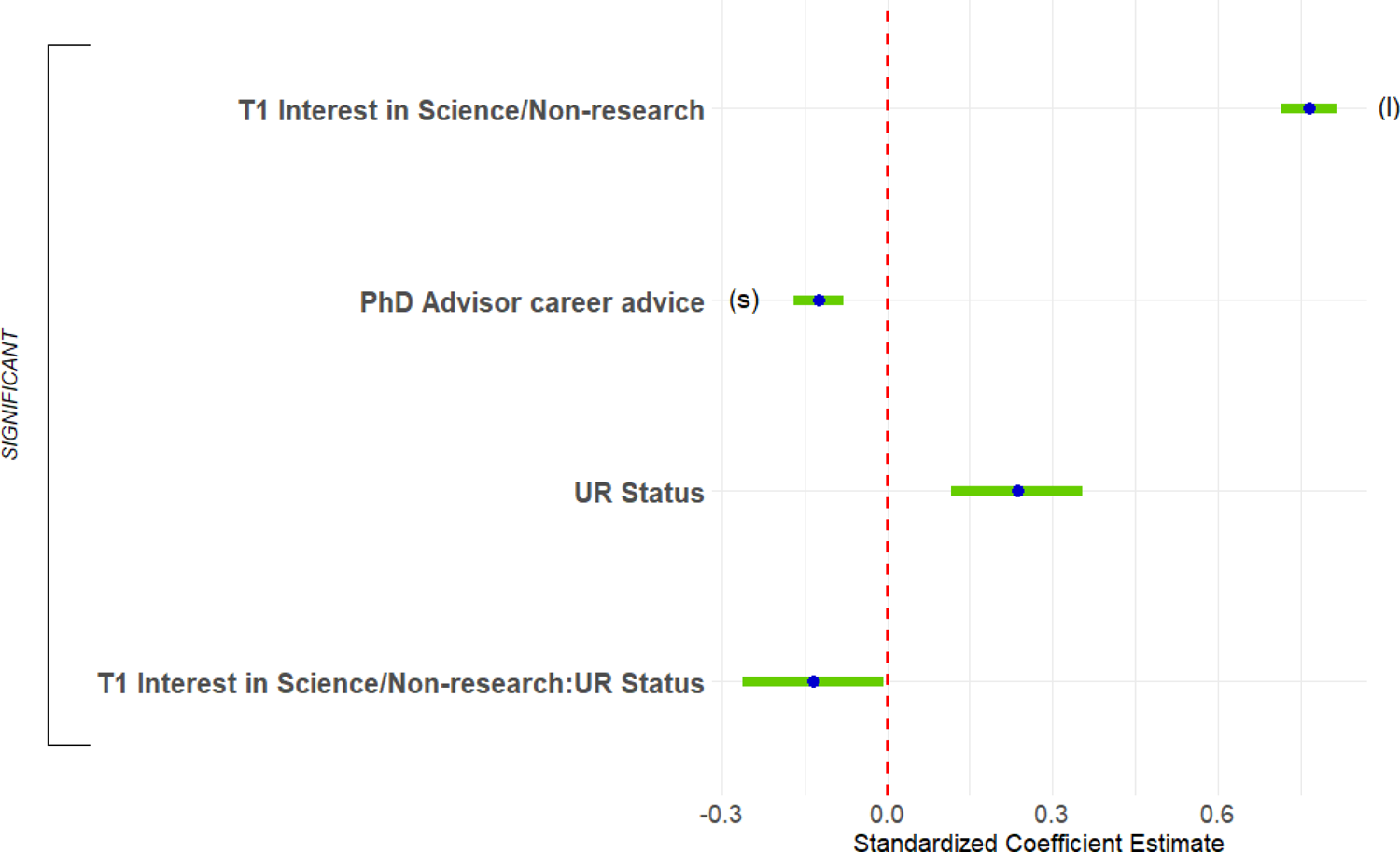
Predicting T2 interest in science-related, non-research positions. Standardized regression coefficients and error bars for linear regression predicting interest at the end of PhD training (T2) in science-related, non-research positions. Dependent variable was level of interest at the end of their PhD training (T2) on a 4-point scale (where 1 represents “no interest” and 4 represents “strong interest”). Independent variables captured level of interest at the start of their PhD training (T1), experiences during PhD training, personal characteristics, objective measures, and interactions with gender and UR status. The entire equation was significant at p < 0.001 and captured 41.7% of the variance (adjusted; Table 1). Effect sizes are labeled when they reach at least “small” size. (l) = large effect size

A single 2-way interaction, between UR status and T1 rating of interest in science-related, non-research positions, was significant for this equation. The association between T1 interest and T2 interest was somewhat stronger for WRs than it was for URs (Figure 17).

**Figure 17.**
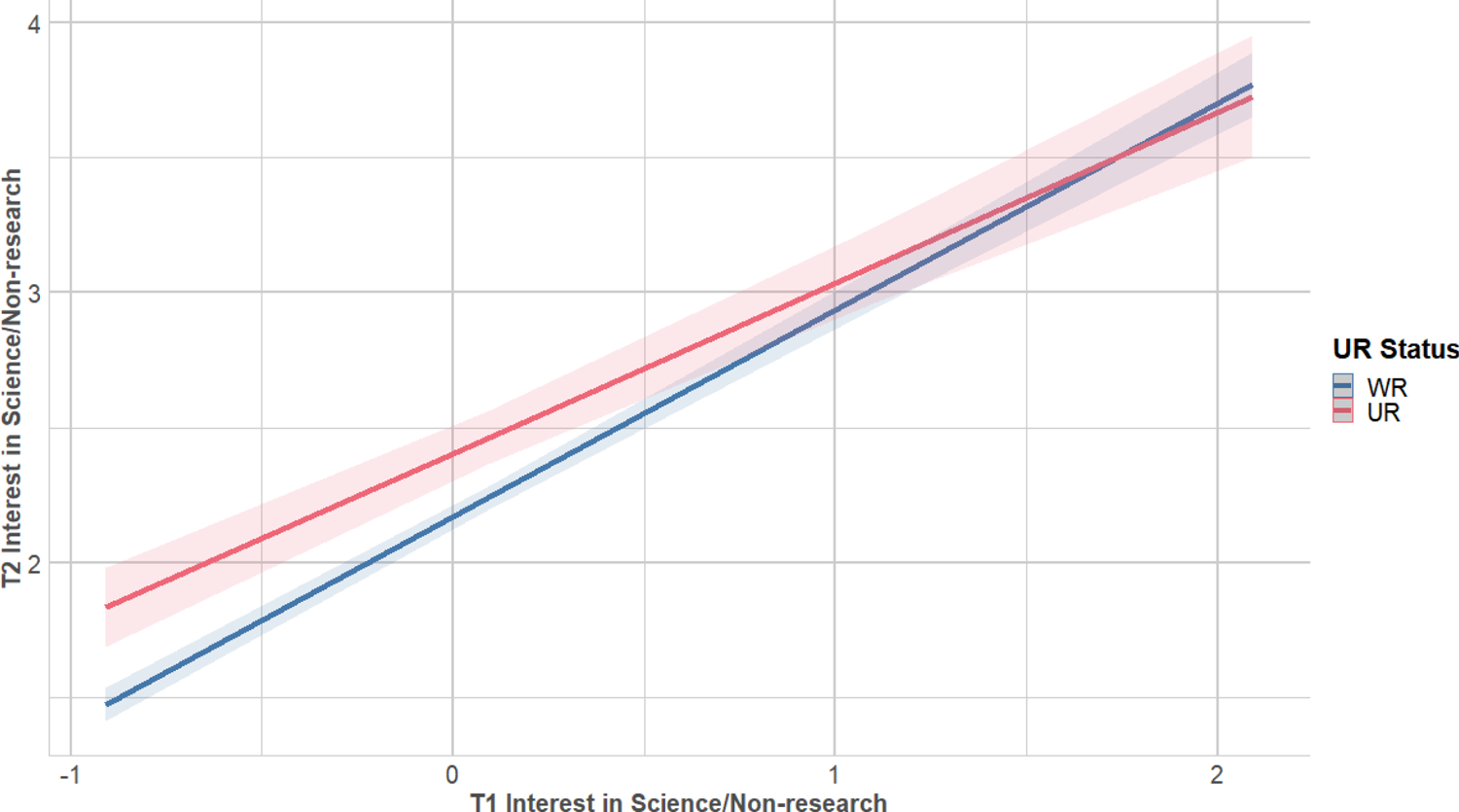
T1 interest in science-related, non-research positions was a stronger predictor for T2 interest for WR than UR respondents. Regression lines predicting interest at the end of PhD training (T2) in science-related, non-research positions from interest at the start of PhD training (T1), split by UR status. Dependent variable was level of interest at the end of PhD training on a 4-point scale (where 1 represents “no interest” and 4 represents “strong interest”). Independent variable was interest at T1, centered for interaction (−2 to 2). Interaction was significant at p < 0.05 (Table 1; Table S10a).

### Predicting Current Interest

We were also interested in which factors predicted current interest in different careers. For this analysis we chose predictor variables that captured issues that were either current, or indicative of participants’ time in postdocs. Descriptions of the procedures can be found in the Methods section, and preliminary steps and results for construction of the regression are reported in Tables S8 and S9 in the *Supporting Information Appendix*.

The final step in this set of analyses was to conduct regression analyses to predict current interest in the different careers. Each equation was constructed by regressing one of the four career interest ratings at T3 (current) on: 1) the variables that had significant correlations with it in the first step (Table S8), and 2) the interactions that were significant predictors of it in the second step (Table S9). The results of these analyses are presented in Table 2. We discuss the regression for each career type in the following sub-sections.

**Table 2.**
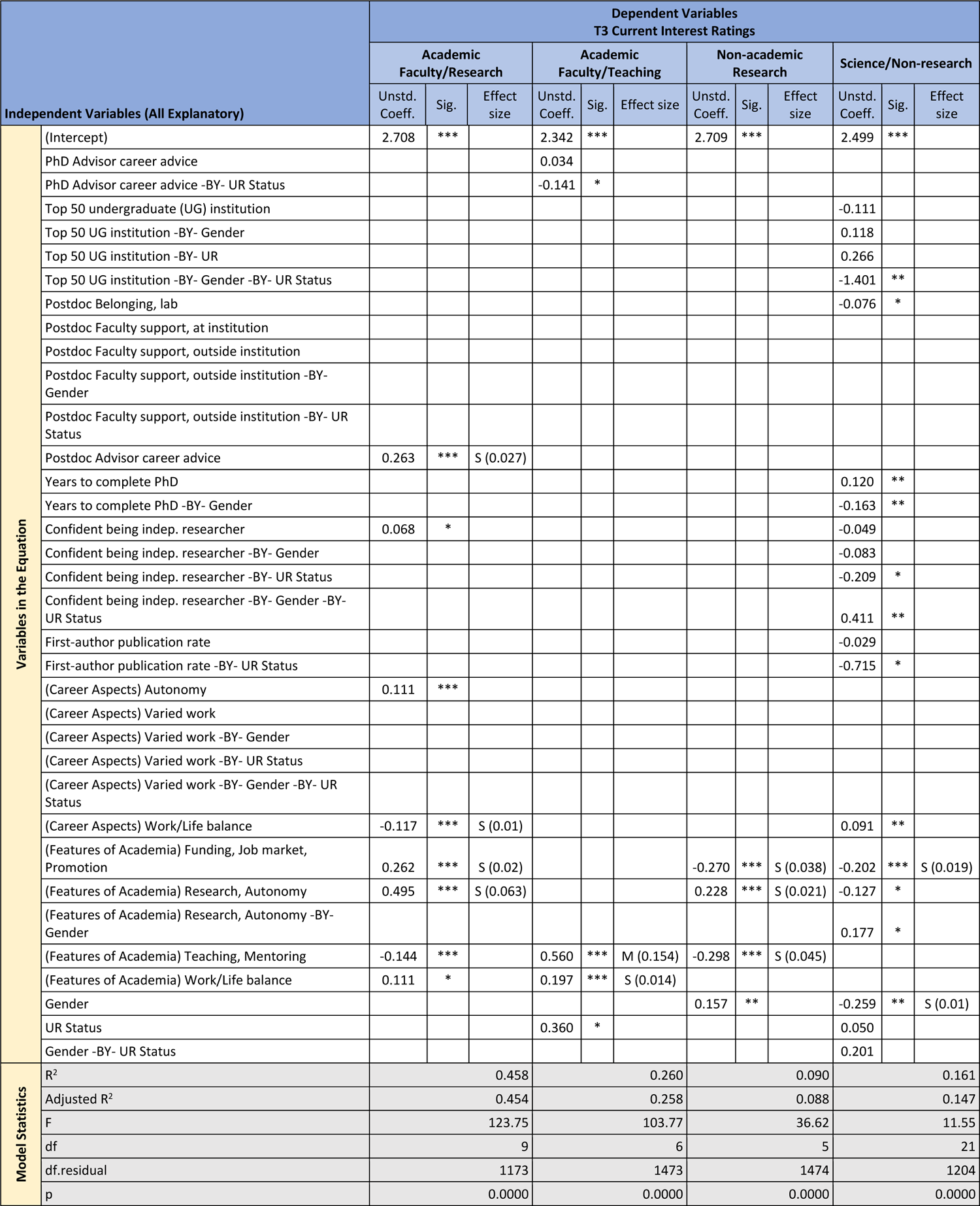
Regressions Predicting Career Interest Ratings at T3 (Current) from All Explanatory Variables. Results of regressions for each of the four career interest T3 ratings (current) on significantly correlated variables (Table S8) and significant predictor interactions (Table S9). Unstd. Coeff. = Unstandardized coefficient; sig. = significance. Effect sizes are labeled when they reach at least “small” size. (S) = small effect size, (M) = medium effect size. * = p < 0.05; ** = p < 0.01; *** = p < 0.001

| Independent Variables (All Explanatory) |  | Dependent Variables<br>T3 Current Interest Ratings |  |  |  |  |  |  |  |  |  |  |  |
| --- | --- | --- | --- | --- | --- | --- | --- | --- | --- | --- | --- | --- | --- |
|  |  | Academic<br>Faculty/Research |  |  | Academic<br>Faculty/Teaching |  |  | Non-academic<br>Research |  |  | Science/Non-research |  |  |
|  |  | Unstd.<br>Coeff. | Sig. | Effect<br>size | Unstd.<br>Coeff. | Sig. | Effect size | Unstd.<br>Coeff. | Sig. | Effect<br>size | Unstd.<br>Coeff. | Sig. | Effect<br>size |
| Variables in the Equation | (Intercept) | 2.708 | *** |  | 2.342 | *** |  | 2.709 | *** |  | 2.499 | *** |  |
|  | PhD Advisor career advice |  |  |  | 0.034 |  |  |  |  |  |  |  |  |
|  | PhD Advisor career advice -BY- UR Status |  |  |  | -0.141 | * |  |  |  |  |  |  |  |
|  | Top 50 undergraduate (UG) institution |  |  |  |  |  |  |  |  |  | -0.111 |  |  |
|  | Top 50 UG institution -BY- Gender |  |  |  |  |  |  |  |  |  | 0.118 |  |  |
|  | Top 50 UG institution -BY- UR |  |  |  |  |  |  |  |  |  | 0.266 |  |  |
|  | Top 50 UG institution -BY- Gender -BY- UR Status |  |  |  |  |  |  |  |  |  | -1.401 | ** |  |
|  | Postdoc Belonging, lab |  |  |  |  |  |  |  |  |  | -0.076 | * |  |
|  | Postdoc Faculty support, at institution |  |  |  |  |  |  |  |  |  |  |  |  |
|  | Postdoc Faculty support, outside institution |  |  |  |  |  |  |  |  |  |  |  |  |
|  | Postdoc Faculty support, outside institution -BY- Gender |  |  |  |  |  |  |  |  |  |  |  |  |
|  | Postdoc Faculty support, outside institution -BY- UR Status |  |  |  |  |  |  |  |  |  |  |  |  |
|  | Postdoc Advisor career advice | 0.263 | *** | S (0.027) |  |  |  |  |  |  |  |  |  |
|  | Years to complete PhD |  |  |  |  |  |  |  |  |  | 0.120 | ** |  |
|  | Years to complete PhD -BY- Gender |  |  |  |  |  |  |  |  |  | -0.163 | ** |  |
|  | Confident being indep. researcher | 0.068 | * |  |  |  |  |  |  |  | -0.049 |  |  |
|  | Confident being indep. researcher -BY- Gender |  |  |  |  |  |  |  |  |  | -0.083 |  |  |
|  | Confident being indep. researcher -BY- UR Status |  |  |  |  |  |  |  |  |  | -0.209 | * |  |
|  | Confident being indep. researcher -BY- Gender -BY- UR Status |  |  |  |  |  |  |  |  |  | 0.411 | ** |  |
|  | First-author publication rate |  |  |  |  |  |  |  |  |  | -0.029 |  |  |
|  | First-author publication rate -BY- UR Status |  |  |  |  |  |  |  |  |  | -0.715 | * |  |
|  | (Career Aspects) Autonomy | 0.111 | *** |  |  |  |  |  |  |  |  |  |  |
|  | (Career Aspects) Varied work |  |  |  |  |  |  |  |  |  |  |  |  |
|  | (Career Aspects) Varied work -BY- Gender |  |  |  |  |  |  |  |  |  |  |  |  |
|  | (Career Aspects) Varied work -BY- UR Status |  |  |  |  |  |  |  |  |  |  |  |  |
|  | (Career Aspects) Varied work -BY- Gender -BY- UR Status |  |  |  |  |  |  |  |  |  |  |  |  |
|  | (Career Aspects) Work/Life balance | -0.117 | *** | S (0.01) |  |  |  |  |  |  | 0.091 | ** |  |
|  | (Features of Academia) Funding, Job market, Promotion | 0.262 | *** | S (0.02) |  |  |  | -0.270 | *** | S (0.038) | -0.202 | *** | S (0.019) |
|  | (Features of Academia) Research, Autonomy | 0.495 | *** | S (0.063) |  |  |  | 0.228 | *** | S (0.021) | -0.127 | * |  |
|  | (Features of Academia) Research, Autonomy -BY- Gender |  |  |  |  |  |  |  |  |  | 0.177 | * |  |
|  | (Features of Academia) Teaching, Mentoring | -0.144 | *** |  | 0.560 | *** | M (0.154) | -0.298 | *** | S (0.045) |  |  |  |
|  | (Features of Academia) Work/Life balance | 0.111 | * |  | 0.197 | *** | S (0.014) |  |  |  |  |  |  |
|  | Gender |  |  |  |  |  |  | 0.157 | ** |  | -0.259 | ** | S (0.01) |
|  | UR Status |  |  |  | 0.360 | * |  |  |  |  | 0.050 |  |  |
|  | Gender -BY- UR Status |  |  |  |  |  |  |  |  |  | 0.201 |  |  |
| Model Statistics | R<sup>2</sup> | 0.458 |  |  | 0.260 |  |  | 0.090 |  |  | 0.161 |  |  |
|  | Adjusted R<sup>2</sup> | 0.454 |  |  | 0.258 |  |  | 0.088 |  |  | 0.147 |  |  |
|  | F | 123.75 |  |  | 103.77 |  |  | 36.62 |  |  | 11.55 |  |  |
|  | df | 9 |  |  | 6 |  |  | 5 |  |  | 21 |  |  |
|  | df.residual | 1173 |  |  | 1473 |  |  | 1474 |  |  | 1204 |  |  |
|  | p | 0.0000 |  |  | 0.0000 |  |  | 0.0000 |  |  | 0.0000 |  |  |

### Predictors of Current Interest in Research-focused Academic Positions

First, we predicted respondents’ ratings of current interest in research-focused academic faculty positions. The full regression equation was significant, accounting for 45.4% of the variance (Figure 18; Adjusted R^2^, Table 2). Four predictors were significant and had sufficient effect size to report. First, more helpful career advice from their PhD advisors was associated with greater interest in academic research positions (2.7% of the variance). Second, respondents for whom work/life balance was less important were more interested in academic research positions (1.0% of the variance). Third, respondents who were more positive about the funding/job market/promotion features of academia were more interested in academic research positions (2.0% of the variance). Finally, respondents who were more positive about the research/autonomy features of academia were more interested in academic research positions (6.3% of the variance). Although other single variables were significant predictors, their effect sizes did not meet the threshold for reporting. In addition, there were no significant interactions for this equation.

**Figure 18.**
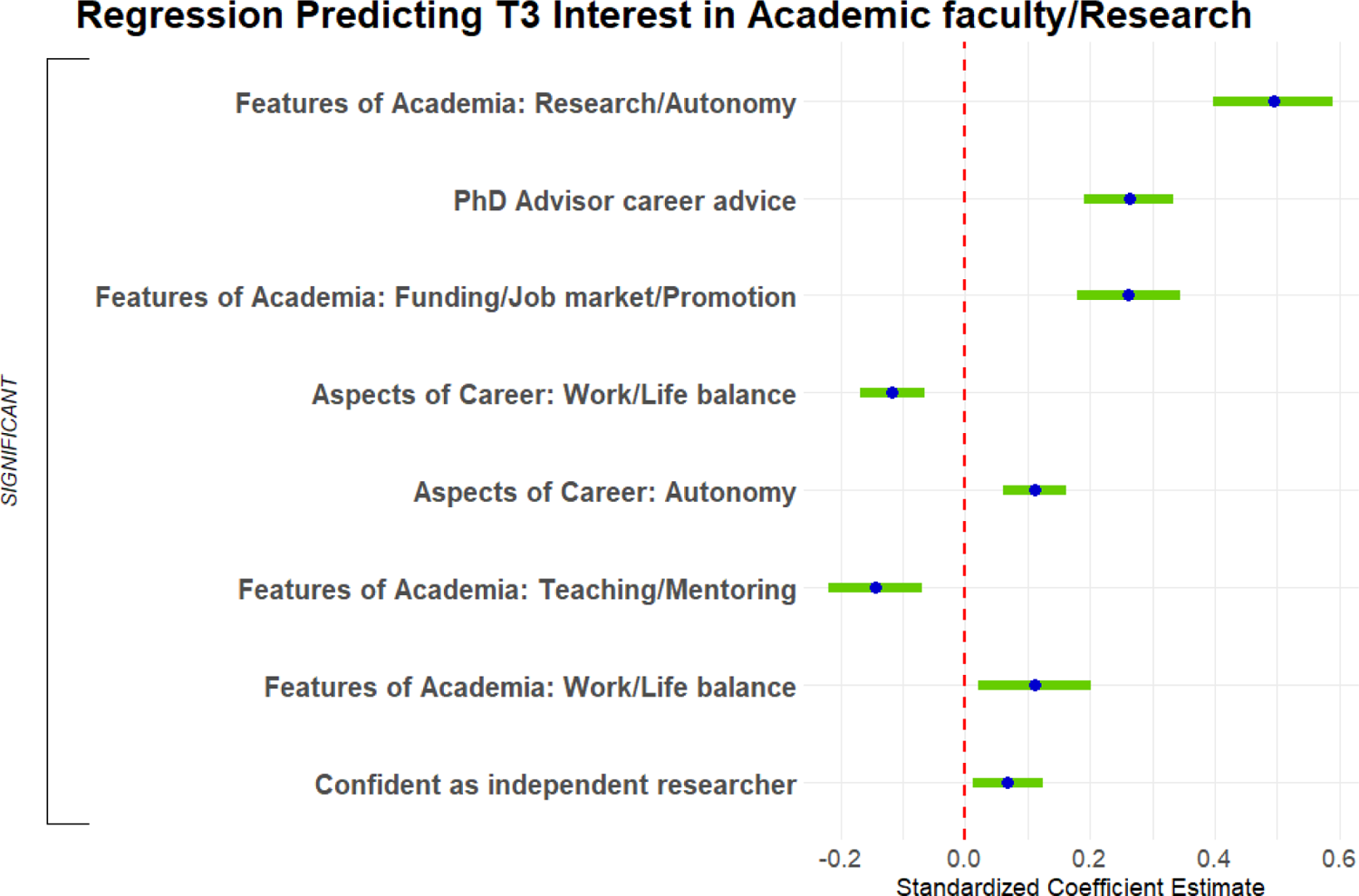
Predicting T3 interest in research-focused academic faculty positions. Standardized regression coefficients and error bars for linear regression predicting current interest (T3) in research-focused academic faculty positions. Dependent variable was current level of interest on a 4-point scale (where 1 represents “no interest” and 4 represents “strong interest”). Independent variables captured experiences during PhD training and postdocs, personal characteristics, objective measures, and interactions with gender and UR status. The entire equation was significant at p < 0.001 and captured 45.4% of the variance (adjusted; Table 2). Effect sizes are labeled when they reach at least “small” size. (s) = small effect size

### Predictors of Current Interest in Teaching-focused Academic Positions

Second, we predicted respondents’ ratings of current interest in teaching-focused academic faculty positions. The full regression equation was significant, accounting for 25.8% of the variance (Figure 19; Adjusted R^2^, Table 2). Two predictors were significant and had sufficient effect size to report. First, perhaps unsurprisingly, respondents who were more positive about the teaching/mentoring features of academia were more interested in academic teaching (15.4% of the variance). Second, respondents for whom work/life balance was important were more interested in academic teaching (1.4% of the variance).

**Figure 19.**
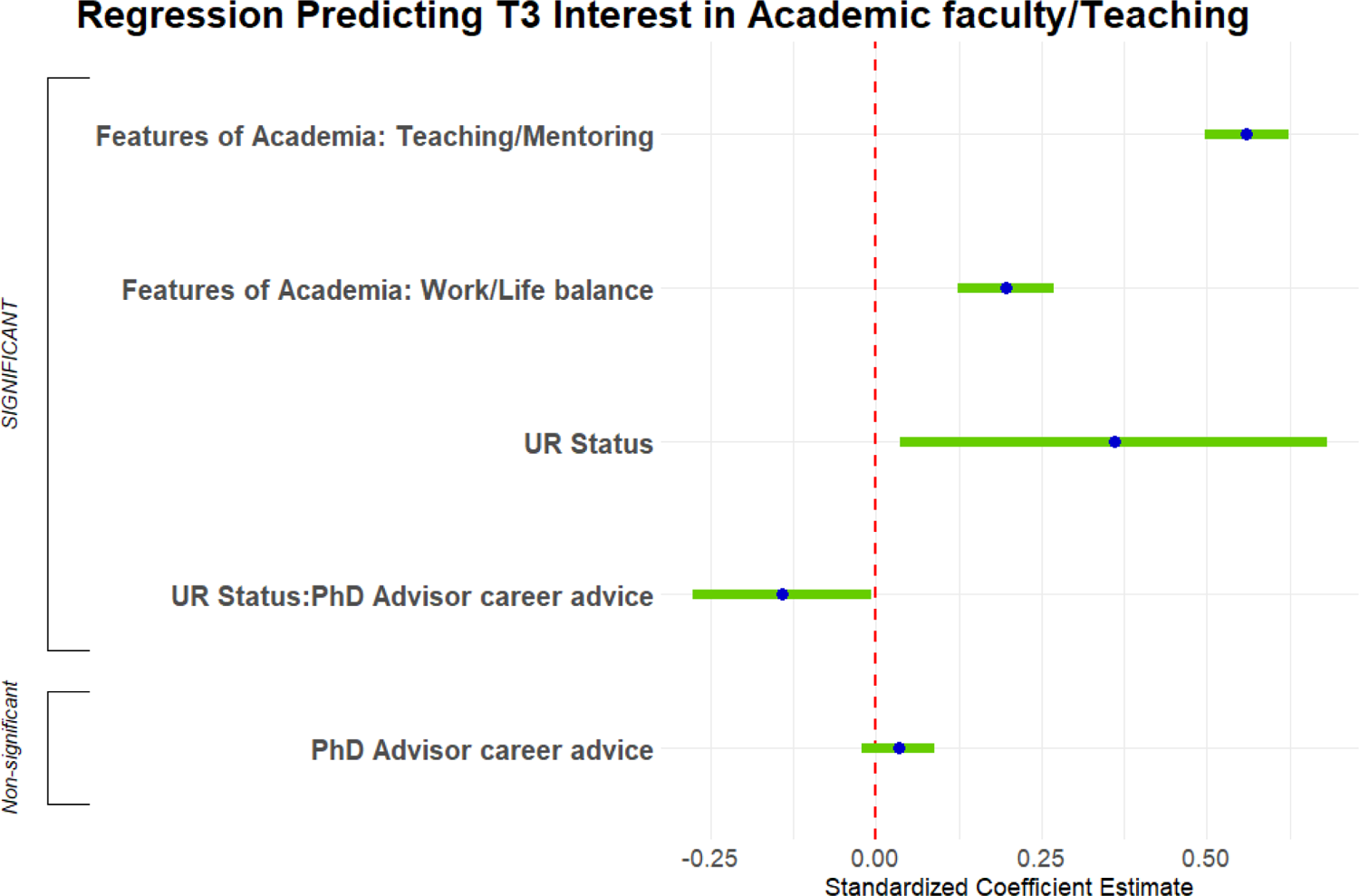
Predicting T3 interest in teaching-focused academic faculty positions. Graph shows standardized regression coefficients and error bars for linear regression predicting current interest (T3) in teaching-focused academic faculty positions. Dependent variable was current level of interest on a 4-point scale (where 1 represents “no interest” and 4 represents “strong interest”). Independent variables captured their experiences during PhD training and postdocs, personal characteristics, objective measures, and interactions with gender and UR status. The entire equation was significant at p < 0.001 and captured 25.8% of the variance (adjusted; Table 2). Effect sizes are labeled when they reach at least “small” size. (s) = small effect size, (m) = medium effect size

A single 2-way interaction, between UR status and helpfulness of career advice from your PhD advisor, was significant. Helpful career advice from the PhD advisor was weakly associated with increased interest in teaching-focused academic faculty positions for WR respondents but was strongly associated with decreased interest in UR respondents (Figure 20).

**Figure 20.**
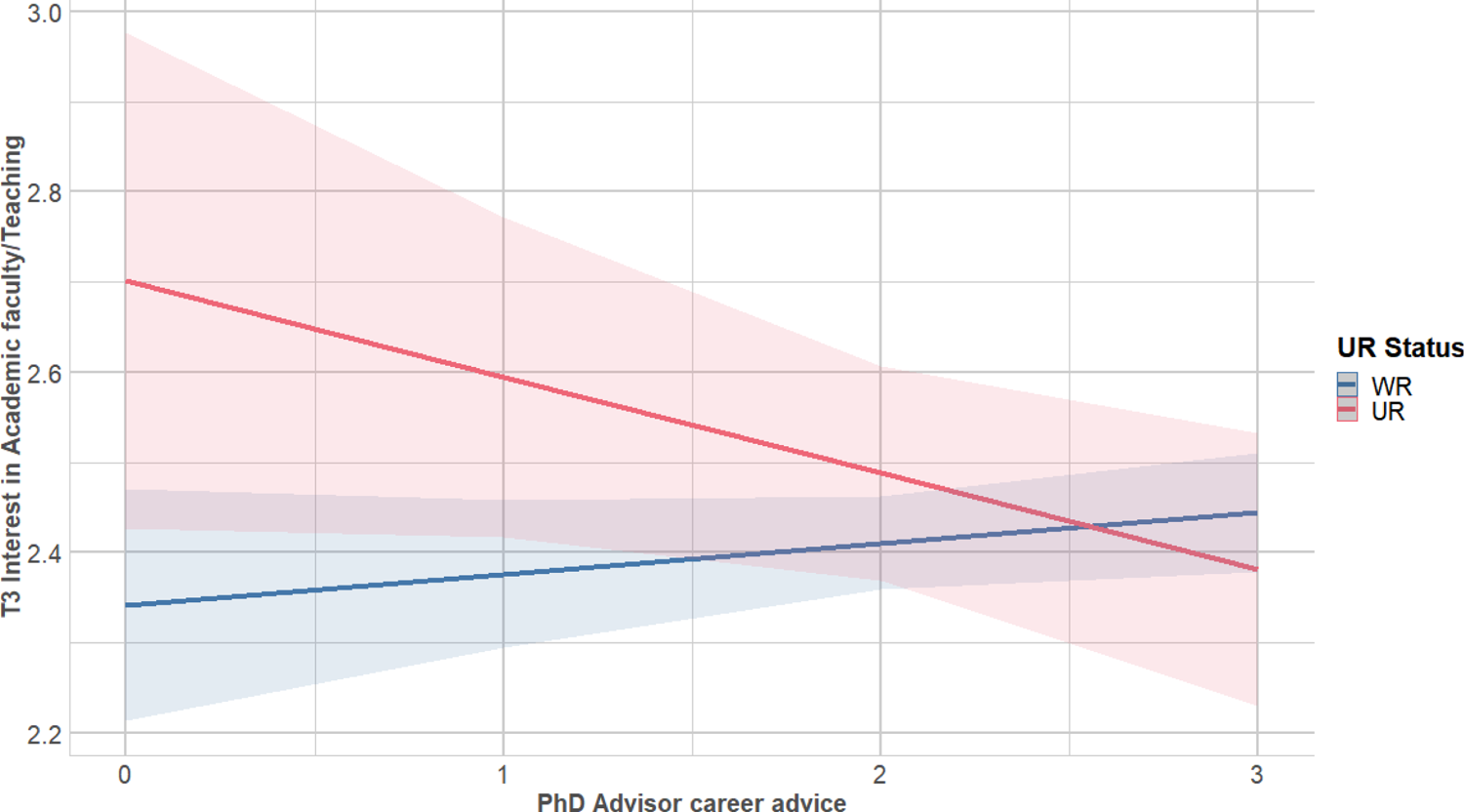
Helpful PhD advisor career advice was associated with increased interest in teaching-focused academic faculty positions for WR respondents but was associated with decreased interest in UR respondents. Regression lines predicting current (T3) interest in teaching-focused academic faculty positions from PhD advisor career advice, split by UR status. Dependent was current level of interest on a 4-point scale (where 1 represents “no interest” and 4 represents “strong interest”). Independent variable was helpfulness of career advice from PhD advisor (0-3, 3 being very helpful). Interaction was significant at p < 0.05 (Table 2; Table S10b).

### Predictors of Current Interest in Non-academic Research Positions

Third, we predicted respondents’ ratings of current interest in non-academic research positions. The full regression equation was significant but accounted for only 8.8% of the variance (Figure 21; Adjusted R^2^, Table 2). Three predictors were significant and had sufficient effect size to report. First, respondents who were more negative about the funding/job market/promotion features of academia were more interested in non-academic research positions (3.8% of the variance). Second, respondents who were more positive about the research/autonomy features of academia were more interested in non-academic research positions (2.1% of the variance). Finally, respondents who were more negative about the teaching/mentoring features of academia were more interested in non-academic research positions (4.5% of the variance). There were no significant interactions for this equation.

**Figure 21.**
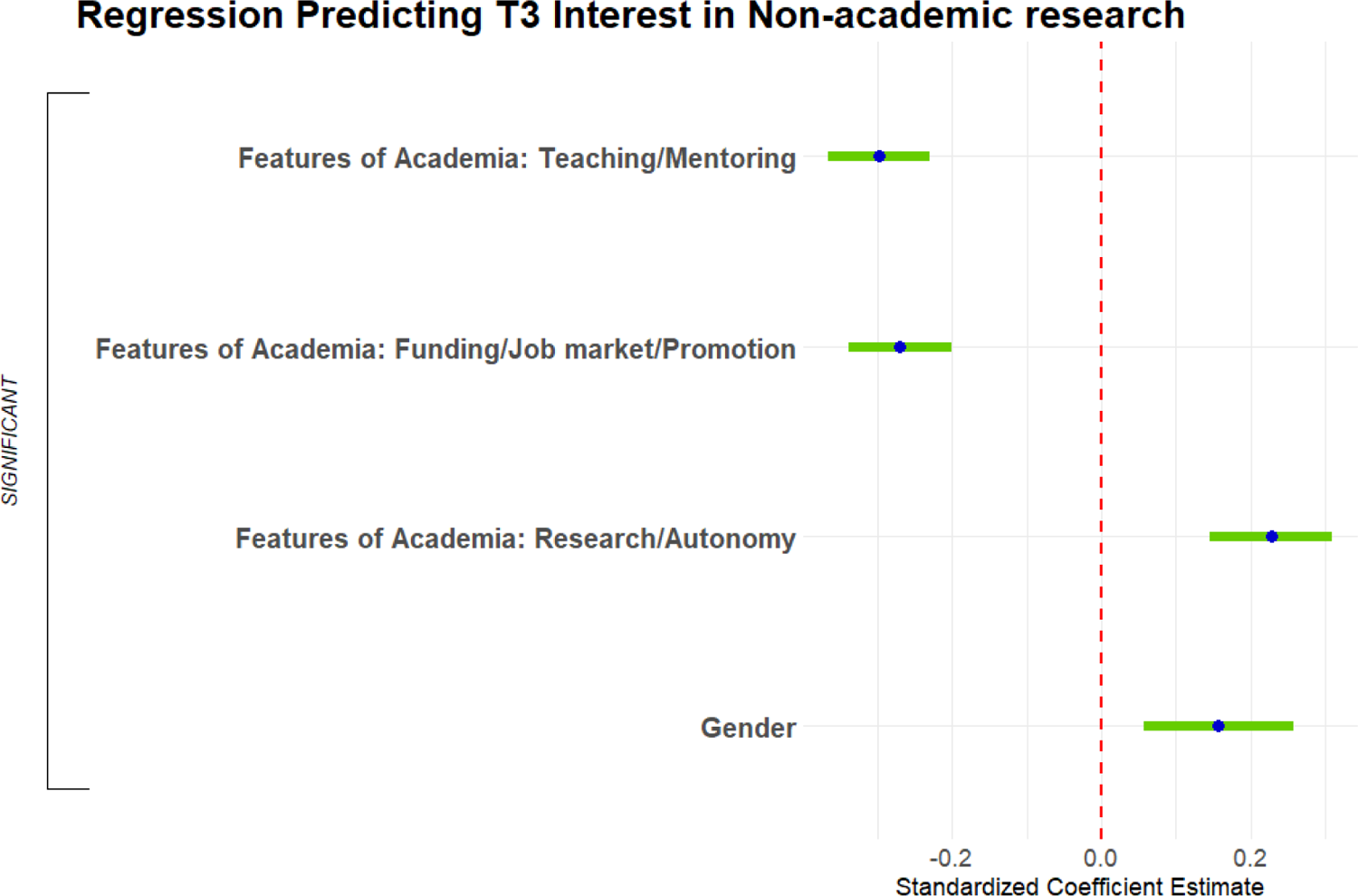
Predicting T3 interest in non-academic research positions. Standardized regression coefficients and error bars for linear regression predicting current interest (T3) in non-academic research positions. Dependent variable was current level of interest on a 4-point scale (where 1 represents “no interest” and 4 represents “strong interest”). Independent variables captured their experiences during PhD training and postdocs, personal characteristics, objective measures, and interactions with gender and UR status. The entire equation was significant at p < 0.001 and captured 8.8% of the variance (adjusted; Table 2). Effect sizes are labeled when they reach at least “small” size. (s) = small effect size

### Predictors of Current Interest in Science-related, Non-research Positions

Finally, we predicted respondents’ ratings of current interest in science-related, non-research positions. The full regression equation was significant, accounting for only 14.7% of the variance (Figure 22; Adjusted R^2^, Table 2). Two predictors were significant and had sufficient effect size to report. First, respondents who were more negative about the funding/job market/promotion features of academia were more interested in science-related, non-research positions (1.9% of the variance). Second, males were less interested in science-related, non-research positions (1.0% of the variance).

**Figure 22.**
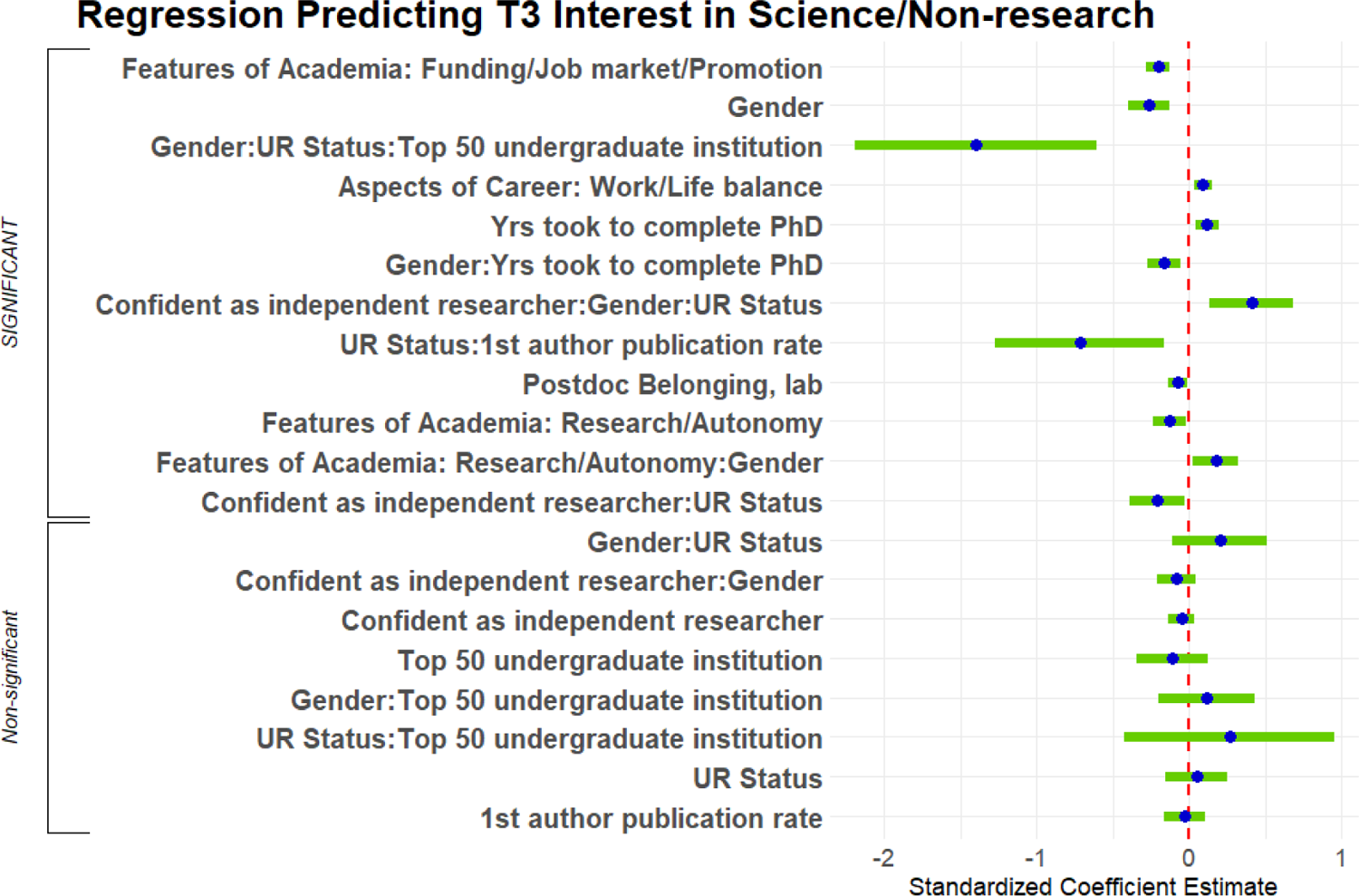
Predicting T3 interest in science-related, non-research positions. Standardized regression coefficients and error bars for linear regression predicting current interest (T3) in science-related, non-research positions. Dependent variable was current level of interest on a 4-point scale (where 1 represents “no interest” and 4 represents “strong interest”). Independent variables captured their experiences during PhD training and postdocs, personal characteristics, objective measures, and interactions with gender and UR status. The entire equation was significant at p < 0.001 and captured 14.7% of the variance (adjusted; Table 2). Effect sizes are labeled when they reach at least “small” size. (s) = small effect size<colcnt=1>

There were no fewer than 6 significant interactions for this equation, although one was a 2-way interaction within the context of a 3-way interaction and will not be reported. The first was a complex 3-way interaction among Gender, UR status, and whether respondents went to a top 50 undergraduate institution. Two groups of males were less interested in science-related, non-research positions than the females in their groups – WR males who were not at a top 50 undergraduate institution and UR males who were at a top 50 undergraduate institution (Figure 23)

**Figure 23.**
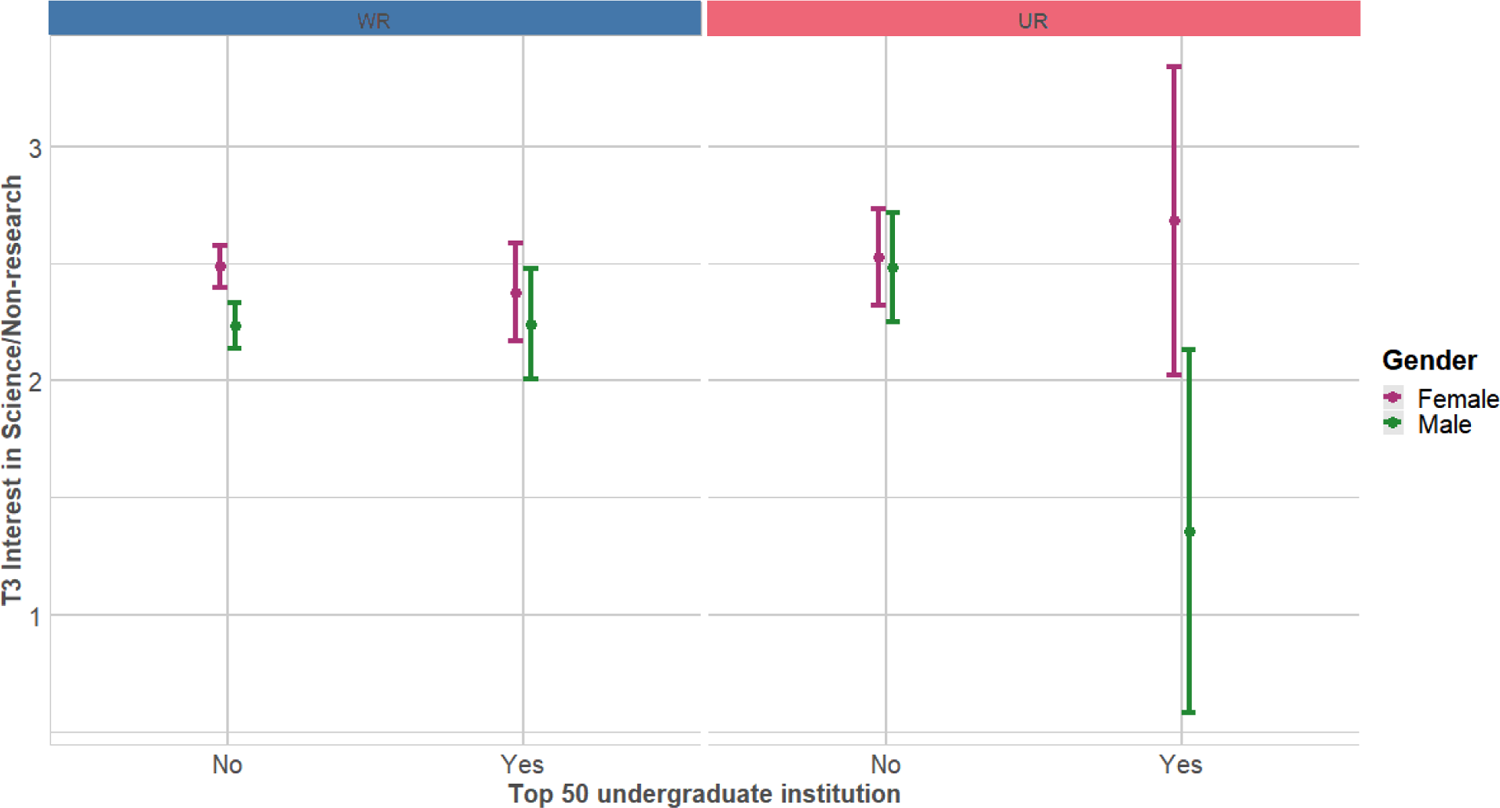
Top 50 undergraduate institution by gender by UR status interaction predicting T3 interest in science-related, non-research positions. Mean/error lines depicting current (T3) interest in science-related, non-research positions, split by gender and UR status, grouped by whether they had graduated from a top 50 undergrad institution. Dependent variable was current level of interest on a 4-point scale (where 1 represents “no interest” and 4 represents “strong interest”). Interaction was significant at p < 0.01 (Table 2; Table S10b).

**Figure 24.**
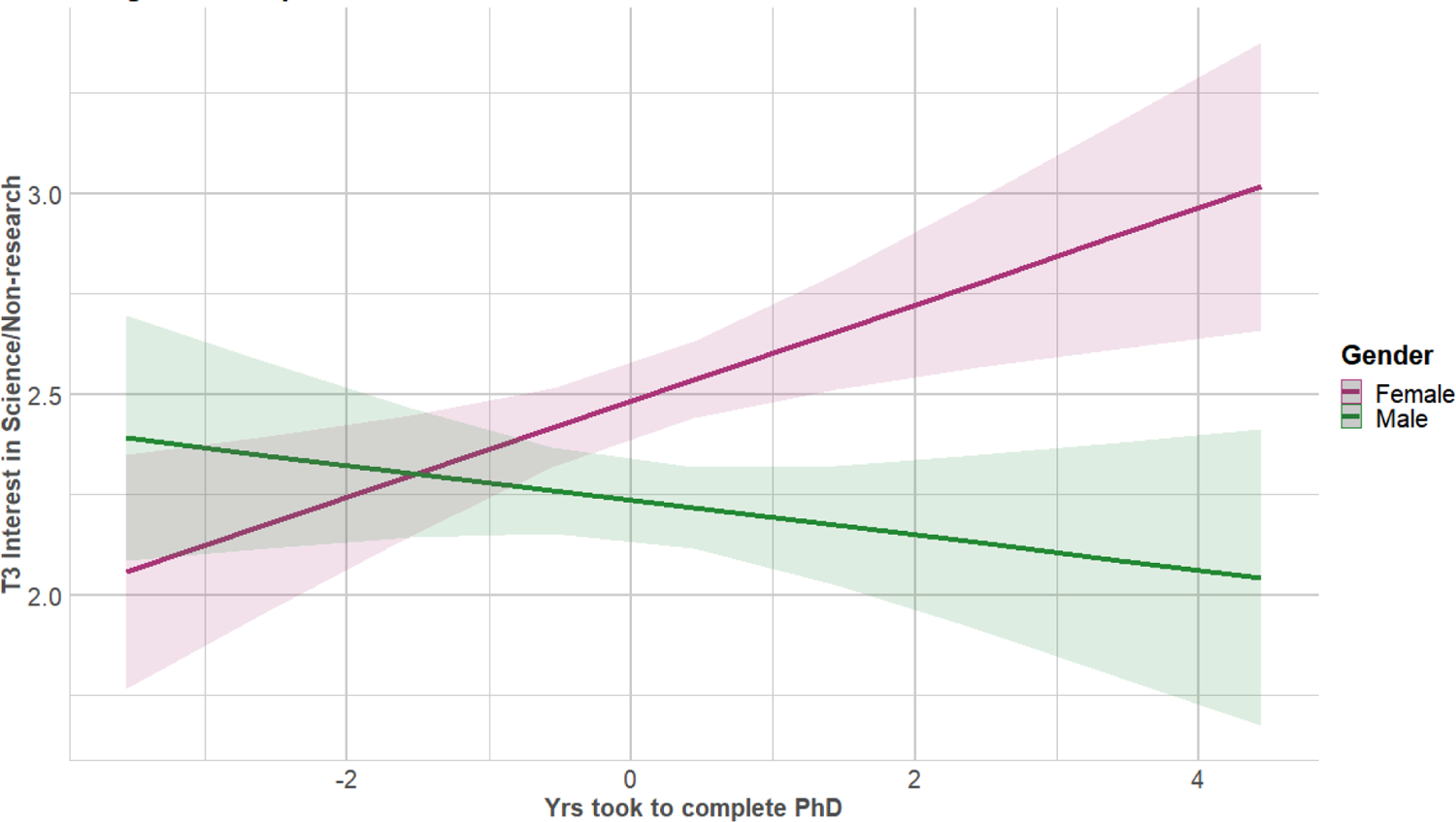
Time to complete PhD was associated with increased interest in science-related, non-research positions for females. Regression lines predicting current (T3) interest in science-related, non-research positions from number of years to complete PhD, split by gender. Dependent variable was current level of interest on a 4-point scale (where 1 represents “no interest” and 4 represents “strong interest”). Independent variable was number of years to complete PhD, centered for interaction (−4 to +5). Interaction was significant at p < 0.01 (Table 2; Table S10b).

**Figure 25.**
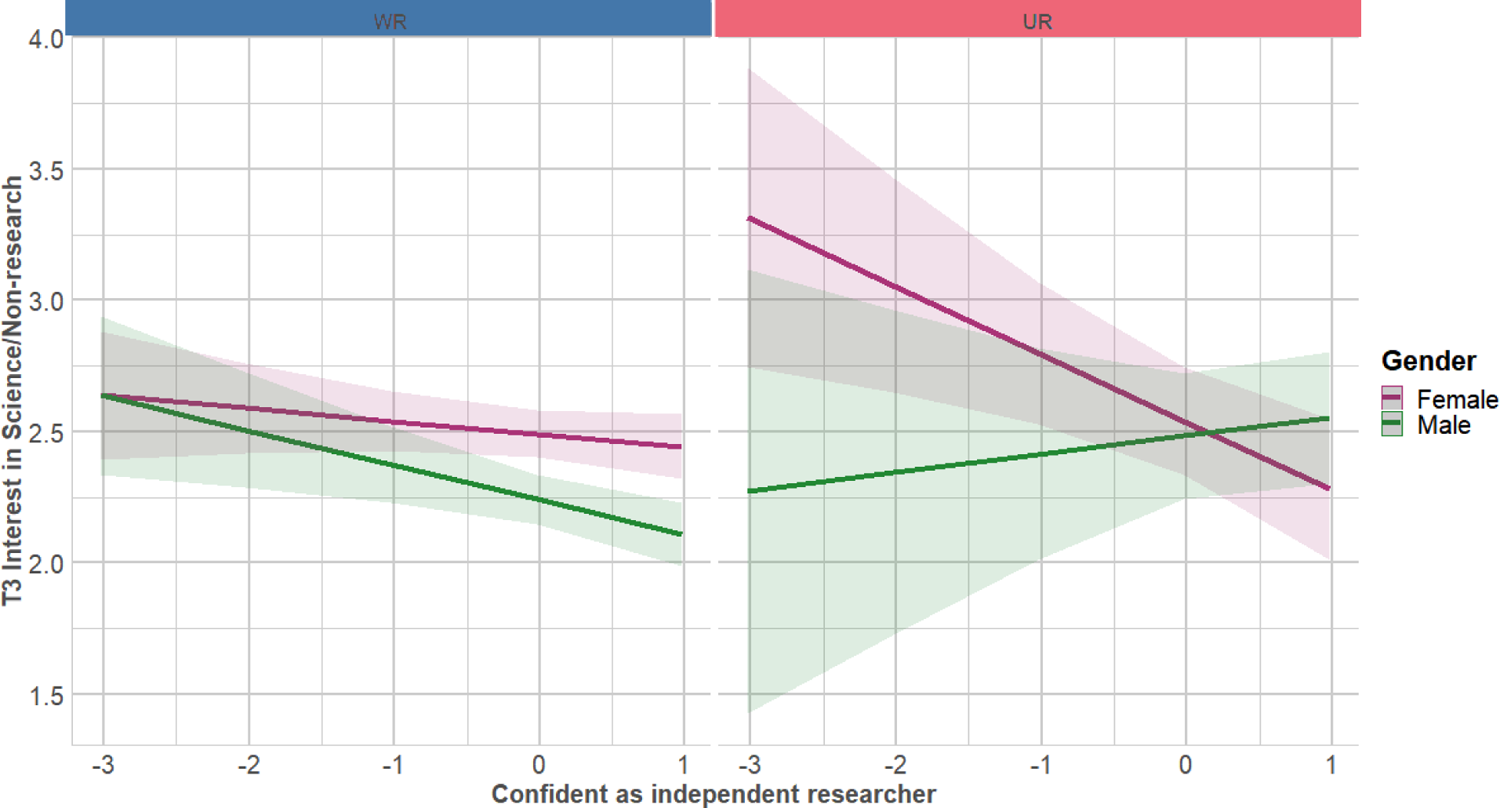
Confidence in being an independent researcher by gender by UR interaction predicting T3 interest in science-related, non-research positions. Regression lines predicting current (T3) interest in science-related, non-research positions from confidence in being an independent researcher, split by gender and UR status.

The second interaction was a 2-way interaction between Gender and years it took to complete PhD. While for females the longer it took to finish their PhD the more interested in science-related, non-research positions they were, for males the longer it took to finish their PhD the less interested in science-related, non-research positions they were (Figure 24). The third interaction was a 3-way interaction among Gender, UR status, and confidence in their potential to be an independent researcher. While for UR females and WR respondents, lower confidence in their potential to be an independent researcher was related to higher interest in science-related, non-research positions, for UR males lower confidence in their potential to be an independent researcher was related to lower interest in science-related, non-research positions (Figure 25).

The fourth interaction was a 2-way interaction between UR status and first-author publication rate. While for URs the lower their first-author publication rate, the more interested in science-related, non-research positions they were, for WRs there was no association (Figure 26). Finally, the fifth interaction was a 2-way interaction between Gender and the Features of Academia “Research/Autonomy” factor. While for females, the lower their interest in the research/autonomy features of academia the more interested in science-related, non-research positions they were, for males there was no association (Figure 27).

**Figure 26.**
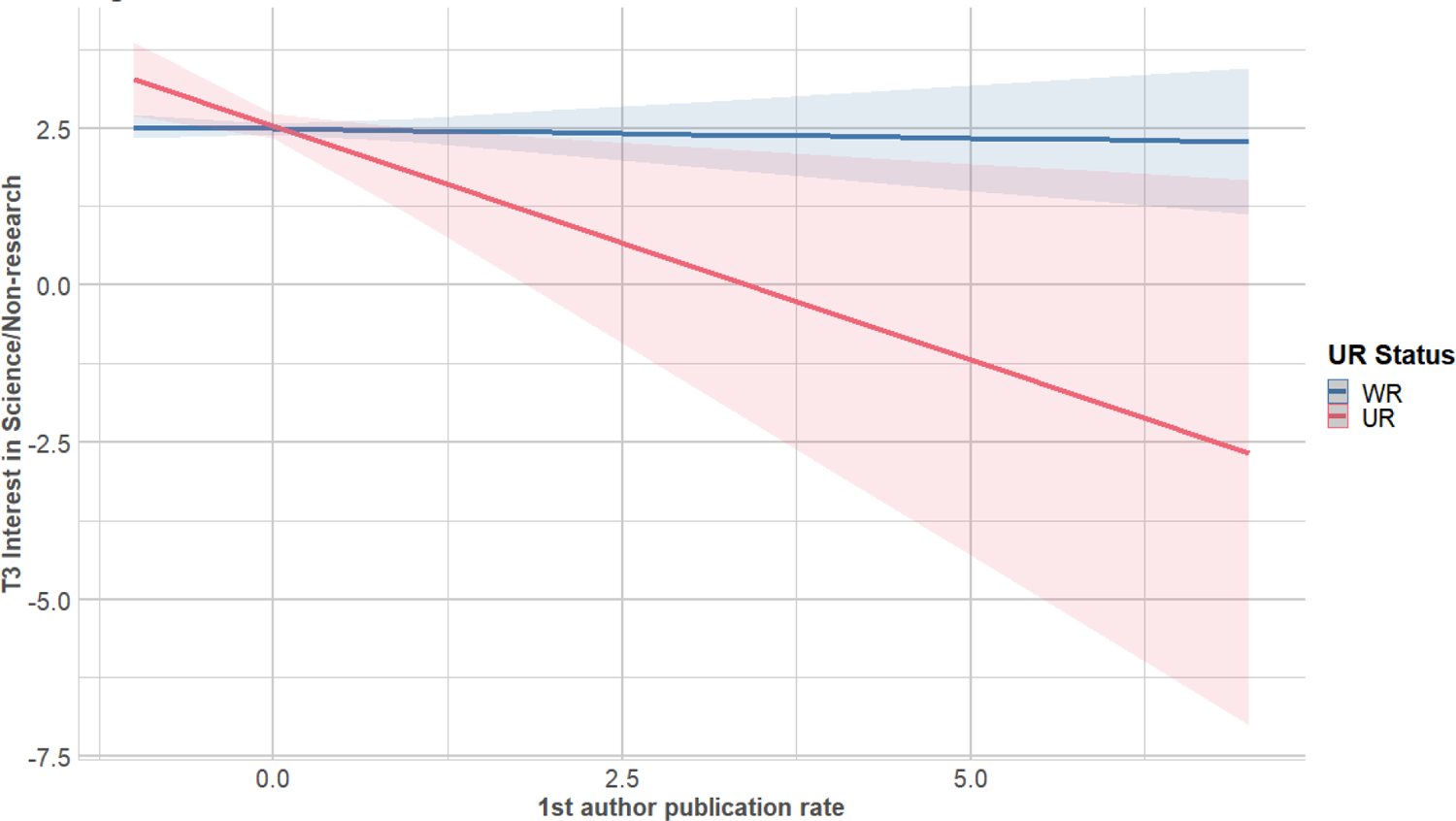
Lower first author publication rate was associated with increased interest in science-related, non-research positions for UR, but not WR, respondents. Regression lines predicting current (T3) interest in science-related, non-research positions from first author publication rate, split by UR status. Dependent variable was current level of interest on a 4-point scale (where 1 represents “no interest” and 4 represents “strong interest”). Independent variable was 1^st^ author publication rate, centered for interaction (−1 to +7). Interaction was significant at p < 0.05 (Table 2; Table S10b).

**Figure 27.**
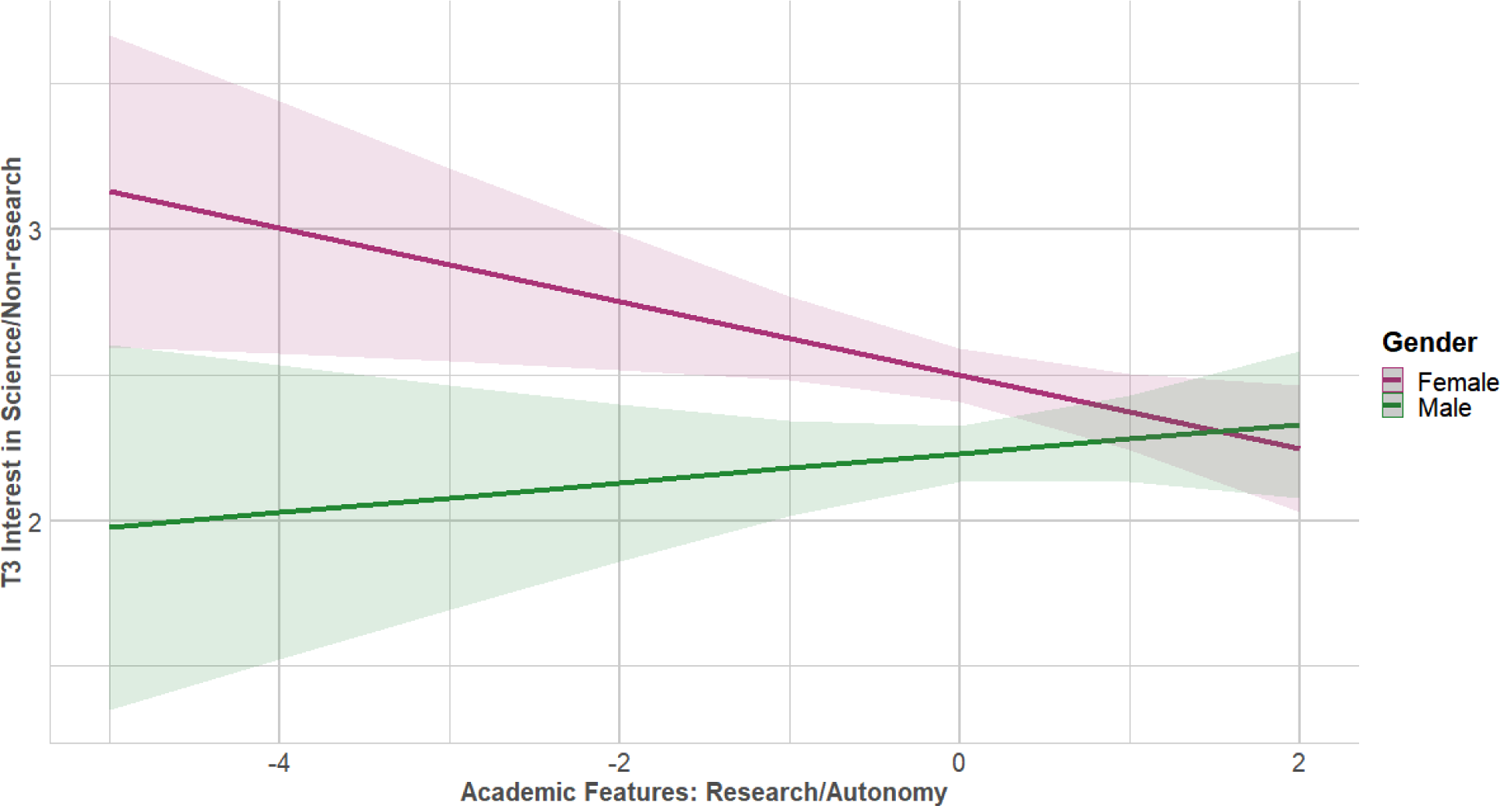
Valuing research/autonomy in academia was associated with a decrease in interest for science-related, non-research positions for females. Regression lines predicting current (T3) interest in science-related, non-research positions from rating of the importance of research/autonomy in academia, split by gender.

Dependent variable was current level of interest on a 4-point scale (where 1 represents “no interest” and 4 represents “strong interest”). Independent variable was confidence in being an independent researcher, centered for interaction (−3 to 1). Interaction was significant at p < 0.01 (Table 2; Table S10b).

Dependent variable was current level of interest on a 4-point scale (where 1 represents “no interest” and 4 represents “strong interest”). Independent variable was factor representing the importance of research/autonomy in academia. Interaction was significant at p < 0.05 (Table 2; Table S10b).

## DISCUSSION

This work was designed to investigate how career interest evolves over time among recent neuroscience PhD graduates, and whether differences in career interests are related to differences in social identity (gender, race, ethnicity), experiences in graduate school, and personal characteristics. We hypothesized that all three factors would be related to changes in career interest during graduate school and beyond.

Several caveats should be mentioned. First, we analyzed survey results from 1,479 US citizen or permanent resident PhD holders who had previously applied for or received NINDS support and obtained their PhD in 2008 or later. Although non-citizens make up a sizeable number of trainees (and the majority of postdocs) in the US (National Science Foundation, 2015), NINDS training and career development awards are generally only available to US citizens during training (with a few exceptions), and thus our analysis was limited to that population. Second, although our response rate of ∼36% of the population (i.e., 45% of opened emails) is in line with other surveys, it is not a random sample. The sample likely has an overrepresentation of people such as those likely to respond to surveys, those with an interest in the topic, and, due to the conventionally public nature of academic contact information, those in academic positions. However, we have a similar sample demographic as other studies (Gibbs et al., 2014). Third, although we collected information on disability status, persons with a disability made up less than 3% of the sample, so were not included as a separate analysis group because of the small sample size. Fourth, the number of explanatory (independent) variables explored in our analyses was quite large. Steps taken to minimize Type I error included false discovery rate control and the use of effect size cutoffs. Fifth, this survey was conducted before the COVID-19 pandemic and does not reflect the likely sizeable influence of this event. Finally, respondents were asked to retrospectively rate their career interests at the start and end of their PhD training. As such, these ratings represent their *current* view of their interest in the past. This fits with the goals of the survey, which include understanding respondents’ views of their career trajectories.

In our sample, we saw similar trends as other retrospective and cross-sectional studies: interest in academia, both research- and teaching-focused positions, goes down over time while interest in non-academic careers goes up over time (Fuhrmann et al., 2011; Gibbs et al., 2014, 2015; C. M. Golde & Dore, 2001; Goulden et al., 2011; Roach & Sauermann, 2017; Sauermann & Roach, 2012; but see: Wood et al., 2020). However, there are differences between sub-groups in our sample, as explored below.

One common factor across all groups is the outsize influence of the academic advisor on career interest. During training, the PhD or postdoc advisor can be a source of guidance, support, and networking opportunities (Barnes, 2009). Alternatively, this relationship can also be a source of frustration, disappointment, and discouragement (Barker, 2011; Barnes, Williams, & Archer, 2010; Burt, McKen, Burkhart, Hormell, & Knight, 2020; Thomas, Willis, & Davis, 2007; Zhao, Golde, & McCormick, 2007).

Previous studies have shown that quality supervision and mentoring are related to persistence in STEM (Berg & Ferber, 1983; de Valero, 2001; Estrada, Hernandez, & Schultz, 2018; Gardner, 2008; Palmer & Young, 2008). We found that career advice from PhD advisors perceived as helpful to the trainee is related to increases in interest in academic research and decreases in interest in science non-research during graduate school. In addition, helpful career advice for postdocs from their advisors is related to higher current interest in academic research. This emphasizes the need for consistent high-quality mentorship and is being incorporated into the NINDS strategic plan for 2021-2025 (NINDS, 2021).

The drop in the representation of trainees from marginalized backgrounds over the course of training has commonly been described as “the leaky pipeline” (McGee, Saran, & Krulwich, 2012; Miller & Wai, 2015), a metaphor that assumes that at PhD program entry all trainees aspire to a faculty research position. In contrast, we found that not only do some individuals enter graduate school with low interest in faculty research positions (and this is the biggest single predictor of interest at end of PhD), but also that interest may vary by social identity group. In particular, women reported being less interested in academic research careers at the start of their PhD—and their interest drops more over time—and are correspondingly more interested in science non-research at the start of their PhD and gain more interest over time compared to their male counterparts.

What may explain these differences? In the realm of individual preferences, women in our study report a higher importance of work/life balance as an aspect of their careers. This is in line with a large body of research that has found that work/family balance challenges influence female graduate students and postdocs away from faculty careers (Lambert et al., 2020; E. D. Martinez et al., 2007; Sassler, Glass, Levitte, & Michelmore, 2017). Beddoes and Pawley (2014) found that faculty identified similar family-related challenges as contributing to low numbers of female STEM faculty. However, they also stressed that the discourse of “personal choice” within an inequitable situation can obscure the systemic pressures that different groups face. For example, in male-female partnerships, a disproportionate amount of family and household responsibilities falls to women who work full-time (Glynn, 2018; Schiebinger & Gilmartin, 2010) and, in particular, the difficulties of balancing caregiving responsibilities with full-time work in STEM may explain why women are more likely than men to leave academia or STEM employment entirely after having children (Cech & Blair-Loy, 2019; Goulden et al., 2011; L. R. Martinez, O’Brien, & Hebl, 2017; Wolfinger, Mason, & Goulden, 2008), and may contribute to lower publication rates among women (Morgan et al., 2021). This has never been in sharper relief than during the current pandemic, which has disproportionately affected women, especially women with young children (Gewin, 2020; Kramer, 2020; Myers et al., 2020; National Academies of Sciences, Engineering, and Medicine, 2021). These issues are further compounded for women of color in academia (Guy & Boards, 2019; Kachchaf, Ko, Hodari, & Ong, 2015). As Beddoes and Pawley (2014) point out, more important than describing the choices that are made is understanding why other careers are more appealing than academic research for certain groups, instead of “writing it off as a matter of individual priorities.” In our study, among all respondents, rating work/life balance as important led to lower interest in academic research and higher interest in academic teaching. Thus, it is not academia itself that is considered incompatible with work/life balance, but rather something unique to academic research-focused positions rather than teaching-focused positions.

Institutions are beginning to recognize that challenges surrounding work/life balance exist and differentially affect mothers in science, including the exorbitant cost of childcare and navigating frequent travel to conferences (Calisi & a Working Group of Mothers in Science, 2018; although see Cummins, 2005 for a discussion of the lack of support for childless women in academia). To that end, some universities offer subsides and other forms of support to working parents (see programs at University of Iowa, Brown University, Stanford University, University of Michigan, University of Massachusetts Amherst, and more). NIH has several programs and policies to support working parents: 1) Most recently, trainees and fellows supported by National Research Service Awards may now request support for childcare costs (Lauer, 2021); 2) Early Stage Investigator status and K99/R00 eligibility is automatically extended by a year for individuals who have had a child after completing their terminal degree, and extensions are available for other life events (National Institutes of Health, 2018, 2019a); 3) NIH has made supplements available to primary investigators of R and K awards who have experienced critical life events (such as childbirth, adoption, or elder care) that have the potential to impact research progress (National Institutes of Health, 2020a, 2020b). More acutely, the COVID-19 pandemic has put into sharp relief both the tensions and some possible solutions for managing work-life integration (Malisch et al., 2020). Additional policies and programs in this area may have a positive impact, not only on female persistence in academic research, but on the culture of work-life integration in science, allowing scientists to navigate many of life’s challenges, such as medical events or other emergency circumstances.

In our survey, women also report that the structural aspects of academia (grant funding, job market, promotion) reduce their interest in academia more than they do for men. Among the entire sample, more negative feelings about the structural aspects of academia (“Funding/Job Market/Promotion”) were related to lower interest in academic and non-academic research and higher interest in science-related non-research careers. While increasing support for parents provides a clear area of intervention; structural issues around scarce funding and attendant issues such as the job market and promotion are harder to ameliorate. NINDS has directed particular attention towards supporting junior faculty in the hopes of easing those concerns, through the Early Stage Investigator policy, postdoctoral to faculty transition awards (K99/R00), the K01 diversity faculty award, research education awards (R25s) for career development of junior faculty, and participation in the NIH Faculty Institutional Recruitment for Sustainable Transformation (FIRST) Program to support cohort hiring of scientists committed to diversity and inclusive excellence, among other efforts.

In addition to differences in their ratings of interest in the different career types, women reported issues that may be roadblocks to academic research careers, such as poorer relationships with their PhD advisors, lower first-author publication rates, and less confidence in their ability to be independent researchers compared to men. For UR women in particular, having lower confidence in their ability to be an independent researcher is related to higher interest in science-related non-research. The impact of bias, exclusion, and structural racism likely contribute to these findings and can become the drivers for what is commonly known as “imposter syndrome,” a controversial term that centers the problem in the individual, rather than the system (Tulshyan & Burey, 2021). These results mirror findings of other studies (Cheryan, Ziegler, Montoya, & Jiang, 2017; E. D. Martinez et al., 2007), and may stem either from women underestimating their ability or men being over-confident in their ability to be independent researchers (Bench, Lench, Liew, Miner, & Flores, 2015). Confidence in one’s ability to perform a task, self-efficacy, has been shown to be an important piece of persistence in STEM careers (Berg & Ferber, 1983; Byars-Winston, Branchaw, Pfund, Leverett, & Newton, 2015).

Members of marginalized groups also report factors that are roadblocks for academic research careers. For example, UR respondents reported lower first-author publication rates. This finding is in line with other work showing lower numbers of publications for women and/or UR scientists (Lambert et al., 2020; Mendoza-Denton et al., 2017; Pezzoni, Mairesse, Stephan, & Lane, 2016). For UR respondents, having a lower publication rate is related to higher interest in non-research science careers. This analysis cannot determine directionality of this association, but the relationship is likely complicated—that is, experiences and perceptions in graduate school influence publication rate and career interests (Fisher et al., 2019), while career interests in turn may influence publication rate. There is also evidence of systemic bias within publishing, as women are consistently underrepresented in scientific journals as authors, in citation counts, and in reference lists (Caplar, Tacchella, & Birrer, 2017; Dworkin et al., 2020; Larivière, Ni, Gingras, Cronin, & Sugimoto, 2013; Shen, Webster, Shoda, & Fine, 2018), and white authors are over-represented among citation lists (Bertolero et al., 2020). Journals are increasingly turning towards new models to address these findings, such as double-blind review (Bernard, 2018) and inclusion and diversity statements for journal submissions (Sweet, 2021).

Consistent with similar studies, our study shows UR respondents feel less like they were a part of the social and intellectual community during both their PhD and postdoc training (Fisher et al., 2019; Stachl & Baranger, 2020). This finding is not particularly surprising, especially in light of the poignant personal stories recently shared from #BlackintheIvory hashtag on Twitter and other venues that illuminate this lack of inclusion in the research enterprise (Armstrong, Lomax, Traylor-Knowles, Samba-Louaka, & Towers, 2020; Dzirasa, 2020; Karanja, Bostic, DaCrema, & Pacheco, 2020; Odekunle, 2020). Sense of belonging has been shown to mediate desire to pursue further studies in a discipline (Gardner, 2008; Good, Rattan, & Dweck, 2012), and a feeling of isolation is related to higher attrition rates from PhD programs (Abbe H. Herzig, 2004; Lovitts, 2001). Conversely, institutional attention to both STEM culture and institutional climate has the potential to enhance persistence in science (K. Griffin, 2018). A fundamental part of the adoption of an identity of a scientist is participation in and integration with a community of practice during academic training (A H Herzig, 2004). Barriers to full participation in the social and intellectual community of the PhD and postdoc training can have a profound effect on development of scientific identity. These barriers may be explicit, in the form of hostile or racist work environments or lack of access to professional development resources or mentorship, or may be implicit, in the form of differing values (Aelenei, Martinot, Sicard, & Darnon, 2020; Gibbs & Griffin, 2013) or alienating cultural assumptions such as a culture perceived as masculine, focused on self-enhancement, or individualistic (Aelenei et al., 2020; Cheryan et al., 2017; A H Herzig, 2004). Supporting the latter possibility, both members of underrepresented groups and women in our study report a lower importance of autonomy in their careers, and among the entire sample, negative feelings about the “Autonomy/Research” aspects of academia were related to lower interest in research careers. As the importance of team science increases in biomedicine, institutions may re-think models or messaging around the idea of the “lone genius” in favor of collaborative structure and recognition (Bennett, Gadlin, & Marchand, 2018). Along those lines, NIH has recently made more explicit calls for team research in investigative neuroscience at different stages and on various scales (David, Fang, Peng, & Gnadt, 2020).

The lack of community felt at their home institution may drive UR trainees to find support elsewhere, as demonstrated by reports of beneficial relationships with faculty outside their PhD institutions (Burt, Williams, & Palmer, 2018; Ellis, 2000). Our study shows that faculty from other institutions had a strong influence on UR respondents. These differences were even more pronounced in UR women during their PhD training compared to UR men, suggesting an intersectional or compounding effect of multiple marginalized social identities (Guy & Boards, 2019; Ong, Wright, Espinosa, & Orfield, 2011). This result also may speak to the success of networking and professional development programs for underrepresented scientists. NINDS supports programs that strive to equip underrepresented scientists with the tools needed to successfully navigate the challenges of academic life and leverage national peer support systems and mentoring networks outside of the primary institution (Jones-London, 2020). These NINDS-funded programs include Society for Neuroscience’s Neuroscience Scholars Program (National Research Council, 2013), the BRAINS program (Margherio, Horner-Devine, Mizumori, & Yen, 2016; Yen, Horner-Devine, Margherio, & Mizumori, 2017), the TRANSCENDS Program (Tagge, Lackland, & Ovbiagele, 2021), and the NIH Blueprint-funded ENDURE and D-SPAN Programs (Jones-London, 2020).

Additionally, the NIH FIRST program, NSF ADVANCE, and the HHMI Inclusive Excellence programs, among others, are attempting to increase inclusion and equity by changing the institutional culture, instead of focusing on “fixing” the individual.

## Conclusion

We saw repeated evidence that individual preferences about careers in general, and academic careers specifically, predict current career interest. These preferences were mediated by social identity and experiences in graduate school and postdoctoral training. Our findings highlight the outsized influence of the advisor in shaping a trainee’s career path, and the ways in which academic culture is perceived as unwelcoming or incongruent with the values or priorities of certain groups. For women, issues of work/life balance and structural issues of academia, and for UR women in particular, lower confidence in their ability to be an independent researcher, affected their interest in academia. Both women and underrepresented men in our study report a lower importance of autonomy in their careers. UR respondents report feeling less like they were a part of the social and intellectual community, but have formed beneficial relationships with faculty outside their PhD institutions that are—for UR women— associated with increased interest in academia. Although the effect sizes are mostly modest for these findings, the results do not reflect shared variance with other variables in the analysis, and likely underestimate their true influence. Our findings suggest several areas for positive growth, ways to change how we think about the impact of mentorship, and policy and programmatic interventions that extend beyond trying to change or “fix” the individual and instead recognize the systemic structures that influence career choices. Importantly, we recognize that not all PhD-holders will or should express interest in academic research, but that career path should be equally available and welcoming to all so that our nation can benefit from the full potential of its diverse workforce.

## METHODS

### Sample

The study population was composed of 1) recent doctoral recipients (CY2008 or later) who were 2) US citizens or permanent residents and 3) had applied for NINDS funding or have been appointed to NINDS training (T32) or research education grants (R25). In addition to capturing post-trainees across a decade, the year 2008 was chosen as a cutoff because 2003 marked a clear turning point in NIH funding: between 1998 and 2003 the NIH budget almost doubled, whereas from 2003 to 2017, when this survey was conducted, NIH funding plateaued in real dollars and decreased in relative purchasing power (FASEB Office of Public Affairs, 2020; Sekar, 2020). Those who were in graduate school between 2003 and the present likely had very different experiences than those who entered graduate school earlier than 2003. Since the average time to Neuroscience PhD is 5-6 years (Lorden, Kuh, & Voytuk, 2011), those graduating in 2008 entered around 2003—hence, the choice of 2008 as the cut-off point.

Potential participants for this study were identified within the NIH’s Information for Management Planning Analysis and Coordination II (IMPACII) database, a database containing administrative data from all extramural grant applications (Institute of Medicine (US) Council on Health Care Technology & Goodman, 1988). A total of 7,405 eligible or likely eligible individuals were identified in IMPACII through these searches. Citizenship and year of PhD conferral information was available for some, but not all, individuals, so not all identified individuals were eligible; respondents were screened for eligibility according to the three criteria above at the beginning of the survey. An email list containing every email address available in the IMPACII system for the 7,405 identified individuals was created to allow email outreach for the survey. Approximately half of US citizen or permanent resident neuroscience PhD recipients are supported by NIH during their PhD (NIH Data Book, n.d.); we do not have exact data on the proportion supported by NINDS nor the proportion that apply for NINDS funding but do not receive it; both types of individuals were eligible for this survey. As determined by the NIH Office of Human Subjects Research, federal regulations for the protection of human subjects do not apply to this activity.

### Dissemination and Data Collection

Unique survey invitations were sent on May 10, 2017 through SurveyMonkey (SurveyMonkey, Inc., 2015) to all identified email addresses (9,758 addresses; an average of 1.3 email addresses per person). Follow-up invitations for those who had not responded were sent through SurveyMonkey every two weeks. For emails that were undeliverable or “bounced,” an attempt was made to find a current email address through online searches. Additionally, mentors of eligible F31 and F32 applicants were asked to forward information about the survey to their trainees’ current email addresses. If individuals independently inquired about the survey, eligibility was confirmed before sending a survey invitation.

All participants consented to participation in the study. All survey responses were anonymous. At survey close, on July 1, 2017, a total of 5,935 emails (61%) had been opened and 3,823 emails (39%) were undeliverable or unopened. The survey received 2,675 responses (approximately 36% of identified individuals). Of these responses, 2,310 were complete, 250 were ineligible, and 115 gave a partial response. Of the 2,310 complete and eligible response, 65 were from participants who filled out the survey more than once. For multiple responses from the same participant, only the first response was kept, for a final total of 2,242 complete, eligible, and unique responses.

### Definitions and Sample Refinement

Several other criteria were applied to further refine the sample for analysis. First, since gender and race/ethnicity were of primary interest for this article, the responses that did not include that information were excluded, leaving 2,065 responses. These responses included all who identified as either male or female and may include transgender respondents who identify as either male or female (only two participants indicated “other” and wrote in a response for gender; they were not included in the sample). Respondents from white and/or Asian backgrounds are referred to as well-represented (WR), while respondents from American Indian/Alaska Native, Black/African American, Hispanic/Latino, and/or Native Hawaiian/Pacific Islander backgrounds are referred to as underrepresented (UR), according to the NSF definition (National Science Foundation, 2015). Second, since the original survey was also aimed at current PhD students, who would not have “end of graduate school” ratings of interest, all current students were dropped, leaving a final sample size of 1,479. Disability status was collected, but persons with a disability made up less than 3% of the sample, so were not included as a separate analysis group because of the small sample size.

### Survey

The survey was a 57-question instrument administered at a single point in time. It asked about respondents’ career interest; experiences in graduate school and postdoctoral training; feelings about careers in general; objective measures of research experience and productivity; and basic demographics (see Survey Text in the *Supporting Information Appendix*). Questions were iteratively developed by synthesizing from several sources, conducting cognitive testing interviews, and refining language where necessary (Gibbs & Griffin, 2013; Gibbs et al., 2015; K. Griffin, Gibbs, Jr., Bennett, Staples, & Robinson, 2015; Layton et al., 2016; Malley, Churchwell, & Stewart, 2006; Meyers et al., 2016; National Institutes of Health, 2012, 2019b; National Postdoctoral Association, n.d.; Sauermann & Roach, 2012; M. V. Sinche, 2016; M. Sinche et al., 2017; US National Science Foundation, 2016a, 2016b; Yoder & Mattheis, 2016).

Respondents were asked to rate their interest in pursuing each of the following career pathways at three time points: the start of their PhD program, the end of their PhD program, and currently. These pathways were: Academic position, research focus (includes physician-scientist); Academic position, teaching focus; Non-academic research (e.g., research in industry, biotech, or government settings); Science-related, non-research (e.g., science outreach, communication, policy, advocacy, or administration). Respondents were also asked about their interest in Other, non-science-related careers; as this did not measure a specific career path, but a variety of possible careers, and to reduce the number of variables and analyses, these responses were not analyzed. Interest was measured on a four-point Likert-type scale where 1= No interest, 2=Low interest, 3=Moderate interest, 4=Strong interest.

Respondents were also asked about their social identity (specifically gender and race/ethnicity), experiences in graduate and postdoctoral training, personal characteristics, and objective measures (Table S2).

Experiences in training included: various aspects of their relationship with their primary training advisor during graduate and postdoctoral training (5-point scale from “very negative” to “very positive”); sources and helpfulness of support and career advice during the graduate and postdoctoral training (4 point scale from “no guidance provided” to “very helpful”); feelings of social and intellectual belonging to lab/research group and department/program during graduate and postdoctoral training (5-point scale from “strongly disagree” to “strongly agree”).

Personal characteristics included: confidence in one’s potential as an independent researcher (measured on a 5-point agreement scale where 1 was “strongly disagree” and 5 was “strongly agree”); aspects of the career or work environment most important to the respondent (choose up to top 5); and features of academia that increase or decrease desire to become a faculty member (5-point scale from “greatly decrease” to “greatly increase”).

Objective measures included years of research prior to PhD program, total years of research, years to complete PhD, total time in postdoctoral training, years since PhD completion, support by NIH prior to the PhD program, first-author publication rate (first-authored publications/total years performing research), time to PhD completion, and undergraduate or doctoral degree from a top 50 research university (as measured by research and development expenditures; National Science Board, 2016).

## ANALYSIS

### Variable Testing and Data Reduction

This work was designed to investigate how career interest evolves over time, and whether changes and/or differences in career interests are associated with gender and race/ethnicity, experiences in graduate school and postdoctoral training, and personal characteristics. Outcome variables were ratings of interest in the different career types, represented on a 4-point scale, at three time-points: start of PhD program (T1), end of PhD program (T2), and current (T3). All independent (explanatory) variables were split between those used to predict interest at the end of respondents’ PhD programs (e.g. feelings of belonging during PhD), and those used to predict current interest (e.g. feelings of belonging during postdoctoral training).

Although the outcome variables are ordinal in nature, it was preferable to treat them as interval in these analyses. Accordingly, we tested their suitability for use as interval variables by using a procedure outlined by Jacoby (Jacoby, 1999, 2012). First, the ordinal variables were converted though ALSOS optimal scaling to create interval-level representations. Then the interval variables were correlated with the original variables and we found that a very strong linear relationship existed: correlations ranged from a low of 0.9644 to a high of 0.9997, all significant at the p<0.0001 level. This process was repeated with all other ordinal variables in the study, and correlations ranged from 0.9377 to 0.9991, all significant at the p<0.0001 level. Therefore, we felt comfortable treating all ordinal-level variables as interval level variables in the analyses.

Finally, data reduction was performed for several constructs in order to reduce multicollinearity and the problem of multiple comparisons and Type I error. We used factor analysis to reduce these constructs (e.g., relationship with advisor) into latent factors. Details of the factor analyses can be found in Table S11 in the *Supporting Information Appendix*.

### General Notes

All data analyses were conducted using versions 3.6.2 or 3.6.3 of the R program (R Core Team, 2018). Individual packages are cited in text when referenced. All interactions were evaluated in the context of component main effects and all lower-level interactions.

### Gender and UR Status Differences: Logistic Regression, Multinomial Logistic Regression, and ANOVA

Gender and UR Status differences on explanatory variables were investigated through three different procedures, depending on the nature of the explanatory (here dependent) variable. For all three sets of analyses, each analysis used the explanatory variables as dependent variables, and Gender, UR Status, and their interaction as the independent variables. False Discovery Rate (FDR) was controlled using Benjamini and Hochberg’s (1995) procedure, which was applied to each analysis using the “BH” option on the mt.rawp2adjp function of R’s Multtest package (Gentleman, Carey, Huber, Irizarry, & Dudoit, 2005).

Gender and UR Status differences on the four dichotomous explanatory variables were investigated using logistic regressions (using glm from the base package with family = “binomial” in R). Gender and UR Status differences on the two multinomial explanatory variables were investigated using multinomial logistic regressions (using multinom from the nnet package in R, Venables & Ripley, 2002). Statistics for individual terms were computed by successively contrasting statistics from the full model to statistics from three sub-models that each had a different term removed using the ANOVA test for model comparison. Finally, Gender and UR Status differences on the 33 continuous explanatory variables were investigated using ANOVA (using aov from the base package in R).

### Differences in Career Interest Ratings Over Time: Repeated Measures MANOVAs

The 4 omnibus repeated measures MANOVAs (one for each career type) were conducted using lmer from the lme4 package (Bates, Mächler, Bolker, & Walker, 2015) in R (R Core Team, 2018). Each MANOVA had all three interest ratings over time for a single career type as dependent variables, and Time as the independent variable. Follow-ups were also conducted using lmer. Estimated marginal means and effect sizes were computed with emeans and eff_size from the emmeans package (Lenth, 2020). Follow-ups were only conducted for significant main effects or interactions from the omnibus MANOVAs.

### Regressions Investigating Change in Career Interest During Graduate School

FDR was controlled in all analyses using Benjamini and Hochberg’s (1995) procedure, as above. In the first step, correlations between outcomes and explanatory variables were computed using corr.test from the psych package in R (Revelle, 2019). Outcome variables were the four Time 2 (end of PhD) ratings of interest in different careers with Time 1 (start of PhD) ratings of interest for the same career covaried out. Explanatory variables were limited to those that asked about graduate school and non-time-bound questions. Explanatory variables were carried over to third step regressions if their correlations with outcome variables were significant to at least the p<0.05 level and they accounted for at least 2% of the variance in the outcome variables.

In the second step, interactions between graduate school-era explanatory variables, Gender, and UR Status predicting change in interest in the different careers over graduate school were tested using glm from the base package in R. Three-way interactions (Gender by UR Status by explanatory variable) for each of the explanatory variables predicting change in interest over graduate school were tested first. If the 3-way interaction was not significant, then all 2-way interactions predicting change in interest over graduate school were tested in a new equation that did not include the 3-way interaction (to preserve shared variance). Interactions were carried over to third step regressions if they were significant at least the p<0.05 level (no effect size requirements were utilized because there is no consensus effect size measure for moderation in the literature, Smithson & Shou, 2017).

In the third step, regressions were computed using lm from the base package in R. This step proceeded in 2 phases. In phase 1, regressions were computed including all explanatory variables and interactions brought forward from steps 1 and 2 (above). In phase 2, all variables and interactions that had semi-partial correlation values of 0 in phase 1 were dropped, and new regression equations were computed. The effect size threshold for reporting results from these analyses was set to 1.0% unique variance captured, Cohen’s small effect for multiple regressions, because each variable’s loss of shared variance in the fitting of the multiple regression model makes the amount unique variance captured a stringent test.

Statistical assumptions for OLS regressions were tested for all regressions in this step using the Breusch-Pagan test (for heteroskedasticity) (bptest) and the Durbin-Watson test (for autocorrelated errors) (dwtest) from the lmtest package in R (A Zeileis & Hothorn, 2002). The means of the residuals for each regression were also checked to ensure that they were close to 0. Results showed that all the residual means were close to 0 and that none of the Durbin-Watson tests were significant. Many of the Breusch-Pagan tests were significant, however, and thus all regression coefficients, standard errors, and t values were corrected for heteroskedasticity using vcovHC from the sandwich package in R (Achim Zeileis, 2004).

### Regressions Predicting Current Interest

These analyses were conducted identically to the regression analyses above. Outcome variables were the four Time 3 (current) ratings of interest in the four different careers. The full set of explanatory variables were used as predictors.

## Supporting information

Survey Text

## ACKNOWLEDGMENTS

We thank the Diversity Working Group at NINDS for their feedback and input (especially Edgardo Falcon-Morales, Katie Pahigiannis, Ashlee Van’t Veer, and Letitia Weigand for initial survey design). Thank you also to DeAnna Adkins, Devon Crawford, Robert Finkelstein, Kenneth Gibbs, Jr., Jordan Gladman, Kalynda Gonzales Stokes, Lyn Jakeman, David Jett, Crystal Lantz, Marguerite Matthews, DP Mohapatra, Nina Schor, Ashlee Van’t Veer, and Robert Zalutsky for feedback on the manuscript draft. We thank Walter Koroshetz and Janine Clayton and the NIH Office of Research on Women’s Health for their leadership and financial support.

## Notes

### Competing Interest Statement

The authors have declared no competing interest.

## REFERENCES

1. Aelenei, C., Martinot, D., Sicard, A., & Darnon, C. (2020). When an academic culture based on self-enhancement values undermines female students’ sense of belonging, self-efficacy, and academic choices. The Journal of social psychology, 160(3), 373–389. doi:10.1080/00224545.2019.1675576

2. Armstrong, A., Lomax, J., Traylor-Knowles, N., Samba-Louaka, A., & Towers, C. (2020). On being black in the ivory tower. Cell, 183(3), 559–560. doi:10.1016/j.cell.2020.10.006

3. Barker, M. J. (2011). Racial context, currency and connections: Black doctoral student and white advisor perspectives on cross-race advising. Innovations in Education and Teaching International, 48(4), 387–400. doi:10.1080/14703297.2011.617092

4. Barnes, B. J. (2009). The nature of exemplary doctoral advisors’ expectations and the ways they may influence doctoral persistence. *Journal of College Student Retention: Research*, Theory & Practice, 11(3), 323–343. doi:10.2190/CS.11.3.b

5. Barnes, B. J., Williams, E. A., & Archer, S. A. (2010). Characteristics that matter most: doctoral students’ perceptions of positive and negative advisor attributes. NACADA Journal, 30(1), 34–46. doi:10.12930/0271-9517-30.1.34

6. Bates, D., Mächler, M., Bolker, B., & Walker, S. (2015). Fitting linear mixed-effects models using lme4. Journal of statistical software, 67(1), 1–48. doi:10.18637/jss.v067.i01

7. Beddoes, K., & Pawley, A. L. (2014). Different people have different priorities‘: work–family balance, gender, and the discourse of choice. Studies in Higher Education, 39(9), 1573–1585. doi:10.1080/03075079.2013.801432

8. Bench, S. W., Lench, H. C., Liew, J., Miner, K., & Flores, S. A. (2015). Gender gaps in overestimation of math performance. Sex roles, 72(11-12), 536–546. doi:10.1007/s11199-015-0486-9

9. Benjamini, Y., & Hochberg, Y. (1995). Controlling the false discovery rate: A practical and powerful approach to multiple testing. Journal of the Royal Statistical Society: Series B (Methodological), 57(1), 289–300. doi:10.1111/j.2517-6161.1995.tb02031.x

10. Bennett, L. M., Gadlin, H., & Marchand, C. (2018). *Collaboration and Team Science: A Field Guide* (2nd ed.). National Cancer Institute.

11. Berg, H. M., & Ferber, M. A. (1983). Men and women graduate students: who succeeds and why? The Journal of higher education, 54(6), 629. doi:10.2307/1981934

12. Bernard, C. (2018). Editorial: Gender Bias in Publishing: Double-Blind Reviewing as a Solution? eNeuro, 5(3). doi:10.1523/ENEURO.0225-18.2018

13. Bertolero, M. A., Dworkin, J. D., David, S. U., Lloreda, C. L., Srivastava, P., Stiso, J., … Bassett, D. S. (2020). Racial and ethnic imbalance in neuroscience reference lists and intersections with gender. BioRxiv. doi:10.1101/2020.10.12.336230

14. Burt, B. A., Williams, K. L., & Palmer, G. J. M. (2018). It takes a village: the role of emic and etic adaptive strengths in the persistence of black men in engineering graduate programs. American educational research journal, 000283121878959. doi:10.3102/0002831218789595

15. Burt, McKen, Burkhart, Hormell, & Knight. (2020). Black Men in Engineering Graduate Education: Experiencing Racial Microaggressions within the Advisor–Advisee Relationship. The Journal of Negro education, 88(4), 493. doi:10.7709/jnegroeducation.88.4.0493

16. Byars-Winston, A. M., Branchaw, J., Pfund, C., Leverett, P., & Newton, J. (2015). Culturally diverse undergraduate researchers’ academic outcomes and perceptions of their research mentoring relationships. International journal of science education, 37(15), 2533–2554. doi:10.1080/09500693.2015.1085133

17. Calisi, R. M., & a Working Group of Mothers in Science. (2018). Opinion: How to tackle the childcare-conference conundrum. Proceedings of the National Academy of Sciences of the United States of America, 115(12), 2845–2849. doi:10.1073/pnas.1803153115

18. Caplar, N., Tacchella, S., & Birrer, S. (2017). Quantitative evaluation of gender bias in astronomical publications from citation counts. Nature Astronomy, 1(6), 0141. doi:10.1038/s41550-017-0141

19. Cech, E. A., & Blair-Loy, M. (2019). The changing career trajectories of new parents in STEM. Proceedings of the National Academy of Sciences of the United States of America, 116(10), 4182–4187. doi:10.1073/pnas.1810862116

20. Cheryan, S., Ziegler, S. A., Montoya, A. K., & Jiang, L. (2017). Why are some STEM fields more gender balanced than others? Psychological Bulletin, 143(1), 1–35. doi:10.1037/bul0000052

21. Cohen, J. (1988). *Statistical power analysis for the behavioral sciences* (2nd ed.). Hillsdale, N.J: L. Erlbaum Associates. doi:10.1016/C2013-0-10517-X

22. Cummins, H. A. (2005). Mommy tracking single women in academia when they are not mommies. Women’s studies international forum, 28(2-3), 222–231. doi:10.1016/j.wsif.2005.04.009

23. David, K. K., Fang, H. Y., Peng, G. C. Y., & Gnadt, J. W. (2020). NIH BRAIN circuits programs: an experiment in supporting team neuroscience. Neuron, 108(6), 1020–1024. doi:10.1016/j.neuron.2020.11.020

24. de Valero, Y. F. (2001). Departmental Factors Affecting Time-to-Degree and Completion Rates of Doctoral Students at One Land-Grant Research Institution. The Journal of higher education, 72(3), 341. doi:10.2307/2649335

25. Dworkin, J. D., Linn, K. A., Teich, E. G., Zurn, P., Shinohara, R. T., & Bassett, D. S. (2020). The extent and drivers of gender imbalance in neuroscience reference lists. Nature Neuroscience, 23(8), 918– 926. doi:10.1038/s41593-020-0658-y

26. Dzirasa, K. (2020). For black scientists, the sorrow is also personal. Cell, 182(2), 263–264. doi:10.1016/j.cell.2020.06.028

27. Ellis, E. M. (2000). Diversity in Higher Education: Race, Gender and The Graduate Student Experience: Recent Research. Retrieved September 14, 2020, from http://web.archive.org/web/20160620200254/www.diversityweb.org/Digest/F00/graduate.htm l

28. Estrada, M., Hernandez, P. R., & Schultz, P. W. (2018). A Longitudinal Study of How Quality Mentorship and Research Experience Integrate Underrepresented Minorities into STEM Careers. CBE life sciences education, 17(1), ar9. doi:10.1187/cbe.17-04-0066

29. Estrada, M., Woodcock, A., Hernandez, P. R., & Schultz, P. W. (2011). Toward a Model of Social Influence that Explains Minority Student Integration into the Scientific Community. Journal of educational psychology, 103(1), 206–222. doi:10.1037/a0020743

30. Estrada, M., Zhi, Q., Nwankwo, E., & Gershon, R. (2019). The influence of social supports on graduate student persistence in biomedical fields. CBE life sciences education, 18(3), ar39. doi:10.1187/cbe.19-01-0029

31. FASEB Office of Public Affairs. (2020). ‘NIH Appropriations’ & Grant Trends: FY 1995-2018. Retrieved from https://www.faseb.org/Portals/2/PDFs/opa/2019/NIH%20Funding%20Trends%20Slide%20Deck%202019%20Update.pdf

32. Fisher, A. J., Mendoza-Denton, R., Patt, C., Young, I., Eppig, A., Garrell, R. L., … Richards, M. A. (2019). Structure and belonging: Pathways to success for underrepresented minority and women PhD students in STEM fields. Plos One, 14(1), e0209279. doi:10.1371/journal.pone.0209279

33. Fuhrmann, C. N., Halme, D. G., O’Sullivan, P. S., & Lindstaedt, B. (2011). Improving graduate education to support a branching career pipeline: recommendations based on a survey of doctoral students in the basic biomedical sciences. CBE life sciences education, 10(3), 239–249. doi:10.1187/cbe.11-02-0013

34. Gardner, S. K. (2008). Fitting the mold of graduate school: A qualitative study of socialization in doctoral education. Innovative Higher Education, 33(2), 125–138. doi:10.1007/s10755-008-9068-x

35. Gazley, J. L., Remich, R., Naffziger-Hirsch, M. E., Keller, J., Campbell, P. B., & McGee, R. (2014). Beyond preparation: identity, cultural capital, and readiness for graduate school in the biomedical sciences. Journal of Research in Science Teaching, 51(8), 1021–1048. doi:10.1002/tea.21164

36. Gentleman, R., Carey, V. J., Huber, W., Irizarry, R. A., & Dudoit, S. (Eds.). (2005). *Bioinformatics and Computational Biology Solutions Using R and Bioconductor*. New York, NY: Springer New York. doi:10.1007/0-387-29362-0

37. Gewin, V. (2020). The career cost of COVID-19 to female researchers, and how science should respond. Nature, 583(7818), 867–869. doi:10.1038/d41586-020-02183-x

38. Gibbs, K. D., Basson, J., Xierali, I. M., & Broniatowski, D. A. (2016). Decoupling of the minority PhD talent pool and assistant professor hiring in medical school basic science departments in the US. eLife, 5. doi:10.7554/eLife.21393

39. Gibbs, K. D., & Griffin, K. A. (2013). What do I want to be with my PhD? The roles of personal values and structural dynamics in shaping the career interests of recent biomedical science PhD graduates. CBE life sciences education, 12(4), 711–723. doi:10.1187/cbe.13-02-0021

40. Gibbs, K. D., McGready, J., Bennett, J. C., & Griffin, K. (2014). Biomedical science ph.d. career interest patterns by race/ethnicity and gender. Plos One, 9(12), e114736. doi:10.1371/journal.pone.0114736

41. Gibbs, K. D., McGready, J., & Griffin, K. (2015). Career Development among American Biomedical Postdocs. CBE life sciences education, 14(4), ar44. doi:10.1187/cbe.15-03-0075

42. Glynn, S. J. (2018, May 18). An Unequal Division of Labor. Retrieved September 13, 2020, from https://www.americanprogress.org/issues/women/reports/2018/05/18/450972/unequal-division-labor/

43. Golde, C., & Dore, T. M. (2004). The Survey of Doctoral Education and Career Preparation: The Importance of Disciplinary Contexts. In D. H. Wulff & A. E. Austin (eds.), Paths to the Professoriate: Strategies for Enriching the Preparation of Future Faculty. San Francisco: Jossey-Bass.

44. Golde, C. M., & Dore, T. M. (2001). ‘At Cross Purposes:’ What the experiences of doctoral students reveal about doctoral education. Philadelphia, PA: A report prepared for The Pew Charitable Trusts.

45. Good, C., Rattan, A., & Dweck, C. S. (2012). Why do women opt out? Sense of belonging and women’s representation in mathematics. Journal of Personality and Social Psychology, 102(4), 700–717. doi:10.1037/a0026659

46. Goulden, M., Mason, M. A., & Frasch, K. (2011). Keeping women in the science pipeline. The Annals of the American Academy of Political and Social Science, 638(1), 141–162. doi:10.1177/0002716211416925

47. Griffin, K. (2018, April 23). Addressing STEM Culture and Climate to Increase Diversity in STEM Disciplines. Higher Education Today.

48. Griffin, K. A. (2016, October 2). Reconsidering the Pipeline Problem: Increasing Faculty Diversity. Higher Education Today. Retrieved from https://www.higheredtoday.org/2016/02/10/reconsidering-the-pipeline-problem-increasing-faculty-diversity/

49. Griffin, K., Gibbs, Jr., K. D., Bennett, J., Staples, C., & Robinson, T. (2015). Respect me for my science“: A Bourdieuian analysis of women scientists” interactions with faculty and socialization into science. Journal of women and minorities in science and engineering, 21(2), 159–179. doi:10.1615/JWomenMinorScienEng.2015011143

50. Guy, B., & Boards, A. (2019). A seat at the table: Exploring the experiences of underrepresented minority women in STEM graduate programs. Journal of prevention & intervention in the community, 47(4), 354–365. doi:10.1080/10852352.2019.1617383

51. Hayter, C. S., & Parker, M. A. (2018). Factors that influence the transition of university postdocs to non-academic scientific careers: An exploratory study. Research Policy, 48(3), 556–570. doi:10.1016/j.respol.2018.09.009

52. Herzig, A H. (2004). Becoming mathematicians: women and students of color choosing and leaving doctoral mathematics. Review of Educational Research, 74(2), 171–214. doi:10.3102/00346543074002171

53. Herzig, Abbe H. (2004). Slaughtering this beautiful math‘: graduate women choosing and leaving mathematics. Gender and education, 16(3), 379–395. doi:10.1080/09540250042000251506

54. Institute of Medicine (US) Council on Health Care Technology, & Goodman, C. (1988). IMPAC (Information for Management Planning Analysis and Coordination).

55. Jacoby, W. G. (1999). Levels of measurement and political research: an optimistic view. American journal of political science, 43(1), 271. doi:10.2307/2991794

56. Jacoby, W. G. (2012). opscale : A Function for Optimal Scaling. Retrieved from https://web.archive.org/web/20180827212513/http://polisci.msu.edu/jacoby/software/optiscale/Jacoby,%20opscale%20MS,%203-26-12.pdf

57. Jones-London, M. (2020). NINDS strategies for enhancing the diversity of neuroscience researchers. Neuron, 107(2), 212–214. doi:10.1016/j.neuron.2020.06.033

58. Kachchaf, R., Ko, L., Hodari, A., & Ong, M. (2015). Career–life balance for women of color: Experiences in science and engineering academia. Journal of diversity in higher education, 8(3), 175–191. doi:10.1037/a0039068

59. Karanja, F., Bostic, R. R., DaCrema, D., & Pacheco, G. G. (2020). Starting Conversations toward Inclusion. Cell, 183(3), 583–586. doi:10.1016/j.cell.2020.09.003

60. Kramer, J. (2020, August 12). Women in Science May Suffer Lasting Career Damage from COVID-19. Scientific American.

61. Lambert, W. M., Wells, M. T., Cipriano, M. F., Sneva, J. N., Morris, J. A., & Golightly, L. M. (2020). Career choices of underrepresented and female postdocs in the biomedical sciences. eLife, 9. doi:10.7554/eLife.48774

62. Larivière, V., Ni, C., Gingras, Y., Cronin, B., & Sugimoto, C. R. (2013). Bibliometrics: global gender disparities in science. Nature, 504(7479), 211–213. doi:10.1038/504211a

63. Larson, R. C., Ghaffarzadegan, N., & Xue, Y. (2014). Too many phd graduates or too few academic job openings: the basic reproductive numberr0in academia. Systems research and behavioral science, 31(6), 745–750. doi:10.1002/sres.2210

64. Lauer, M. (2021, March 2). Announcement of Childcare Costs for Ruth L. Kirschstein National Research Service Award (NRSA) Supported Individual Fellows. Retrieved March 8, 2021, from https://nexus.od.nih.gov/all/2021/03/02/announcement-of-childcare-costs-for-ruth-l-kirschstein-national-research-service-award-nrsa-supported-individual-fellows/

65. Layton, R. L., Brandt, P. D., Freeman, A. M., Harrell, J. R., Hall, J. D., & Sinche, M. (2016). Diversity Exiting the Academy: Influential Factors for the Career Choice of Well-Represented and Underrepresented Minority Scientists. CBE life sciences education, 15(3). doi:10.1187/cbe.16-01- 0066

66. Lenth, R. (2020). emmeans: Estimated Marginal Means, akaLeast-Squares Means (R package version 1.4.5). Computer software, N/A. Retrieved from https://CRAN.R-project.org/package=emmeans

67. Lenzi, R. N., Korn, S. J., Wallace, M., Desmond, N. L., & Labosky, P. A. (2020). The NIH “BEST” programs: Institutional programs, the program evaluation, and early data. The FASEB Journal, 34(3), 3570– 3582. doi:10.1096/fj.201902064

68. Lorden, J. F., Kuh, C. V., & Voytuk, J. A. (2011). Neuroscience and Neurobiology: Combining Data from the Program and Student Surveys. National Academies Press (US).

69. Lovitts, B. E. (2001). Leaving the Ivory Tower: The Causes and Consequences of Departure from Doctoral Study. Rowman & Littlefield Publishers, Inc., 4720 Boston Way, Lanham, MD 20706 (paperback: ISBN-0-7425-0942-9, $29.95; hardcover: ISBN-0-7425-0941-9, $75). Tel: 800-462-6420 (Toll Free); Fax: 301-459-2118; Web site: http://www.rowmanlittlefield.com.

70. Malisch, J. L., Harris, B. N., Sherrer, S. M., Lewis, K. A., Shepherd, S. L., McCarthy, P. C., … Deitloff, J. (2020). Opinion: In the wake of COVID-19, academia needs new solutions to ensure gender equity. Proceedings of the National Academy of Sciences of the United States of America, 117(27), 15378–15381. doi:10.1073/pnas.2010636117

71. Malley, J., Churchwell, J., & Stewart, A. (2006). Assessing the climate for doctoral students at the University of Michigan. Ann Arbor, MI: UM ADVANCE Project, University of Michigan. Retrieved from http://www.umich.edu/~advproj/PhD_Report.pdf

72. Margherio, C., Horner-Devine, M. C., Mizumori, S. J. Y., & Yen, J. W. (2016). Learning to Thrive: Building Diverse Scientists’ Access to Community and Resources through the BRAINS Program. CBE life sciences education, 15(3). doi:10.1187/cbe.16-01-0058

73. Martinez, E. D., Botos, J., Dohoney, K. M., Geiman, T. M., Kolla, S. S., Olivera, A., … Cohen-Fix, O. (2007). Falling off the academic bandwagon. Women are more likely to quit at the postdoc to principal investigator transition. EMBO Reports, 8(11), 977–981. doi:10.1038/sj.embor.7401110

74. Martinez, L. R., O’Brien, K. R., & Hebl, M. R. (2017). Fleeing the ivory tower: gender differences in the turnover experiences of women faculty. Journal of Women’s Health, 26(5), 580–586. doi:10.1089/jwh.2016.6023

75. McGee, R., Saran, S., & Krulwich, T. A. (2012). Diversity in the biomedical research workforce: developing talent. The Mount Sinai Journal of Medicine, New York, 79(3), 397–411. doi:10.1002/msj.21310

76. Mendoza-Denton, R., Patt, C., Fisher, A., Eppig, A., Young, I., Smith, A., & Richards, M. A. (2017). Differences in STEM doctoral publication by ethnicity, gender and academic field at a large public research university. Plos One, 12(4), e0174296. doi:10.1371/journal.pone.0174296

77. Meyers, F. J., Mathur, A., Fuhrmann, C. N., O’Brien, T. C., Wefes, I., Labosky, P. A., … Chalkley, R. (2016). The origin and implementation of the Broadening Experiences in Scientific Training programs: an NIH common fund initiative. The FASEB Journal, 30(2), 507–514. doi:10.1096/fj.15-276139

78. Miller, D. I., & Wai, J. (2015). The bachelor’s to Ph.D. STEM pipeline no longer leaks more women than men: a 30-year analysis. Frontiers in Psychology, 6, 37. doi:10.3389/fpsyg.2015.00037

79. Morgan, A. C., Way, S. F., Hoefer, M. J. D., Larremore, D. B., Galesic, M., & Clauset, A. (2021). The unequal impact of parenthood in academia. Science Advances, 7(9). doi:10.1126/sciadv.abd1996

80. Myers, K. R., Tham, W. Y., Yin, Y., Cohodes, N., Thursby, J. G., Thursby, M. C., … Wang, D. (2020). Unequal effects of the COVID-19 pandemic on scientists. Nature human behaviour, 4(9), 880– 883. doi:10.1038/s41562-020-0921-y

81. National Academies of Sciences, Engineering, and Medicine. (2021). Investigating the Potential Impact of COVID-19 on the Careers of Women in Academic Science, Engineering, and Medicine. Washington, DC: The National Academies Press.

82. National Academy of Sciences (US). (2011). Expanding underrepresented minority participation. Washington (DC): National Academies Press (US). doi:10.17226/12984

83. National Institutes of Health. (2012). Biomedical Research Workforce Working Group Report. National Institutes of Health.

84. National Institutes of Health. (2018, September 24). NOT-OD-18-235: Update on NIH Extension Policy for Early Stage Investigator Status (ESI). Retrieved December 21, 2020, from https://grants.nih.gov/grants/guide/notice-files/NOT-OD-18-235.html

85. National Institutes of Health. (2019a, October 30). NOT-OD-20-011: NIH Extension Policy for Eligibility Window for Pathway to Independence Awards (K99/R00). Retrieved December 20, 2020, from https://grants.nih.gov/grants/guide/notice-files/NOT-OD-20-011.html

86. National Institutes of Health. (2019b, November 22). NOT-OD-20-031: Notice of NIH’s Interest in Diversity. Retrieved November 18, 2020, from https://grants.nih.gov/grants/guide/notice-files/NOT-OD-20-031.html

87. National Institutes of Health. (2020a, January 2). NOT-OD-20-054: Notice of Special Interest: Administrative Supplements to Promote Research Continuity and Retention of NIH Mentored Career Development (K) Award Recipients and Scholars. Retrieved December 21, 2020, from https://grants.nih.gov/grants/guide/notice-files/not-od-20-054.html

88. National Institutes of Health. (2020b, January 17). NOT-OD-20-055: Notice of Special Interest (NOSI): Administrative Supplement for Continuity of Biomedical and Behavioral Research Among First-Time Recipients of NIH Research Project Grant Awards. Retrieved December 20, 2020, from https://grants.nih.gov/grants/guide/notice-files/NOT-OD-20-055.html

89. National Postdoctoral Association. (n.d.). Core Competencies. Retrieved November 19, 2020, from https://www.nationalpostdoc.org/page/CoreCompetencies

90. National Research Council. (2013). *Seeking solutions: Maximizing American talent by advancing women of color in academia: Summary of a conference*. Washington, D.C.: National Academies Press. doi:10.17226/18556

91. National Science Board. (2016). Science and Engineering Indicators 2016 (No. NSB-2016-1). Arlington, VA: National Science Foundation.

92. National Science Foundation. (2015). Women, Minorities, and Persons with Disabilities in Science and Engineering: 2015 (No. Special Report NSF 15-311). Arlington, VA: National Science Foundation, National Center for Science and Engineering Statistics.

93. National Science Foundation. (2020). Scientists and Engineers Statistical Data System (SESTAT). Retrieved February 11, 2021, from https://ncsesdata.nsf.gov/sestat/

94. NIH Data Book. (n.d.). National Statistics on Graduate Students. Retrieved February 28, 2021, from https://report.nih.gov/nihdatabook/report/268

95. NINDS. (2021, February 1). NOT-NS-21-021: Request for Information on the 2021-2026 National Institute of Neurological Disorders and Stroke Strategic Plan. Retrieved March 9, 2021, from https://grants.nih.gov/grants/guide/notice-files/NOT-NS-21-021.html

96. Odekunle, E. A. (2020). To see a face like mine. Cell, 183(3), 564–567. doi:10.1016/j.cell.2020.10.009

97. Ong, M., Wright, C., Espinosa, L., & Orfield, G. (2011). Inside the double bind: A synthesis of empirical research on undergraduate and graduate women of color in science, technology, engineering, and mathematics. Harvard educational review, 81(2), 172–209. doi:10.17763/haer.81.2.t022245n7X4752v2

98. Palmer, R. T., & Young, E. M. (2008). Determined to Succeed: Salient Factors that Foster Academic Success for Academically Unprepared Black Males at a Black College. *Journal of College Student Retention: Research*, Theory and Practice, 10(4), 465–482. doi:10.2190/CS.10.4.d

99. Pezzoni, M., Mairesse, J., Stephan, P., & Lane, J. (2016). Gender and the publication output of graduate students: A case study. Plos One, 11(1), e0145146. doi:10.1371/journal.pone.0145146

100. R Core Team. (2018). R: A language and environment for statistical computing (3.6.3). Computer software, Vienna, Austria: R Foundation for Statistical Computing. Retrieved from https://www.R-project.org/

101. Revelle, W. (2019). psych Procedures for Psychological, Psychometric, and Personality Research (R package version 1.9.12). Computer software, Evanston, Illinois: Northwestern University. Retrieved from https://CRAN.R-project.org/package=psych

102. Roach, M., & Sauermann, H. (2017). The declining interest in an academic career. Plos One, 12(9), e0184130. doi:10.1371/journal.pone.0184130

103. Sassler, S., Glass, J., Levitte, Y., & Michelmore, K. M. (2017). The missing women in STEM? Assessing gender differentials in the factors associated with transition to first jobs. Social science research, 63, 192–208. doi:10.1016/j.ssresearch.2016.09.014

104. Sauermann, H., & Roach, M. (2012). Science PhD career preferences: levels, changes, and advisor encouragement. Plos One, 7(5), e36307. doi:10.1371/journal.pone.0036307

105. Schiebinger, L., & Gilmartin, S. K. (2010, January 1). Housework Is an Academic Issue. Academe, 96(1).

106. Sekar, K. (2020). National Institutes of Health (NIH) Funding: FY1995-FY2021. Congressional Research Service. Retrieved from https://fas.org/sgp/crs/misc/R43341.pdf

107. Shen, Y. A., Webster, J. M., Shoda, Y., & Fine, I. (2018). Persistent underrepresentation of women’s science in high profile journals. BioRxiv. doi:10.1101/275362

108. Sinche, M., Layton, R. L., Brandt, P. D., O’Connell, A. B., Hall, J. D., Freeman, A. M., … Brennwald, P. J. (2017). An evidence-based evaluation of transferrable skills and job satisfaction for science PhDs. Plos One, 12(9), e0185023. doi:10.1371/journal.pone.0185023

109. Sinche, M. V. (2016). *Next gen phd: A guide to career paths in science*. Cambridge, MA and London, England: Harvard University Press. doi:10.4159/9780674974791

110. Smithson, M., & Shou, Y. (2017). Moderator effects differ on alternative effect-size measures. Behavior research methods, 49(2), 747–757. doi:10.3758/s13428-016-0735-z

111. Stachl, C. N., & Baranger, A. M. (2020). Sense of belonging within the graduate community of a research-focused STEM department: Quantitative assessment using a visual narrative and item response theory. Plos One, 15(5), e0233431. doi:10.1371/journal.pone.0233431

112. SurveyMonkey, Inc. (2015). SurveyMonkey. Retrieved November 17, 2020, from http://www.surveymonkey.com

113. Sweet, D. J. (2021). New at cell press: the inclusion and diversity statement. Cell, 184(1), 1–2. doi:10.1016/j.cell.2020.12.019

114. Tagge, R., Lackland, D. T., & Ovbiagele, B. (2021). The TRANSCENDS program: Rationale and overview. Journal of the Neurological Sciences, 420, 117218. doi:10.1016/j.jns.2020.117218

115. Thomas, K. M., Willis, L. A., & Davis, J. (2007). Mentoring minority graduate students: issues and strategies for institutions, faculty, and students. Equal Opportunities International, 26(3), 178– 192. doi:10.1108/02610150710735471

116. Tulshyan, R., & Burey, J.-A. (2021, February 11). Stop Telling Women They Have Imposter Syndrome. Harvard Business Review.

117. US National Science Foundation. (2016a). 2015 Survey of Doctorate Recipients. Retrieved from https://www.nsf.gov/statistics/srvydoctoratework/surveys/srvydoctoratework_2015.pdf

118. US National Science Foundation. (2016b). 2016 Survey of Earned Doctorates. Retrieved from https://www.nsf.gov/statistics/srvydoctorates/surveys/srvydoctorates_2016.pdf

119. Venables, W. N., & Ripley, B. D. (2002). Modern Applied Statistics with S (Statistics and Computing) (4th ed., p. 510). New York: Springer.

120. Wolfinger, N. H., Mason, M. A., & Goulden, M. (2008). Problems in the pipeline: gender, marriage, and fertility in the ivory tower. The Journal of higher education, 79(4), 388–405. doi:10.1080/00221546.2008.11772108

121. Wood, C. V., Jones, R. F., Remich, R. G., Caliendo, A. E., Langford, N. C., Keller, J. L., … McGee, R. (2020). The National Longitudinal Study of Young Life Scientists: Career differentiation among a diverse group of biomedical PhD students. Plos One, 15(6), e0234259. doi:10.1371/journal.pone.0234259

122. Yen, J. W., Horner-Devine, M. C., Margherio, C., & Mizumori, S. J. Y. (2017). The BRAINS program: transforming career development to advance diversity and equity in neuroscience. Neuron, 94(3), 426–430. doi:10.1016/j.neuron.2017.03.049

123. Yoder, J. B., & Mattheis, A. (2016). Queer in STEM: workplace experiences reported in a national survey of LGBTQA individuals in science, technology, engineering, and mathematics careers. Journal of homosexuality, 63(1), 1–27. doi:10.1080/00918369.2015.1078632

124. Zeileis, A, & Hothorn, T. (2002, December). Diagnostic Checking in Regression Relationships. R News, 2(3), 7–10. Retrieved from https://cran.r-project.org/doc/Rnews/Rnews_2002-3.pdf

125. Zeileis, Achim. (2004). Econometric Computing with HC and HAC Covariance Matrix Estimators. Journal of statistical software, 11(10). doi:10.18637/jss.v011.i10

126. Zhao, C., Golde, C. M., & McCormick, A. C. (2007). More than a signature: how advisor choice and advisor behaviour affect doctoral student satisfaction. Journal of Further and Higher Education, 31(3), 263–281. doi:10.1080/03098770701424983

