## Supporting Information for "Factors that Influence Career Choice Among Different Populations of Neuroscience Trainees"

### SUPPORTING INFORMATION APPENDIX

#### RESULTS

##### Descriptive Statistics

Table S1 presents descriptive statistics for the sample.

Table S2 presents basic descriptive statistics for all dependent and independent variables in the study. Dependent variables are ratings of interest in the different career types on a 4-point scale (No interest, Low interest, Moderate interest, Strong interest) for three time-points: start of Ph.D. program (T1), end of Ph.D. program (T2), and current (T3).

Independent (Explanatory) variables are split between those used to predict interest at the start and end of respondents' Ph.D. programs, which captured issues that were contemporaneous to their participation in their Ph.D. programs, and those used to predict current interest, which captured issues that were post-Ph.D. program. Variables that are the result of factor analysis (see Data Analysis) are indicated.

##### Differences by Gender and UR in Explanatory Variables

Table S3 presents abbreviated results of analyses to investigate possible differences by gender and UR status on the Explanatory variables. A summary of the results for analyses of the 4 dichotomous explanatory variables is presented in Table S3a. Of these, only whether the respondents were in the first generation in their family to graduate from a 4-year college or university had any significant differences, by UR status (odds ratio=2.8331, $p<0.001$, medium)—URs were more likely than WRs to be first in their families to graduate. Results for the multinomial logistic regression analyses on the 2 categorical explanatory variables are presented in Table S3b, which shows a 2-way interaction of gender and UR status for respondents' current positions (odds ratio=14.9449, $p<0.05$) that is discussed in the main text. Finally, several significant differences were found in the ANOVA analyses investigating gender and UR status differences on the continuous variables presented in Table S3c.

Significant findings from Table S3 were followed-up by examining differences either in means or slopes for subsamples defined by the moderating variable in question. These results are organized by moderating variable and are presented in Table S4. The False Discovery Rate (FDR) for these analyses was controlled using Benjamini and Hochberg's (1995) procedure, which was applied to each of the 39 analyses using the "BH" option on the mt.rawp2adjp function of R's Multtest package (Pollard, Dudoit, & van der Laan, 2005).

##### Mean Differences in Career Interest Ratings Over Time

Table S5a presents the results from four separate omnibus repeated measures ANOVAs that were conducted to ascertain whether there were differences in the 4 career interest ratings over Time (within-subjects ordinal independent variable), and whether Gender or UR Status (between-subjects independent variables) were moderators of those differences. All degrees of freedom for follow-ups

(Table S5b) were estimated using the Kenward-Roger method, and p values were adjusted to control for family-wise error, FWE, using Tukey.

**Predicting Change in Interest**

*Preliminary Steps*

The first step in the construction of the regression analyses that predicted change in interest over the course of PhD programs was correlation of all graduate school era explanatory variables with respondents' ratings of their interest in the 4 different career types at the end of their PhD programs (T2). Because the final regressions would all include ratings of interest in the same career at the beginning of respondents' PhD programs (T1) among the independent variables (thus the analysis of change in interest), career interest at T2 was regressed on T1 ratings, and correlations with the independent variables were computed with residuals from those procedures (adjusted outcomes). All correlations are presented in Table S6. The FDR for these analyses was again controlled using Benjamini and Hochberg's (1995) procedure through Multtest, which was applied to each set of analyses (all IVs correlated with one DV). In addition, to prevent spurious findings due to the large sample size, only correlations that account for more than 2% of the variance (2 times the 1% threshold for "small" effect size) are reported and used in the final regressions.

As Table S6 shows, only 4 of the 65 correlations between the graduate school era explanatory variables and the 5 adjusted outcomes were both statistically significant and accounted for more than 2% of the variance. This low number is undoubtedly related to the stringent requirements (FDR and effect size) that were applied to each analysis. All 3 of the variables related to changes in interest in academic research-focused careers related to graduate school experiences, especially advisors. Positive relationships with their PhD advisors (r=.21), feeling that they belonged to their labs' intellectual environment (r=.21), and feeling that career advice from their PhD advisors was helpful (r=.31) all related to positive change in interest in academic research-focused careers. In addition, feeling that career advice from their PhD advisors was helpful related to negative change in interest in science-related non-research careers (r=-.15) and non-science careers (r=-.19). These 5 variables were retained for the final regression models.

The second step in the construction of the regression analyses that predicted change in interest in the different career types over the course of graduate school was to investigate interactions between the graduate school era explanatory variables and gender and UR status. Because computing one large regression with all 13 explanatory variables and their interactions with gender, UR status, and the combination of the two for each dependent variable would present multicollinearity problems, it was decided to take a piecemeal approach and investigate each explanatory variable and their interactions separately from the other explanatory variables, and any significant interactions would then be entered into the final regressions. In each analysis if the three-way interaction (explanatory variable by gender by UR status) was not significant, then the analysis war re-run dropping it, to preserve shared variance for the 2-way interactions. Again, FDR was controlled using Benjamini and Hochberg's (1995) procedure through Multtest, which was applied to each set of analyses. Abbreviated results for this step are presented in Table S7. A total of 6 interactions were still significant predictors of change in career interest across graduate school after adjustment for FDR. Because these findings are preliminary, not from the final regression models, follow-up tests to determine their nature were conducted after the final regressions are computed. All 6 interactions were retained for the final regression models.

Table S10a presents follow-up results for significant interactions in the final regressions reported in Table 1 in the article. All follow-ups were computed using the emmeans package in R package (Lenth, 2020).

**Predicting Current Interest**

*Preliminary Steps*

The first step in the construction of the regression analyses that predicted current interest in the 4 different careers was correlation of all explanatory variables with respondents' ratings of their current interest. All correlations are presented in Table S8. The FDR for these analyses was again controlled using Benjamini and Hochberg's (1995) procedure through Multtest, which was applied to each set of analyses (all IVs correlated with one DV). In addition, again to prevent spurious findings due to the large sample size, only correlations that account for more than 2% of the variance (2 times the threshold for "small" effect size) are reported and used in the final regressions.

As Table S8 shows, 32 of the 195 correlations between all of the explanatory variables and the 4 outcomes were both statistically significant and accounted for more than 2% of the variance. Again, this low number is undoubtedly related to the stringent requirements (FDR and effect size) that were applied to each analysis. Although there are too many results to enumerate for this intermediate step, many of the results were predicting current interest in research-focused academic positions. All significant correlations that met the effect size threshold were retained for the final regression analyses.

The second step in the construction of the regression analyses that predicted current interest in the different career types was to investigate interactions between the explanatory variables and Gender and UR status. As with the previous regressions, each explanatory variable was tested for the three-way interaction (explanatory variable by gender by UR status) first, and if that was not significant, then the analysis was re-run dropping it, to preserve shared variance for the 2-way interactions. Again, FDR was controlled using Benjamini and Hochberg's (1995) procedure through Multtest, which was applied to each set of analyses. Abbreviated results for this step are presented in Table S9. A total of 18 interactions were still significant predictors of current interest after adjustment for FDR. Follow-up tests to determine the nature of each interaction were conducted after the final regressions are computed. All 18 interactions were retained for the final regression models.

Table S10b presents follow-up results for significant interactions in the final regressions reported in Table 2 in the article. All follow-ups were computed using the emmeans package in R package (Lenth, 2020).

### METHODS

***General Notes***

Dichotomous variables were recoded to 0 and 1. Prior to any analyses, all continuous variables were visually checked for outliers by plotting and comparing to similar curves, and any outliers were recoded to the largest/smallest value that fit the visual curve (cap method). Using this method only 4 observations were capped. Continuous predictor variables were centered when analyzing interactions.

***Factor Analyses***

Table S11 presents a summary of the factor analyses, which reduced 32 questions to 17 factor variables and performed prior to any analyses.

For each analysis the number of factors to extract was ascertained using BIC scores computed through the VSS function from the psych package (Revelle, 2019) in R. Then, the fa function (also from the psych package) was used to compute maximum-likelihood solutions, with oblique rotation performed using the “promax” option. A summary of each analysis is presented in Table S11 and will be mentioned briefly below.

Table S11a presents the factor analysis for the 12 questions that asked respondents to choose which aspects of careers or work environments were most important to them. This analysis resulted in 7 factors, with one of 7 questions loading heavily on each factor, and the other 5 questions loading only slightly on any of the factors. The final factors were named: “Autonomy,” “Collaboration,” “Make a difference,” “Varied work,” “Ability to do the job,” “Geographic location,” and “Work/Life balance”, and collectively accounted for 66% of the variance in the source questions.

Table S11b presents the factor analysis for the 12 questions that asked respondents to rate how much the item either increases/d or decreases/d their desire to become a faculty member. This analysis resulted in 4 factors, with every question loading at least 0.42 on one factor. The final factors were named: “Funding, Job market, Promotion”, “Research, Autonomy”, “Teaching, Mentoring”, and “Work/Life balance,” and collectively accounted for 41% of the variance in the source questions.

Table S11c presents the factor analysis for the 3 questions that asked respondents about their relationships with their PhD advisors. This analysis resulted in 1 factor, “Relationship with PhD Advisor,” with every question loading at least 0.84 on the factor. The factor accounted for 76% of the variance in the source questions.

Table S11d presents the factor analysis for the 4 questions that asked respondents about their sense of belonging to different groups during their PhD training. This analysis resulted in 2 factors, with every question loading at least 0.41 on one factor. The final factors were named: “PhD: Belong to social community/Department” and “PhD: Belong to intellectual community/Lab,” and collectively accounted for 73% of the variance in the source questions.

Table S11e presents the factor analysis for the 3 questions that asked respondents about their relationships with their postdoc advisors. This analysis resulted in 1 factor, “Relationship with postdoc Advisor,” with every question loading at least 0.88 on the factor. The factor accounted for 80% of the variance in the source questions.

Table S11f presents the factor analysis for the 4 questions that asked respondents about their sense of belonging to different groups during their postdoc training. This analysis resulted in 2 factors, with every question loading at least 0.46 on one factor. The final factors were named: “Postdoc: Belong to social community/Department” and “Postdoc: Belong to intellectual community/Lab,” and collectively accounted for 77% of the variance in the source questions.

### TABLES

**Table S1. Study sample characteristics.** Basic demographic information about the sample of 1,479 PhD neuroscientists who responded to the survey. UR = underrepresented, WR = well represented.

| Variable | Female | Male | Total |
|---|---|---|---|
| Gender | 793 (54%) | 686 (46%) | 1,479 |

| Variable | WR | UR | Total |
|---|---|---|---|
| UR Status | 1246 (84%) | 233 (16%) | 1,479 |

| Variable | Female WR | Male WR | Female UR | Male UR | Total |
|---|---|---|---|---|---|
| Social Identity | 660 (45%) | 586 (40%) | 133 (9%) | 100 (7%) | 1,479 |

| Variable | No | Yes | Total |
|---|---|---|---|
| Have a disability? | 1,415 (98%) | 36 (3%) | 1,451 |

| Variable | No | Yes | Total |
|---|---|---|---|
| First person or among the first generation to graduate from 4 year college? | 1,125 (77%) | 346 (24%) | 1,471 |

| PhD Field | n (%) |
|---|---|
| Neuroscience | 705 (48%) |
| Cellular/Molecular Biology | 137 (9%) |
| Biological Sciences | 111 (8%) |
| Biochemistry/Chemistry | 84 (6%) |
| Bioengineering | 82 (6%) |
| Psychology | 80 (5%) |
| Pharmacology/Toxicology | 60 (4%) |
| Physiology | 49 (3%) |
| Genetics | 34 (2%) |
| Biostatistics, Epidemiology, Public Health, Clinical Sciences | 29 (2%) |
| Physics | 22 (2%) |
| Microbiology and Immunology | 21 (1%) |
| Engineering or Computer Science | 20 (1%) |
| Pathology | 19 (1%) |
| Kinesiology | 14 (1%) |
| Communications/Language | 6 (%) |
| Bioinformatics | 3 (%) |
| Other | 3 (%) |
| Total | 1,479 |

| **Current Position** | **n (%)** |
|---|---|
| Postdoc | 612 (41%) |
| Academic Faculty/Research | 396 (27%) |
| Science, non-research | 150 (10%) |
| Research, non-academic | 135 (9%) |
| Academic Faculty/Teaching | 101 (7%) |
| Non-science | 60 (4%) |
| Unemployed | 25 (2%) |
| Total | 1,479 |

**Table S2. Descriptive information for all variables.** Descriptive statistics for all dependent and independent variables in the study, by category of variable. Min = minimum value, Max = maximum value, N = number in group, SD = standard deviation.

**Table S2a: Outcome Variables**

| **Dependent Variable**<br>**Career Interest Ratings** | **N** | **Min** | **Max** | **Median** | **Mean** | **SD** |
|---|---|---|---|---|---|---|
| ***(At start of PhD)*** | | | | | | |
| Academic Faculty/Research | 1479 | 1 | 4 | 4 | 3.58 | 0.7000 |
| Academic Faculty/Teaching | 1479 | 1 | 4 | 3 | 2.75 | 0.9200 |
| Non-academic Research | 1479 | 1 | 4 | 2 | 2.50 | 0.9300 |
| Science/Non-research | 1479 | 1 | 4 | 2 | 1.91 | 0.8300 |
| ***(At end of PhD)*** | | | | | | |
| Academic Faculty/Research | 1479 | 1 | 4 | 4 | 3.26 | 0.9800 |
| Academic Faculty/Teaching | 1479 | 1 | 4 | 3 | 2.56 | 0.9800 |
| Non-academic Research | 1479 | 1 | 4 | 3 | 2.69 | 0.9600 |
| Science/Non-research | 1479 | 1 | 4 | 2 | 2.20 | 1.0000 |
| ***(Current)*** | | | | | | |
| Academic Faculty/Research | 1479 | 1 | 4 | 3 | 2.99 | 1.1700 |
| Academic Faculty/Teaching | 1479 | 1 | 4 | 2 | 2.43 | 1.0500 |
| Non-academic Research | 1479 | 1 | 4 | 3 | 2.78 | 1.0200 |
| Science/Non-research | 1479 | 1 | 4 | 2 | 2.41 | 1.0700 |

**Table S2b: Graduate School Era Explanatory Variables**

| **Independent Variable**<br>**Predictors of T1 (Start PhD) and T2 (End PhD) interest** | **N** | **Min** | **Max** | **Median** | **Mean** | **SD** |
|---|---|---|---|---|---|---|
| PhD Advisor relationship (factor) | 1479 | -3.59 | 0.66 | 0.44 | 0.00 | 0.9600 |
| PhD Belonging, department/social (factor) | 1479 | -3.31 | 1.16 | 0.05 | 0.00 | 1.0000 |
| PhD Belonging, lab/intellectual (factor) | 1479 | -4.83 | 0.87 | 0.52 | 0.00 | 1.0000 |
| PhD Faculty support, at institution | 1479 | 1 | 4 | 3.00 | 3.22 | 0.8100 |
| PhD Faculty support, outside of institution | 1479 | 1 | 4 | 3.00 | 2.49 | 0.9200 |
| PhD Advisor career advice | 1479 | 1 | 4 | 3.00 | 3.19 | 0.9600 |
| Years of research prior to PhD program | 1479 | 0 | 16 | 2.00 | 2.61 | 1.8900 |
| Times supported by NIH (pre-PhD) | 1204 | 0 | 3 | 1.00 | 0.63 | 0.6400 |
| Top 50 undergraduate institution | 1472 | Yes: 208 | No: 1264 | No | 0.14 | |

**Table S2c: Postdoc and Later Explanatory Variables**

| Independent Variable<br>Predictors of T3 (Current) interest | N | Min | Max | Median | Mean | SD |
|---|---|---|---|---|---|---|
| Postdoc Advisor relationship (factor) | 1231 | -3.1 | 0.8 | 0.37 | 0.00 | 0.9600 |
| Postdoc Belonging, department/social (factor) | 1231 | -2.36 | 1.53 | -0.04 | 0.00 | 1.0000 |
| Postdoc Belonging, lab/intellectual (factor) | 1231 | -3.73 | 0.96 | 0.41 | 0.00 | 1.0000 |
| Postdoc Faculty support, at institution | 1231 | 1 | 4 | 3.00 | 2.88 | 1.0000 |
| Postdoc Faculty support, outside of institution | 1231 | 1 | 4 | 3.00 | 2.70 | 0.9400 |
| Postdoc Advisor career advice | 1182 | 1 | 3 | 2.00 | 2.26 | 0.7400 |
| Total years of research | 1479 | 4 | 31 | 12.00 | 12.05 | 3.5100 |
| Times supported by NIH (post-PhD) | 1204 | 0 | 3 | 1.00 | 0.70 | 0.6800 |
| Years it took to complete PhD | 1479 | 2 | 10 | 6.00 | 5.56 | 1.0500 |
| Years since completed PhD | 1479 | 0 | 9 | 5.00 | 4.92 | 2.5700 |
| # of postdoc positions | 1478 | 1 | 5 | 2.00 | 2.03 | 0.6400 |
| Total time in postdoctoral training | 1241 | 0 | 9 | 3.00 | 3.31 | 1.8600 |
| First-author publication rate | 1479 | 0 | 7 | 0.33 | 0.41 | 0.3600 |
| (Career Aspects) Autonomy (factor) | 1479 | -1.47 | 1.84 | -0.48 | 0.00 | 1.0000 |
| (Career Aspects) Make a difference (factor) | 1479 | -1.55 | 1.39 | 0.57 | 0.00 | 1.0000 |
| (Career Aspects) Collaboration (factor) | 1479 | -1.16 | 1.81 | -0.57 | 0.00 | 1.0000 |
| (Career Aspects) Varied work (factor) | 1479 | -1.08 | 1.94 | -0.53 | 0.00 | 1.0000 |
| (Career Aspects) Ability to do job (factor) | 1479 | -1.23 | 1.71 | -0.56 | 0.00 | 1.0000 |
| (Career Aspects) Geographic location (factor) | 1479 | -1.23 | 1.68 | -0.57 | 0.00 | 1.0000 |
| (Career Aspects) Work/Life balance (factor) | 1479 | -1.41 | 1.13 | 0.78 | 0.00 | 1.0000 |
| (Features of Academia) Funding, Job market, Promotion (factor) | 1479 | -1.86 | 2.92 | -0.12 | 0.00 | 0.9000 |
| (Features of Academia) Research, Autonomy (factor) | 1479 | -4.16 | 1.36 | 0.20 | 0.00 | 0.8600 |
| (Features of Academia) Teaching, Mentoring (factor) | 1479 | -3.38 | 1.36 | 0.13 | 0.00 | 0.8100 |
| (Features of Academia) Work/Life balance (factor) | 1479 | -1.73 | 2.1 | -0.06 | 0.00 | 0.7200 |
| Confident being independent researcher | 1479 | 1 | 5 | 4.00 | 4.02 | 1.0800 |
| Top 50 doctoral institution | 1476 | Yes: 770 | No: 706 | No | 0.52 | |

**Table S3. Differences in explanatory variables by Gender and UR status.** Abbreviated results of analyses to investigate possible differences by gender and UR status on the Explanatory variables. UR = underrepresented, WR = well represented. Effect size: S = small effect size, M = medium effect size. * = p < 0.05, ** = p < 0.01, *** = p < 0.001.

**Table S3a: Logistic Regressions for Dichotomous Explanatory Variables**

| | **Coefficients from Logistic Regression** (Odds Ratio, Significance, Effect Size) | | |
|---|---|---|---|
| **Dichotomous Explanatory Variable** | **Gender Main Effect** | **UR Main Effect** | **Gender by UR Interaction** |
| Top 50 undergraduate institution | 1.0082 | 0.6273 S | 0.8904 |
| Top 50 doctoral institution | 1.0831 | 0.6726 | 1.3575 |
| Have a disability? | 1.8265 S | 2.5480 M | 0.7204 |
| First generation in family to graduate from 4 yr. college? | 0.9760 | 2.8331 *** M | 0.9313 |

**Table S3b: Multinomial Logistic Regressions for Categorical Explanatory Variables**

| | **Coefficient from Multinomial Logistic Regression** (Likelihood Ratio, Significance, Effect Size) | | |
|---|---|---|---|
| **Categorical Explanatory Variable** | **Gender Main Effect** | **UR Main Effect** | **Gender by UR Interaction** |
| Current position | 0.0000 | 0.0000 | 14.9449 * |
| Field of doctoral degree | 0.0000 | 0.0000 | 9.3872 |

**Table S3c: ANOVAs for Continuous Explanatory Variables**

| | **Coefficient from ANOVA** (Coefficient, Significance, Effect Size) | | |
|---|---|---|---|
| **Continuous Explanatory Variable** | **Gender Main Effect** | **UR Main Effect** | **Gender by UR Interaction** |
| PhD Advisor relationship (factor) | 0.1708 *** S | -0.2097 | 0.2694 |
| PhD Belonging, department/social (factor) | -0.0587 | -0.3533 * | 0.3422 * |
| PhD Belonging, lab/intellectual (factor) | 0.0573 ** | -0.4375 ** | 0.4601 ** |
| PhD Faculty support, at institution | 0.0615 | -0.0902 | -0.0143 |
| PhD Faculty support, outside of institution | -0.0427 | 0.234 *** S | 0.0559 |
| PhD Advisor career advice | 0.1452 ** | -0.0551 | 0.0871 |
| Years of research prior to PhD program | -0.2415 | 0.2129 | 0.1168 |
| Times supported by NIH (pre-PhD) | -0.0026 | -0.1188 | 0.1257 |
| Postdoc Advisor relationship (factor) | 0.1601 * | -0.0801 | -0.011 |
| Postdoc Belonging, department/social (factor) | 0.0213 | -0.1439 | 0.1587 |
| Postdoc Belonging, lab/intellectual (factor) | 0.1379 * | -0.3326 *** S | 0.0739 |

| | | | | | | | |
|---|---|---|---|---|---|---|---|
| Postdoc Faculty support, at institution | 0.0692 | | | 0.1104 | | | -0.1051 |
| Postdoc Faculty support, outside of institution | -0.0424 | | | 0.2102 | * | | -0.0726 |
| Postdoc Advisor career advice | 0.1425 | ** | | 0.0676 | | | -0.0193 |
| Total years of research | 0.0847 | | | -0.1698 | | | 0.7932 |
| Times supported by NIH (post-PhD) | -0.0376 | | | -0.1109 | | | 0.0026 |
| Years it took to complete PhD | -0.0678 | | | 0.2469 | * | | -0.1467 |
| Years since completed PhD | 0.317 | * | | -0.379 | | | 0.4044 |
| # of postdoc positions | -0.0629 | | | -0.0485 | | | 0.2054 |
| Total time in postdoctoral training | 0.1908 | | | -0.3053 | | | -0.0499 |
| First-author publication rate | 0.086 | *** | S | -0.0951 | *** | S | -0.01 |
| (Career Aspects) Autonomy (factor) | 0.206 | *** | S | -0.2741 | ** | | 0.0784 |
| (Career Aspects) Make a difference (factor) | -0.1185 | | | -0.1043 | | | 0.1888 |
| (Career Aspects) Collaboration (factor) | -0.0573 | | | -0.2117 | | | 0.2805 |
| (Career Aspects) Varied work (factor) | -0.0746 | | | -0.0733 | | | 0.2105 |
| (Career Aspects) Ability to do job (factor) | -0.0533 | | | -0.1039 | | | 0.1033 |
| (Career Aspects) Geographic location (factor) | -0.1103 | | | -0.1695 | | | 0.2493 |
| (Career Aspects) Work/Life balance (factor) | -0.2646 | *** | S | 0.0374 | | | -0.0912 |
| (Features of Academia) Funding, Job market, Promotion (factor) | 0.2514 | *** | S | 0.0815 | | | 0.0518 |
| (Features of Academia) Research, Autonomy (factor) | 0.1986 | *** | S | -0.0958 | | | 0.1738 |
| (Features of Academia) Teaching, Mentoring (factor) | -0.0681 | | | 0.0353 | | | 0.1137 |
| (Features of Academia) Work/Life balance (factor) | 0.1333 | *** | | 0.1308 | ** | | 0.0644 |
| Confident being independent researcher | 0.4035 | *** | S | 0.033 | | | 0.1469 |

**Table S4. Follow-ups for Gender and UR Status Differences on Explanatory Variables.** Follow-up analyses were performed on significant findings in Table S3 by examining differences either in means or slopes for subsamples defined by whichever was significant of gender, UR status, or their interaction. UR = underrepresented, WR = well represented. N = number in group, M = mean, n = number in subgroup, SD = standard deviation. Effect size: (-) = negligible effect size, (s) = small effect size. * = p < 0.05, ** = p < 0.01, *** = p < 0.001.

**Table S4a: Contingency table for Gender by Current Position association**

| | Gender | | | | | |
|---|---|---|---|---|---|---|
| **Current Position** (significance) (effect size) | **Female** | **%** | **Male** | **%** | **Total** | **%** |
| Academic Faculty/Research (**) (s) | 179 | 45.2% | 217 | 54.8% | 396 | 100.0% |
| Academic Faculty/Teaching (*) (s) | 65 | 64.4% | 36 | 35.6% | 101 | 100.0% |
| Science/Non-research (*) (s) | 118 | 78.7% | 32 | 21.3% | 150 | 100.0% |

**Table S4b: Means for continuous explanatory variables split by Gender**

| | | | Gender | | | | | | |
|---|---|---|---|---|---|---|---|---|---|
| | Overall | | Female | | Male | | | | |
| **Dependent Variable** | M | N | M | n | M | n | **Mean Diff** | **Pooled SD** | **Cohen's d** |
| PhD Advisor relationship (factor) (***) (s) | 0.00 | 1479 | -0.10 | 793 | 0.12 | 686 | -0.21 | 0.9500 | -0.23 |
| PhD Belonging, lab/intellectual (factor) (**) (-) | 0 | 1479 | -0.06 | 793 | 0.07 | 686 | -0.13 | 1 | -0.135 |
| PhD Advisor career advice (**) (-) | 3.19 | 1479 | 3.11 | 793 | 3.27 | 686 | -0.16 | 0.9600 | -0.17 |
| Postdoc Advisor relationship (factor) (*) (-) | 0.00 | 1231 | -0.07 | 665 | 0.09 | 566 | -0.16 | 0.9600 | -0.17 |
| Postdoc Belonging, lab/intellectual (factor) (*) (-) | 0 | 1231 | -0.07 | 665 | 0.08 | 566 | -0.15 | 1 | -0.153 |
| Postdoc Advisor career advice (**) (-) | 2.26 | 1182 | 2.20 | 636 | 2.34 | 546 | -0.14 | 0.7400 | -0.19 |
| Years since completed PhD (*) (-) | 4.92 | 1479 | 4.74 | 793 | 5.13 | 686 | -0.38 | 2.56 | -0.15 |
| First-author publication rate (***) (s) | 0.41 | 1479 | 0.37 | 793 | 0.45 | 686 | -0.09 | 0.3600 | -0.24 |
| (Career Aspects) Autonomy (factor) (***) (s) | 0.00 | 1479 | -0.10 | 793 | 0.12 | 686 | -0.22 | 0.9900 | -0.23 |
| (Career Aspects) Work/Life balance (factor) (***) (s) | 0.00 | 1479 | 0.13 | 793 | -0.15 | 686 | 0.28 | 0.9900 | 0.28 |
| (Features of Academia) Funding, Job market, Promotion (factor) (***) (s) | 0.00 | 1479 | -0.12 | 793 | 0.14 | 686 | -0.26 | 0.8900 | -0.29 |
| (Features of Academia) Research, Autonomy (factor) (***) (s) | 0.00 | 1479 | -0.10 | 793 | 0.12 | 686 | -0.23 | 0.8600 | -0.26 |
| (Features of Academia) Work/Life balance (factor) (***) (-) | 0.00 | 1479 | -0.06 | 793 | 0.07 | 686 | -0.14 | 0.7100 | -0.20 |
| Confident being independent researcher (***) (s) | 4.02 | 1479 | 3.82 | 793 | 4.25 | 686 | -0.42 | 1.0600 | -0.40 |

**Table S4c: Contingency table for UR Status by Family College Graduation History association**

| UR Status (significance) (effect size) | 1st generation to graduate from 4yr college | | | | Total | % |
|---|---|---|---|---|---|---|
| | No | % | Yes | % | | |
| Well-represented (***) (s) | 989 | 79.8% | 251 | 20.2% | 1240 | 100.0% |
| Under-represented (**) (-) | 136 | 58.9% | 95 | 41.1% | 231 | 100.0% |
| Total | 1125 | 76.5% | 346 | 23.5% | 1471 | 100.0% |

**Table S4d: Means for continuous explanatory variables split by UR Status**

| Dependent Variable | Overall | | UR Status | | | | Mean Diff | Pooled SD | Cohen's d |
|---|---|---|---|---|---|---|---|---|---|
| | | | WR | | UR | | | | |
| | M | n | M | n | M | n | | | |
| PhD Belonging, department/social (factor) (*) (-) | 0 | 1479 | 0.03 | 1246 | -0.17 | 233 | 0.2 | 1 | 0.205 |
| PhD Belonging, lab/intellectual (factor) (**) (-) | 0 | 1479 | 0.04 | 1246 | -0.2 | 233 | 0.24 | 0.99 | 0.238 |
| PhD Faculty support, outside of institution (***) (s) | 2.49 | 1479 | 2.45 | 1246 | 2.71 | 233 | -0.26 | 0.92 | -0.282 |
| Postdoc Belonging, lab/intellectual (factor) (***) (s) | 0 | 1231 | 0.05 | 1034 | -0.25 | 197 | 0.3 | 0.99 | 0.302 |
| Postdoc Faculty support, outside of institution (*) (-) | 2.7 | 1231 | 2.67 | 1034 | 2.85 | 197 | -0.18 | 0.94 | -0.188 |
| Years it took to complete PhD (*) (-) | 5.56 | 1479 | 5.53 | 1246 | 5.71 | 233 | -0.18 | 1.05 | -0.174 |
| First-author publication rate (***) (s) | 0.41 | 1479 | 0.42 | 1246 | 0.32 | 233 | 0.1 | 0.36 | 0.278 |
| (Career Aspects) Autonomy (factor) (**) (-) | 0 | 1479 | 0.04 | 1246 | -0.2 | 233 | 0.24 | 0.99 | 0.241 |
| (Features of Academia) Work/Life balance (factor) (**) (-) | 0 | 1479 | -0.03 | 1246 | 0.13 | 233 | -0.16 | 0.71 | -0.222 |

**Table S4e: Follow-ups for continuous explanatory variables that had significant interactions of Gender by UR Status**

| Dependent Variable / Interaction Context | Overall | | UR Status | | | | Mean Diff | Pooled SD | Cohen's d |
|---|---|---|---|---|---|---|---|---|---|
| | | | WR | | UR | | | | |
| | F | p | M | n | M | n | | | |
| PhD Belonging, lab/intellectual BY UR, within Females (***) (s) | 19.5 | 0 | 0.01 | 660 | -0.43 | 133 | 0.44 | 1.04 | 0.42 |
| PhD Belonging, lab/intellectual BY UR, within Males (-) (-) | 0.05 | 0.8209 | 0.07 | 586 | 0.09 | 100 | -0.02 | 0.92 | -0.025 |
| PhD Belonging, department/social BY UR within Females (***) (s) | 13.74 | 0.0002 | 0.06 | 660 | -0.29 | 133 | 0.35 | 1 | 0.352 |
| PhD Belonging, department/social BY UR within Males (-) (-) | 0.01 | 0.917 | 0 | 586 | -0.01 | 100 | 0.01 | 0.98 | 0.011 |

**Table S5. Omnibus MANOVA Results for Change in Career Interest Ratings Over Time.** Results from four separate omnibus repeated measures ANOVAs to ascertain whether there were differences in the 4 career interest ratings over time (within-subjects ordinal independent variable), and whether Gender or UR Status (between-subjects independent variables) were moderators of those differences. UR = underrepresented, WR = well represented. SD = standard deviation. * = p < 0.05, ** = p < 0.01, *** = p < 0.001.

**Table S5a: Omnibus MANOVA means**

| Dependent Variables: T2 (End of PhD) Career Interest Ratings | Independent Variable: Time | | | | | | | |
|---|---|---|---|---|---|---|---|---|
| | T1 (Start PhD) | | Sig | T2 (End PhD) | | Sig | T3 (Current) | |
| | Mean | SD | T1 vs T2 | Mean | SD | T2 vs T3 | Mean | SD |
| Academic Faculty/Research | 3.58 | 0.0345 | *** | 3.23 | 0.0345 | *** | 2.98 | 0.0345 |
| Academic Faculty/Teaching | 2.74 | 0.0354 | *** | 2.58 | 0.0354 | ** | 2.48 | 0.0354 |
| Non-academic Research | 2.50 | 0.0348 | *** | 2.71 | 0.0348 | ** | 2.82 | 0.0348 |
| Science/Non-research | 1.95 | 0.0344 | *** | 2.31 | 0.0344 | *** | 2.48 | 0.0344 |

**Table S5b: Follow-ups for significant interactions**

| Dependent Variables: T2 (End of PhD) Career Interest Ratings by Interaction Context | Levels of Moderator | Independent Variable: Time | | | | | | | | |
|---|---|---|---|---|---|---|---|---|---|---|
| | | T1 (Start PhD) | | Sig | T2 (End PhD) | | Sig | T3 (Current) | | Sig |
| | | Mean | SD | L1 vs L2 | Mean | SD | L1 vs L2 | Mean | SD | L1 vs L2 |
| Academic Faculty/Research by GENDER by TIME | 1 Female | 3.48 | 0.0455 | *** | 3.08 | 0.0455 | *** | 2.80 | 0.0455 | *** |
| | 2 Male | 3.69 | 0.0518 | | 3.38 | 0.0518 | | 3.15 | 0.0518 | |
| Academic Faculty/Teaching by UR STATUS by TIME | 1 WR | 2.76 | 0.0279 | | 2.55 | 0.0279 | | 2.40 | 0.0279 | * |
| | 2 UR | 2.72 | 0.0651 | | 2.61 | 0.0651 | | 2.55 | 0.0651 | |
| Science/Non-research by GENDER by TIME | 1 Female | 2.06 | 0.0454 | *** | 2.45 | 0.0454 | *** | 2.70 | 0.0454 | *** |
| | 2 Male | 1.83 | 0.0517 | | 2.16 | 0.0517 | | 2.27 | 0.0517 | |

**Table S6. Correlations for T1->T2 Regressions.** Career interest at T2 was regressed on T1 ratings, and correlations with the independent variables were computed with residuals from those procedures (adjusted outcomes). T1 = Time 1 (Start of PhD), T2 = Time 2 (End of PhD). * = p < 0.05, ** = p < 0.01, *** = p < 0.001. Shaded: > 2% variance

| Independent Variable (Graduate School Era Explanatory) | Dependent Variable<br>T2/End of Graduate School Career Interest Ratings<br>(Correlation, Significance) | | | | | | | |
|---|---|---|---|---|---|---|---|---|
| | Academic Faculty/Research | | Academic Faculty/Teaching | | Non-academic Research | | Science/Non-research | |
| PhD Advisor relationship (factor) | 0.2100 | *** | 0.0547 | | -0.0650 | | -0.1200 | ** |
| PhD Belonging, department/social (factor) | 0.0800 | | -0.0055 | | -0.0220 | | -0.0170 | |
| PhD Belonging, lab/intellectual (factor) | 0.2100 | *** | 0.0225 | | -0.0384 | | -0.0900 | |
| PhD Faculty support, at institution | 0.1300 | *** | 0.0198 | | -0.0423 | | -0.0160 | |
| PhD Faculty support, outside of institution | 0.0529 | | 0.0490 | | -0.0501 | | -0.0232 | |
| PhD Advisor career advice | 0.3100 | *** | 0.1200 | ** | -0.1100 | ** | -0.1500 | *** |
| Years of research prior to PhD program | -0.0109 | | 0.0047 | | -0.0309 | | -0.0716 | |
| Top 50 undergraduate institution | 0.0133 | | -0.0297 | | -0.0082 | | -0.0277 | |
| Times supported by NIH (pre-PhD) | 0.0699 | | 0.0405 | | -0.0522 | | 0.0010 | |
| Have a disability? | -0.0802 | | -0.0541 | | 0.0720 | | 0.0155 | |
| First person/generation to graduate from 4yr college? | -0.0139 | | 0.0206 | | 0.0522 | | 0.0565 | |
| Gender | -0.1000 | * | 0.0279 | | -0.0316 | | 0.1300 | *** |
| UR Status | -0.0475 | | 0.0373 | | 0.0372 | | 0.1000 | * |

**Table S7. Interaction Tests for T1->T2 Regressions.** Abbreviated results for investigation of interactions between the graduate school era explanatory variables and gender and UR status. IV = independent variable, T1 = Time 1 (Start of PhD), T2 = Time 2 (End of PhD). * = p < 0.05, ** = p < 0.01, *** = p < 0.001.

| Independent Variable (Graduate School Era Explanatory) | Dependent Variable T2/End of PhD Interest Ratings (Coefficient, Significance) | | | | | | | | | | | |
|---|---|---|---|---|---|---|---|---|---|---|---|---|
| | Academic Faculty/Research | | | Academic Faculty/Teaching | | | Non-academic Research | | | Science/Non-research | | |
| | IV Interaction with … | | | IV Interaction with … | | | IV Interaction with … | | | IV Interaction with … | | |
| | Gender | UR Status | Gender* UR Status | Gender | UR Status | Gender* UR Status | Gender | UR Status | Gender* UR Status | Gender | UR Status | Gender* UR Status |
| T1 Interest | -0.0207 | -0.0079 | 0.0754 | 0.0414 | 0.0214 | 0.0776 | 0.1280 ** | -0.0472 | -0.0610 | -0.0342 | -0.1436 * | 0.1309 |
| PhD Advisor relationship (factor) | -0.0324 | -0.0913 | -0.0689 | 0.0172 | -0.0518 | -0.0830 | -0.0442 | 0.0331 | 0.0126 | 0.0305 | 0.0485 | 0.1232 |
| PhD Belonging, department/social (factor) | -0.0367 | -0.0772 | 0.0655 | -0.0020 | 0.0234 | -0.1257 | -0.0117 | 0.0277 | -0.1800 | -0.0236 | 0.0460 | -0.2272. |
| PhD Belonging, lab/intellectual (factor) | -0.0491 | -0.0370 | 0.0483 | -0.0005 | 0.0043 | -0.0503 | 0.0081 | 0.0145 | -0.1050 | 0.0567 | 0.0771 | -0.1291 |
| Times supported by NIH (pre-PhD) | -0.1061 | -0.1205 | 0.0036 | 0.0746 | 0.0301 | 0.0944 | 0.0338 | 0.0644 | -0.0388 | 0.0704 | -0.1829. | -0.0547 |
| PhD Faculty support, at institution | -0.0632 | -0.0819 | -0.0631 | -0.0530 | -0.0731 | -0.2996 * | 0.0611 | 0.1307 | 0.1200 | 0.1059. | 0.0155 | -0.0932 |
| PhD Faculty support, outside of institution | -0.0301 | -0.0072 | -0.3206 * | 0.0377 | -0.0906 | -0.0883 | 0.0137 | 0.0550 | -0.0170 | 0.0452 | 0.0188 | 0.0326 |
| PhD Advisor career advice | -0.0131 | -0.0878 | -0.1382 | 0.0143 | -0.1152 * | -0.0407 | -0.0484 | 0.0304 | -0.0021 | 0.0287 | 0.0530 | -0.1867 |
| Years of research prior to PhD program | -0.0337 | -0.0200 | 0.0213 | -0.0166 | -0.0306 | 0.0485 | -0.0134 | -0.0537 | 0.0575 | 0.0125 | -0.0506 | -0.0331 |
| Top 50 undergraduate institution | -0.0912 | -0.2088 | -0.3470 | 0.1115 | 0.0927 | -0.4387 | -0.1796 | 0.1007 | 0.2456 | 0.0729 | -0.0244 | -0.1355 |
| Have a disability? | 0.4502 | -0.1048 | 0.1651 | 0.2723 | 0.3085 | 0.1494 | 0.2100 | -0.1842 | -0.6510 | -0.0500 | 0.1373 | 0.6229 |
| First person/ generation to graduate from 4yr college? | -0.0728 | 0.0154 | 0.0232 | 0.0985 | 0.0644 | 0.0034 | -0.1562 | -0.0896 | -0.1299 | 0.0279 | 0.0551 | 0.1903 |

**Table S8. Correlations for T3 Regressions.** Correlation of all explanatory variables with current interest ratings. * = p < 0.05, ** = p < 0.01, *** = p < 0.001. Shaded: > 2% variance.

| | **Dependent Variable T3/Current Career Interest Ratings** (Correlation, Significance) | | | | | | | |
|---|---|---|---|---|---|---|---|---|
| **Independent Variable (All Explanatory)** | **Academic/Research** | | **Academic/Teaching** | | **Non-academic Research** | | **Science/Non-research** | |
| PhD Advisor relationship (factor) | 0.12 | ** | 0.00 | | 0.01 | | -0.07 | |
| PhD Belonging, department/social (factor) | 0.02 | | 0.00 | | -0.02 | | 0.01 | |
| PhD Belonging, lab/intellectual (factor) | 0.14 | *** | -0.02 | | 0.00 | | -0.06 | |
| PhD Faculty support, at institution | 0.13 | *** | 0.05 | | 0.02 | | 0.00 | |
| PhD Faculty support, outside of institution | 0.11 | * | 0.08 | | -0.02 | | -0.02 | |
| PhD Advisor career advice | 0.30 | *** | 0.10 | | -0.07 | | -0.13 | *** |
| Years of research prior to PhD program | -0.01 | | 0.01 | | 0.04 | | -0.02 | |
| Top 50 undergraduate institution | 0.03 | | -0.07 | | 0.01 | | -0.03 | |
| Times supported by NIH (pre-PhD) | 0.04 | | 0.02 | | -0.04 | | -0.03 | |
| Have a disability? | -0.10 | * | -0.06 | | 0.07 | | 0.04 | |
| First person/generation to graduate from 4yr college? | -0.03 | | 0.03 | | 0.02 | | 0.06 | |
| Gender | -0.15 | *** | 0.02 | | -0.08 | | 0.20 | *** |
| UR Status | -0.03 | | 0.05 | | 0.03 | | 0.09 | |
| Postdoc Advisor relationship (factor) | 0.26 | *** | 0.07 | | 0.01 | | -0.14 | *** |
| Postdoc Belonging, department/social (factor) | 0.14 | *** | 0.04 | | -0.03 | | -0.11 | * |
| Postdoc Belonging, lab/intellectual (factor) | 0.21 | *** | 0.04 | | 0.00 | | -0.15 | *** |
| Postdoc Faculty support, at institution | 0.20 | *** | 0.06 | | -0.07 | | -0.11 | * |
| Postdoc Faculty support, outside of institution | 0.16 | *** | 0.04 | | -0.10 | | -0.05 | |
| Postdoc Advisor career advice | 0.35 | *** | 0.11 | * | -0.07 | | -0.15 | *** |
| Total years of research | 0.14 | *** | -0.01 | | 0.07 | | -0.09 | |
| Top 50 doctoral institution | 0.01 | | -0.06 | | 0.00 | | -0.05 | |
| Years it took to complete PhD | -0.15 | *** | -0.02 | | 0.04 | | 0.12 | ** |
| Years since completed PhD | 0.11 | * | 0.02 | | -0.09 | | -0.09 | |
| # of postdoc positions | 0.13 | *** | 0.05 | | 0.04 | | -0.03 | |
| Total time in postdoctoral training | 0.12 | * | 0.03 | | -0.06 | | -0.09 | |
| First-author publication rate | 0.22 | *** | 0.00 | | -0.04 | | -0.14 | *** |
| Times supported by NIH (post-PhD) | 0.03 | | 0.10 | | -0.03 | | 0.04 | |
| (Career Aspects) Autonomy (factor) | 0.30 | *** | 0.03 | | -0.12 | ** | -0.15 | *** |
| (Career Aspects) Make a difference (factor) | 0.08 | | 0.09 | | -0.03 | | 0.01 | |
| (Career Aspects) Collaboration (factor) | 0.17 | *** | -0.01 | | -0.07 | | -0.05 | |
| (Career Aspects) Varied work (factor) | 0.04 | | 0.01 | | -0.05 | | 0.00 | |
| (Career Aspects) Ability to do job (factor) | 0.14 | *** | 0.10 | * | -0.11 | * | -0.05 | |
| (Career Aspects) Geographic location (factor) | 0.01 | | 0.06 | | 0.04 | | -0.01 | |
| (Career Aspects) Work/Life balance (factor) | -0.23 | *** | 0.05 | | 0.07 | | 0.15 | *** |
| (Features of Academia) Funding, Job market, Promotion (factor) | 0.53 | *** | 0.19 | *** | -0.18 | *** | -0.27 | *** |
| (Features of Academia) Research, Autonomy (factor) | 0.60 | *** | 0.15 | *** | -0.04 | | -0.26 | *** |
| (Features of Academia) Teaching, Mentoring (factor) | 0.21 | *** | 0.49 | *** | -0.21 | *** | 0.02 | |
| (Features of Academia) Work/Life balance (factor) | 0.38 | *** | 0.32 | *** | -0.13 | *** | -0.16 | *** |
| Confident being independent researcher | 0.32 | *** | -0.02 | | -0.02 | | -0.22 | *** |

**Table S9. Interaction Tests for T3 Regressions.** Abbreviated results for investigation of interactions between the explanatory variables and Gender and UR status * = p < 0.05, ** = p < 0.01, *** = p < 0.001.

| | Dependent Variable T2/End of Graduate School Career Interest Ratings (Coefficient, Significance) | | | | | | | | | | | |
|---|---|---|---|---|---|---|---|---|---|---|---|---|
| | Academic Faculty/Research | | | Academic Faculty/Teaching | | | Non-academic Research | | | Science/Non-research | | |
| | IV Interaction with ... | | | IV Interaction with ... | | | IV Interaction with ... | | | IV Interaction with ... | | |
| **Independent Variable (Explanatory)** | **Gender** | **UR Status** | **Gender*UR Status** | **Gender** | **UR Status** | **Gender*UR Status** | **Gender** | **UR Status** | **Gender*UR Status** | **Gender** | **UR Status** | **Gender*UR Status** |
| PhD Advisor relationship (factor) | -0.0523 | -0.0602 | -0.1831 | 0.0167 | -0.0488 | -0.0732 | -0.0391 | 0.0709 | -0.2921 | -0.0279 | 0.0065 | -0.1487 |
| PhD Belonging, department/social (factor) | 0.016 | -0.0331 | -0.0433 | 0.1005 | 0.0134 | -0.0931 | -0.0064 | 0.0333 | -0.0207 | -0.0107 | 0.0188 | 0.0014 |
| PhD Belonging, lab/intellectual (factor) | -0.0384 | -0.0094 | 0.0407 | 0.0283 | -0.038 | -0.0078 | 0.0059 | 0.0399 | -0.2256 | 0.0629 | 0.0108 | -0.107 |
| PhD Faculty support, at institution | 0.0526 | -0.1066 | -0.1465 | 0.0682 | -0.0804 | -0.2887 | 0.0523 | 0.1867 . | -0.0392 | 0.1261 | 0.0229 | 0.0215 |
| PhD Faculty support, outside of institution | -0.0563 | -0.1238 | -0.1751 | 0.0428 | -0.0647 | 0.0343 | -0.0556 | -0.0029 | -0.1402 | 0.0474 | 0.0326 | 0.0937 |
| PhD Advisor career advice | 0.045 | -0.1437 | -0.1276 | 0.0424 | -0.1849 * | 0.0449 | -0.0709 | 0.0189 | -0.0833 | 0.0237 | -0.133 . | -0.1773 |
| Years of research prior to PhD program | -0.0545 | -0.0478 | 0.0273 | 0.0016 | -0.0145 | 0.0256 | -0.0162 | -0.0268 | 0.1248 | 0.0615 . | 0.0764 . | 0.0964 |
| Undergraduate institution in Top 50 | -0.1235 | -0.6032 . | 0.0113 | 0.3055 | -0.1971 | 0.4236 | -0.1384 | 0.4487 | 0.2229 | 0.0265 | 0.0774 | -1.2763 * |
| Times supported by NIH (pre-PhD) | -0.1145 | -0.1228 | -0.3564 | 0.0089 | 0.0566 | -0.0663 | -0.0158 | 0.1819 | -0.2983 | 0.0056 | -0.1184 | -0.1814 |
| Have a disability? | 0.5554 | -0.0553 | -0.4893 | 0.0526 | 0.3642 | -0.3194 | 0.1122 | -0.0638 | 0.0138 | -0.4714 | 0.1144 | 0.7524 |
| First person/gen to graduate from 4yr college? | -0.1651 | 0.1293 | 0.0145 | 0.0056 | 0.2049 | 0.1764 | -0.1567 | -0.1442 | -0.5518 | 0.1516 | 0.1059 | 0.3035 |
| Postdoc Advisor relationship (factor) | -0.041 | 0.0443 | -0.2107 | -0.0202 | 0.0719 | -0.0913 | -0.0157 | -0.1371 | -0.0281 | 0.0466 | -0.0239 | 0.0706 |
| Postdoc Belonging, department/social (factor) | 0.0697 | 0.0414 | -0.1254 | 0.0016 | -0.0068 | 0.0569 | 0.0143 | 0.1366 | 0.0613 | -0.0551 | -0.0688 | 0.1809 |
| Postdoc Belonging, lab/intellectual (factor) | -0.0254 | 0.0598 | -0.0837 | -0.0718 | -0.0059 | 0.0149 | -0.0154 | 0.0421 | -0.2371 | 0.0042 | -0.0327 | 0.0091 |
| Postdoc Faculty support, at institution | -0.1181 . | -0.0519 | -0.1924 | 0.0487 | 0.006 | -0.2375 | 0.0132 | 0.0788 | 0.2314 | 0.0956 | -0.0141 | 0.0928 |
| Postdoc Faculty support, outside of institution | -0.0745 | -0.0442 | 0.0928 | 0.02 | 0.0317 | 0.0956 | 0.0212 | -0.0917 | 0.053 | 0.0293 | 0.0917 | -0.182 |
| Postdoc Advisor career advice | 0.0144 | -0.0897 | 0.105 | -0.0765 | -0.0347 | 0.0808 | -0.0744 | 0.0681 | -0.3674 | 0.0684 | 0.0096 | -0.0015 |
| Total years of research | -0.023 | -0.0084 | 0.0033 | 0.0227 | 0.0314 | 0.0288 | -0.0164 | -0.0195 | 0.043 | 0.0155 | 0.0183 | 0.058 |
| Top 50 doctoral institution | 0.1103 | -0.4523 * | -0.3617 | -0.0535 | -0.2493 | -0.255 | 0.0428 | 0.0198 | 0.3349 | -0.0813 | 0.1385 | -0.3963 |
| Years it took to complete PhD | 0.0611 | -0.0671 | 0.1005 | 0.0514 | -0.0417 | 0.0379 | 0.0769 | -0.0465 | -0.0701 | -0.1367 ** | 0.0458 | 0.0764 |
| Years since completed PhD | 0.0014 | -0.0069 | 0.0104 | 0.0193 | 0.0383 | 0.0548 | 0.002 | 0.004 | 0.0699 | -0.0212 | 0.009 | 0.0327 |
| # of postdoc positions | -0.0613 | 0.1307 | 0.2268 | 0.0217 | 0.2015 | 0.0364 | -0.0235 | -0.1305 | 0.0114 | 0.1373 | 0.0079 | 0.1794 |
| Total time in postdoctoral training | -0.0128 | 0.0246 | -0.0015 | 0.0548 | 0.0808 | 0.0859 | -0.0436 | -0.0068 | -0.0242 | -0.0273 | -0.0247 | 0.0296 |
| First-author publication rate | -0.6141 *** | 0.6982 * | 0.3673 | 0.0094 | -0.1624 | 0.2097 | -0.1361 | -0.3559 | -0.2577 | 0.4352 * | -0.8002 * | 0.0854 |
| Times supported by NIH (post-PhD) | 0.0849 | 0.0484 | 0.294 | 0.0452 | -0.1083 | -0.1654 | 0.0192 | -0.0127 | -0.1495 | 0.0582 | -0.03 | -0.1218 |

**Table S9 (cont.)**

| Independent Variable (Explanatory) | Dependent Variable T2/End of Graduate School Career Interest Ratings (Coefficient, Significance) | | | | | | | | | | | |
|---|---|---|---|---|---|---|---|---|---|---|---|---|
| | Academic Faculty/Research | | | Academic Faculty/Teaching | | | Non-academic Research | | | Science/Non-research | | |
| | IV Interaction with … | | | IV Interaction with … | | | IV Interaction with … | | | IV Interaction with … | | |
| | Gender | UR Status | Gender*UR Status | Gender | UR Status | Gender*UR Status | Gender | UR Status | Gender*UR Status | Gender | UR Status | Gender*UR Status |
| (Career Aspects) Autonomy (factor) | 0.0169 | -0.0032 | -0.2021 | 0.0126 | 0.0394 | -0.0752 | 0.0001 | 0.0052 | 0.0004 | 0.0902 | 0.0674 | 0.0169 |
| (Career Aspects) Make a difference (factor) | 0.028 | 0.0619 | -0.0602 | 0.0177 | -0.0359 | -0.3067 | 0.0881 | -0.0111 | 0.105 | 0.023 | 0.1119 | -0.0364 |
| (Career Aspects) Collaboration (factor) | 0.0221 | -0.0614 | 0.0315 | 0.0842 | 0.0649 | -0.2927 | -0.0018 | -0.0685 | -0.1411 | 0.0587 | 0.1379 . | -0.0297 |
| (Career Aspects) Varied work (factor) | -0.0215 | -0.0805 | -0.1779 | 0.0532 | -0.0109 | -0.1835 | 0.0227 | 0.0246 | -0.055 | 0.0479 | 0.0391 | -0.1955 |
| (Career Aspects) Ability to do job (factor) | 0.0308 | -0.059 | 0.1067 | 0.0816 | 0.0396 | -0.2017 | -0.0377 | 0.0072 | -0.2737 | 0.0391 | 0.1139 | -0.1189 |
| (Career Aspects) Geographic location (factor) | -0.0555 | 0.1692 . | -0.0539 | -0.0018 | -0.0075 | -0.164 | 0.0123 | -0.0632 | -0.2379 | -0.0465 | -0.1243 | -0.1661 |
| (Career Aspects) Work/Life balance (factor) | 0.0822 | 0.0887 | -0.022 | -0.01 | 0.0165 | 0.0834 | 0.0974 | 0.0192 | -0.0304 | -0.0922 | -0.0847 | -0.0731 |
| (Features of Academia) Funding, Job market, Promotion (factor) | -0.1044 . | -0.0163 | 0.0113 | -0.0003 | 0.0746 | 0.0965 | -0.1029 | 0.0757 | -0.01 | 0.1668 ** | 0.0225 | 0.0344 |
| (Features of Academia) Research, Autonomy (factor) | -0.0602 | -0.0232 | 0.2296 | 0.0327 | 0.0453 | -0.0425 | -0.1427 * | -0.0762 | -0.1497 | 0.1768 ** | -0.0361 | -0.1202 |
| (Features of Academia) Teaching, Mentoring (factor) | -0.0276 | 0.1047 | 0.1799 | -0.0037 | -0.0921 | 0.0529 | 0.0341 | -0.0475 | -0.3779 . | 0.1044 | 0.0446 | -0.1403 |
| (Features of Academia) Work/Life balance (factor) | -0.1534 . | 0.0306 | -0.0135 | -0.0257 | 0.1091 | 0.2224 | -0.0744 | 0.0274 | -0.1657 | 0.2195 ** | 0.0486 | -0.112 |
| Confident being independent researcher | -0.0296 | -0.0289 | -0.2088 | -0.0447 | 0.0828 | -0.0768 | -0.0036 | -0.0193 | 0.1696 | 0.0562 | -0.1217 . | 0.3541 * |

**Table S10: Follow-up analyses for regressions predicting interest.** Follow-up results for significant interactions in the final regressions reported in Tables 1 and 2. UR = underrepresented, WR = well represented. * = $p < 0.05$, ** = $p < 0.01$, *** = $p < 0.001$.

**Table S10a: Follow-ups for significant interactions in regressions predicting T2 interest**

<table>
<tr><th rowspan="2">Dependent Variable: T2 (End of PhD) Career Interest Rating</th><th colspan="6">Interaction</th></tr>
<tr><th>Gender</th><th>UR Status</th><th>Independent Variable</th><th>Moderator Groups</th><th>Group Slope</th><th>Significance of Test of Differences in Slopes</th></tr>
<tr><td rowspan="4">Academic Faculty/Research</td><td rowspan="4">Yes</td><td rowspan="4">Yes</td><td rowspan="4">PhD Faculty support, outside of institution</td><td>WR / Female</td><td>0.008</td><td rowspan="2">n.s.</td></tr>
<tr><td>WR / Male</td><td>0.037</td></tr>
<tr><td>UR / Female</td><td>0.132</td><td rowspan="2">*</td></tr>
<tr><td>UR / Male</td><td>-0.143</td></tr>
<tr><td rowspan="4">Academic Faculty/Teaching</td><td rowspan="4">Yes</td><td rowspan="4">Yes</td><td rowspan="4">PhD Faculty support, at institution</td><td>WR / Female</td><td>-0.010</td><td rowspan="2">n.s.</td></tr>
<tr><td>WR / Male</td><td>0.012</td></tr>
<tr><td>UR / Female</td><td>0.079</td><td rowspan="2">**</td></tr>
<tr><td>UR / Male</td><td>-0.210</td></tr>
<tr><td rowspan="2">Research/Non-academic</td><td rowspan="2">Yes</td><td rowspan="2">No</td><td rowspan="2">T1 interest</td><td>Female</td><td>0.536</td><td rowspan="2">**</td></tr>
<tr><td>Male</td><td>0.667</td></tr>
<tr><td rowspan="2">Science/Non-research</td><td rowspan="2">No</td><td rowspan="2">Yes</td><td rowspan="2">T1 interest</td><td>WR</td><td>0.766</td><td rowspan="2">*</td></tr>
<tr><td>UR</td><td>0.631</td></tr>
</table>

**Table S10b: Follow-ups for significant interactions in regressions predicting T3 interest**

| Dependent Variable: T3 (Current) Career Interest Rating | Interaction | | | | | |
|---|---|---|---|---|---|---|
| | Gender | UR Status | Independent Variable | Moderator Groups | Group Slope | Significance of Test of Differences in Slopes |
| Academic Faculty/Teaching | No | Yes | PhD advisor career advice | WR | 0.034 | * |
| | | | | UR | -0.107 | |
| Science/Non-research | Yes | Yes | Top 50 undergraduate institution | Not / WR / Female | 2.49 | *** |
| | | | | Not / WR / Male | 2.23 | |
| | | | | Not / UR / Female | 2.53 | n.s. |
| | | | | Not / UR / Male | 2.49 | |
| | | | | Top 50 / WR / Female | 2.38 | n.s. |
| | | | | Top 50 / WR / Male | 2.24 | |
| | | | | Top 50 / UR / Female | 2.68 | ** |
| | | | | Top 50 / UR / Male | 1.36 | |
| | Yes | No | Years it took to complete PhD | Female | 0.120 | ** |
| | | | | Male | -0.043 | |
| | Yes | Yes | Confident being independent researcher | WR / Female | -0.049 | n.s. |
| | | | | WR / Male | -0.132 | |
| | | | | UR / Female | -0.258 | * |
| | | | | UR / Male | 0.070 | |
| | No | Yes | First-author publication rate | WR | -0.029 | * |
| | | | | UR | -0.744 | |
| | Yes | No | Research, Autonomy (factor) | Female | -0.127 | * |
| | | | | Male | 0.051 | |

**Table S11: Factor Analyses.** Summary of factor analyses, which reduced 32 questions to 17 factor variables. Shaded: variables that had the highest factor loadings for that factor.

**Table S11a: Factor Analysis for Important Aspects of Career Questions**

| Explanatory Variable | Standardized Factor Loadings | | | | | | |
|---|---|---|---|---|---|---|---|
| | "Autonomy" | "Collaboration" | "Make a difference" | "Varied work" | "Ability to do job" | "Geographic location" | "Work/life balance |
| Career aspects: Low stress | -0.08 | -0.06 | -0.05 | -0.08 | -0.04 | -0.08 | 0.08 |
| Career aspects: High autonomy | 1.04 | -0.11 | -0.14 | -0.1 | -0.1 | -0.07 | -0.04 |
| Career aspects: Work-life balance | -0.02 | -0.04 | -0.03 | -0.02 | -0.04 | -0.03 | 0.97 |
| Career aspects: Personal competence/ability to do the job | -0.05 | -0.08 | -0.09 | -0.07 | 1.03 | -0.07 | -0.04 |
| Career aspects: Frequent collaboration and working with others | -0.05 | 1.04 | -0.09 | -0.08 | -0.08 | -0.07 | -0.03 |
| Career aspects: Job market (ease of entering/advancing in field) | -0.24 | -0.1 | -0.14 | -0.08 | -0.09 | -0.03 | -0.12 |
| Career aspects: Geographic location of job or career prospects | -0.05 | -0.07 | -0.07 | -0.07 | -0.07 | 0.98 | -0.03 |
| Career aspects: Job security | -0.15 | -0.17 | -0.09 | -0.17 | -0.09 | -0.15 | -0.05 |
| Career aspects: Monetary compensation (e.g. salary benefits) | -0.23 | -0.11 | -0.16 | -0.09 | -0.19 | -0.1 | -0.05 |
| Career aspects: Ability to make a difference | -0.06 | -0.09 | 1.04 | -0.08 | -0.09 | -0.06 | -0.03 |
| Career aspects: Varied, diverse work | -0.05 | -0.08 | -0.08 | 1.04 | -0.07 | -0.06 | -0.02 |
| Career aspects: Intellectually stimulating | 0.11 | 0.02 | 0.06 | 0.05 | -0.04 | -0.07 | -0.14 |
| **Summary Statistics** | | | | | | | |
| Eigenvalue/SS Loading | 1.25 | 1.18 | 1.18 | 1.17 | 1.15 | 1.03 | 0.99 |
| Proportion of Variance | 0.10 | 0.10 | 0.10 | 0.10 | 0.10 | 0.09 | 0.08 |
| Cumulative Variance | 0.10 | 0.30 | 0.20 | 0.40 | 0.49 | 0.58 | 0.66 |

**Table S11b: Factor Analysis for Salient Features of Academia Questions**

| Explanatory Variable | Standardized Factor Loadings | | | |
|---|---|---|---|---|
| | "Funding, Job market, Promotion" | "Research, Autonomy" | "Teaching, Mentoring" | "Work/Life balance" |
| Features of Academia: Compensation in academia | 0.45 | -0.03 | 0.05 | 0.08 |
| Features of Academia: Obtaining research funding | 0.82 | 0.02 | 0.01 | -0.24 |
| Features of Academia: Ability to get published | 0.42 | 0.34 | -0.15 | -0.06 |
| Features of Academia: Teaching | 0.04 | -0.21 | 0.70 | 0.00 |
| Features of Academia: Conducting research | -0.05 | 0.79 | -0.10 | 0.02 |
| Features of Academia: Academic job market in my field | 0.73 | -0.16 | 0.01 | 0.07 |
| Features of Academia: Tenure and promotion process | 0.54 | 0.04 | 0.07 | 0.08 |
| Features of Academia: Work/life balance | 0.34 | 0.07 | -0.07 | 0.50 |
| Features of Academia: Geographic location of job or career prospects | 0.42 | -0.03 | 0.03 | 0.23 |
| Features of Academia: Ability to make positive impact on society | -0.01 | 0.45 | 0.24 | 0.07 |
| Features of Academia: Mentoring the future generation of scientists | -0.03 | 0.31 | 0.61 | -0.09 |
| Features of Academia: High autonomy | -0.08 | 0.54 | 0.00 | 0.06 |
| **Summary Statistics** | | | | |
| Eigenvalue/SS Loading | 2.17 | 1.41 | 0.97 | 0.40 |
| Proportion of Variance | 0.18 | 0.12 | 0.08 | 0.03 |
| Cumulative Variance | 0.18 | 0.30 | 0.38 | 0.41 |

**Table S11c: Factor Analysis for PhD Advisor Relationship Questions**

| Explanatory Variable | Standardized Factor Loadings |
|---|---|
| | "Relationship with PhD advisor" |
| Your relationship with your (PhD) primary training advisor | 0.84 |
| My (PhD) advisor made me feel included in the lab | 0.85 |
| My (PhD) advisor appreciated my contributions | 0.93 |
| **Summary Statistics** | |
| Eigenvalue/SS Loading | 2.28 |
| Proportion of Variance | 0.76 |

**Table S11d: Factor Analysis for PhD Group Belonging Questions**

| Explanatory Variable | Standardized Factor Loadings "PhD: Belong to social community/ Department" | "PhD: Belong to intellectual community/Lab" |
|---|---|---|
| I belong/ed to the intellectual community of my (PhD) research group | -0.22 | 1.12 |
| I belong/ed to the social community of my (PhD) research group | 0.32 | 0.41 |
| I belong/ed to the intellectual community of my (PhD) department/program | 0.55 | 0.22 |
| I belong/ed to the social community of my (PhD) department/program | 1.12 | -0.23 |
| **Summary Statistics** | | |
| Eigenvalue/SS Loading | 1.56 | 1.37 |
| Proportion of Variance | 0.39 | 0.34 |
| Cumulative Variance | 0.39 | 0.73 |

**Table S11e: Factor Analysis for Postdoc Advisor Relationship Questions**

| Explanatory Variable | Standardized Factor Loadings "Relationship with postdoc advisor" |
|---|---|
| Your relationship with your (postdoc) primary training advisor | 0.88 |
| My (postdoc) advisor made me feel included in the lab | 0.88 |
| My (postdoc) advisor appreciated my contributions | 0.93 |
| **Summary Statistics** | |
| Eigenvalue/SS Loading | 2.4 |
| Proportion of Variance | 0.8 |

**Table S11f: Factor Analysis for Postdoc Group Belonging Questions**

| Explanatory Variable | Standardized Factor Loadings | |
|---|---|---|
| | "Postdoc: Belong to social community/ Department" | "Postdoc: Belong to intellectual community/ Lab" |
| I belong/ed to the intellectual community of my (postdoc) research group | -0.2 | 1.11 |
| I belong/ed to the social community of my (postdoc) research group | 0.3 | 0.46 |
| I belong/ed to the intellectual community of my (postdoc) department/program | 0.68 | 0.15 |
| I belong/ed to the social community of my (postdoc) department/program | 1.11 | -0.2 |
| **Summary Statistics** | | |
| Eigenvalue/SS Loading | 1.69 | 1.37 |
| Proportion of Variance | 0.42 | 0.34 |
| Cumulative Variance | 0.42 | 0.77 |

**References**

Benjamini, Y., & Hochberg, Y. (1995). Controlling the false discovery rate: A practical and powerful approach to multiple testing. *Journal of the Royal Statistical Society: Series B (Methodological)*, *57*(1), 289–300. doi:10.1111/j.2517-6161.1995.tb02031.x

Lenth, R. (2020). emmeans: Estimated Marginal Means, akaLeast-Squares Means (R package version 1.4.5). Computer software, N/A. Retrieved from https://CRAN.R-project.org/package=emmeans

Pollard, K. S., Dudoit, S., & van der Laan, M. J. (2005). Multiple Testing Procedures: the multtest Package and Applications to Genomics. In R. Gentleman, V. J. Carey, W. Huber, R. A. Irizarry, & S. Dudoit (eds.), *Bioinformatics and computational biology solutions using R and bioconductor* (pp. 249–271). New York, NY: Springer New York. doi:10.1007/0-387-29362-0_15

Revelle, W. (2019). psych Procedures for Psychological, Psychometric, and Personality Research (R package version 1.9.12). Computer software, Evanston, Illinois: Northwestern University. Retrieved from https://CRAN.R-project.org/package=psych
