## Supplementary material for "Factors that Influence Career Choice Among Different Populations of Neuroscience Trainees": Survey Text

This document contains a plain-text version of the survey that highlights differences across participant categories. The survey is largely the same across participants, with some minor wording changes and additional questions depending on career stage.

Annotations are as follows:

- Differences in question wording are highlighted in red text.
- Explanatory information is in [red, bracketed text].

All questions are required except for demographic questions.

Survey should be read downward, following the columns. Questions displayed across two or three columns are similar across those participant groups.

**Factors that Influence Career Choice among Neuroscience Trainees**

Public reporting burden for this collection of information is estimated to average 20 minutes, including the time for reviewing instructions, searching existing data sources, gathering and maintaining the data needed, and completing and reviewing the collection of information. The NIH Office of Extramural Research has determined that this survey and associated planned analyses fit the definition of research conducted by NIH; thus, per the 21st Century Cures Act, it is not subject to Paperwork Reduction Act requirements.

**Background**

The scientific workforce landscape has changed, and individuals are pursuing a variety of career paths after completion of doctoral and/or postdoctoral training (NIH Biomedical Research Workforce Working Group Report 2012). The National Institute of Neurological Disorders and Stroke (NINDS) would like to know more about the reasons and factors that influence individual career choices. With this survey, we seek input from current or recent trainees in the neuroscience field to help inform future training programs and initiatives to better serve the neuroscience community. The NINDS is also committed to the development of a biomedical research workforce that is representative of the diversity in American society. We strongly encourage the participation of diverse neuroscientists in this survey including minorities, persons with disabilities, those from disadvantaged backgrounds, and women.

**Eligibility**

This survey is open to current PhD trainees (including MD/PhD trainees) or those who finished their PhD in 2008 or later who are US citizens or permanent residents and who have applied for funding from NINDS or participated in an NINDS-funded program.

**Why Should You Participate?**

Your input and feedback will help us improve our current programs, develop appropriate training opportunities, and provide programmatic support for career success for current and future NINDS trainees. Participation in this survey is voluntary and should only take 20 minutes to complete.

**Privacy and Use of Information**

The information you provide will be kept private, and will not be disclosed to anyone but the researchers conducting this study, except as otherwise required by law. Responses are anonymous. All responses will be reported in aggregate.

**Contact**

If you have any questions about this survey, please contact Dr. Michelle Jones-London, Chief, Office of Programs to Enhance Neuroscience Workforce Diversity.

**Eligibility Questions**

1. Have you ever applied for funding from the National Institute of Neurological Disorders and Stroke (NINDS) or participated in an NINDS-funded program? *This **does not** include NINDS research grants awarded to your training advisor that may have supported your graduate or postdoctoral work. It **does** include applications to fellowship programs, such as the NRSA F30, F31, or F32; appointment to a training award, or T32; mentored career awards, such as the K01, K08, K22, or K99/R00; research grants you have applied for, such as the R01 or R21; receipt of a Diversity or Re-Entry Supplement; or participation in an NINDS-funded research education grant (R25), such as the Neuroscience Scholars Program, MINDS, or BRAINS.*
   - ☐ No
   - ☐ Yes

2. Are you currently enrolled in a PhD or MD/PhD program or did you earn your PhD in 2008 or later?
   - ☐ No
   - ☐ Yes

3. Are you currently a citizen or permanent resident of the United States?
   - ☐ No
   - ☐ Yes

**[Answering "No" to any of the above three questions disqualifies the participant]**

**Section 1. Career Stage**

4. What is your current career stage?
   - ☐ PhD or MD/PhD Student
   - ☐ Postdoctoral Fellow *(A postdoctoral appointment, or "postdoc," is a temporary position awarded in academe, industry, non-profit organization, or government primarily for gaining additional education and training in research.)*
   - ☐ Professional in the workforce
   - ☐ Unemployed
   - ☐ Other (please specify): ____________________

**[Skip logic is employed to tailor the survey to career stage]**

| PhD Student | Postdoctoral Fellow | Professional in the workforce/Unemployed/Other |
|---|---|---|

5. Please rate your level of interest in the following careers **when you began your PhD program**.

| | No interest | Low interest | Moderate interest | Strong Interest |
|---|---|---|---|---|
| Academic position, research focus (includes physician-scientist) | ☐ | ☐ | ☐ | ☐ |
| Academic position, teaching focus | ☐ | ☐ | ☐ | ☐ |
| Non-academic research (e.g., research in industry, biotech, or government settings) | ☐ | ☐ | ☐ | ☐ |
| Science-related, non-research (e.g., science outreach, communication, policy, advocacy, or administration) | ☐ | ☐ | ☐ | ☐ |
| Other, non-science-related | ☐ | ☐ | ☐ | ☐ |

6. Please rate your level of interest in the following careers **at PhD completion**:

| | No interest | Low interest | Moderate interest | Strong Interest |
|---|---|---|---|---|
| Academic position, research focus (includes physician-scientist) | ☐ | ☐ | ☐ | ☐ |
| Academic position, teaching focus | ☐ | ☐ | ☐ | ☐ |
| Non-academic research (e.g., research in industry, biotech, or government settings) | ☐ | ☐ | ☐ | ☐ |
| Science-related, non-research (e.g., science outreach, communication, policy, advocacy, or administration) | ☐ | ☐ | ☐ | ☐ |
| Other, non-science-related | ☐ | ☐ | ☐ | ☐ |

| PhD Student | Postdoctoral Fellow | Professional in the workforce/Unemployed/Other |
|---|---|---|

7. Please rate your **current level of interest** in the following careers:

| | No interest | Low interest | Moderate interest | Strong Interest |
|---|---|---|---|---|
| Academic position, research focus (includes physician-scientist) | ☐ | ☐ | ☐ | ☐ |
| Academic position, teaching focus | ☐ | ☐ | ☐ | ☐ |
| Non-academic research (e.g., research in industry, biotech, or government settings) | ☐ | ☐ | ☐ | ☐ |
| Science-related, non-research (e.g., science outreach, communication, policy, advocacy, or administration) | ☐ | ☐ | ☐ | ☐ |
| Other, non-science-related | ☐ | ☐ | ☐ | ☐ |

8. Since you began your PhD program, has your primary career goal changed from a research-based career to a career outside of research?
   - ☐ No, my primary career goal is still a research-based career.
   - ☐ No, my primary career goal was never a research-based career.
   - ☐ Yes, my primary career goal changed from a research-based career to a career outside of research.

| PhD Student | Postdoctoral Fellow | Professional in the workforce/Unemployed/Other |
|---|---|---|
| 9. Think about the post-training career(s) you are **currently** interested in. Which aspects of the career or work environment are most important to you? (Choose **up to 5** top reasons) | 9. Think about the post-training career(s) you are **currently** interested in. Which aspects of the career or work environment are most important to you? (Choose **up to 5** top reasons) | 9. Think about why you chose your **current** post-training career. Which aspects of the career or work environment were most important to you? (Choose **up to 5** top reasons) |

- ☐ Low stress
- ☐ High autonomy
- ☐ Work/life balance
- ☐ Personal competence/ability to do the job
- ☐ Frequent collaboration and working with others
- ☐ Job market (ease of entering/advancing in field)
- ☐ Geographic location of job or career prospects
- ☐ Job security
- ☐ Monetary compensation (e.g., salary, benefits)
- ☐ Ability to make a difference
- ☐ Varied, diverse work
- ☐ Intellectually stimulating
- ☐ Other (please specify) _____________________

10. Listed below are some features of academia that influence people's interest in becoming a faculty member. For each item please indicate how much the item either increases/d or decreases/d your desire to become a faculty member.

| | Greatly decreases/d my desire to become a faculty member | Slightly decreases/d my desire to become a faculty member | No effect on my desire to become a faculty member | Slightly increases/d my desire to become a faculty member | Greatly increases/d my desire to become a faculty member |
|---|---|---|---|---|---|
| Compensation in academia (salary, benefits) | ☐ | ☐ | ☐ | ☐ | ☐ |
| Obtaining research funding | ☐ | ☐ | ☐ | ☐ | ☐ |
| Ability to get published | ☐ | ☐ | ☐ | ☐ | ☐ |
| Teaching | ☐ | ☐ | ☐ | ☐ | ☐ |
| Conducting research | ☐ | ☐ | ☐ | ☐ | ☐ |
| Academic job market in my field | ☐ | ☐ | ☐ | ☐ | ☐ |
| Tenure and promotion process | ☐ | ☐ | ☐ | ☐ | ☐ |
| Work/life balance | ☐ | ☐ | ☐ | ☐ | ☐ |
| Geographic location of job or career prospects | ☐ | ☐ | ☐ | ☐ | ☐ |
| Ability to make positive impact on society | ☐ | ☐ | ☐ | ☐ | ☐ |
| Mentoring the future generation of scientists | ☐ | ☐ | ☐ | ☐ | ☐ |
| High autonomy | ☐ | ☐ | ☐ | ☐ | ☐ |

11. I am confident in my potential to be an independent researcher.

Strongly disagree    Disagree    Neither agree nor disagree    Agree    Strongly agree

| PhD Student | Postdoctoral Fellow | Professional in the workforce/Unemployed/Other |
|---|---|---|

These questions concern your experience during your **PhD training**.

12. Please make an overall assessment of your relationship with your **primary training advisor.**

| Very negative | Negative | Neutral | Positive | Very positive |
|---|---|---|---|---|

13. Please indicate your agreement with the following statements about your **primary PhD training advisor**.

| | Strongly disagree | Disagree | Neither agree nor disagree | Agree | Strongly agree |
|---|---|---|---|---|---|
| My advisor made me feel included in the lab | ☐ | ☐ | ☐ | ☐ | ☐ |
| My advisor appreciated my contributions | ☐ | ☐ | ☐ | ☐ | ☐ |

14. **During a PhD program**, many people can provide advice and mentorship in developing skills in a variety of areas, such as teaching, work-life balance, preparing publications and grants, and navigating institutional barriers. In the chart below, please rate the helpfulness of the support provided in these areas by the following groups.. *Please check all that apply.*

| | N/A or no guidance provided | Not helpful | Somewhat helpful | Very helpful |
|---|---|---|---|---|
| Primary training advisor | ☐ | ☐ | ☐ | ☐ |
| Faculty at primary institution | ☐ | ☐ | ☐ | ☐ |
| Faculty outside of primary institution | ☐ | ☐ | ☐ | ☐ |
| Peers (other students or postdocs) | ☐ | ☐ | ☐ | ☐ |

15. More specifically, how would you characterize the advice you have received from the following **during your PhD training** to help you plan your **future career**?

15. More specifically, how would you characterize the advice you received from the following **during your PhD training** to help you plan your **future career**?

| | N/A or no guidance provided | Not helpful | Somewhat helpful | Very helpful |
|---|---|---|---|---|
| Peers | ☐ | ☐ | ☐ | ☐ |
| Advisor | ☐ | ☐ | ☐ | ☐ |
| Department/Program | ☐ | ☐ | ☐ | ☐ |
| Institution | ☐ | ☐ | ☐ | ☐ |

| PhD Student | Postdoctoral Fellow | Professional in the workforce/Unemployed/Other |
|---|---|---|

16. These statements focus on your experiences during your PhD program. How strongly do you agree or disagree with the following statements:

| | Strongly disagree | Disagree | Neither agree nor disagree | Agree | Strongly agree |
|---|---|---|---|---|---|
| I belong/ed to the intellectual community of my research group. | ☐ | ☐ | ☐ | ☐ | ☐ |
| I belong/ed to the social community of my research group. | ☐ | ☐ | ☐ | ☐ | ☐ |
| I belong/ed to the intellectual community of my department/program. | ☐ | ☐ | ☐ | ☐ | ☐ |
| I belong/ed to the social community of my department/program. | ☐ | ☐ | ☐ | ☐ | ☐ |

17. My PhD program helped me develop/continue to develop this skill:

| | Strongly disagree | Disagree | Neither agree nor disagree | Agree | Strongly agree |
|---|---|---|---|---|---|
| Leadership | ☐ | ☐ | ☐ | ☐ | ☐ |
| Oral communication | ☐ | ☐ | ☐ | ☐ | ☐ |
| Written communication | ☐ | ☐ | ☐ | ☐ | ☐ |
| Teaching | ☐ | ☐ | ☐ | ☐ | ☐ |
| Collaboration and teamwork | ☐ | ☐ | ☐ | ☐ | ☐ |
| Data analysis, interpretation, and management | ☐ | ☐ | ☐ | ☐ | ☐ |
| Problem-solving | ☐ | ☐ | ☐ | ☐ | ☐ |
| Project management | ☐ | ☐ | ☐ | ☐ | ☐ |
| Time management | ☐ | ☐ | ☐ | ☐ | ☐ |
| Creativity/innovative thinking | ☐ | ☐ | ☐ | ☐ | ☐ |
| Interpersonal relations (manage conflict; relate to others) | ☐ | ☐ | ☐ | ☐ | ☐ |
| Ethical awareness and integrity | ☐ | ☐ | ☐ | ☐ | ☐ |
| Ability to supervise others | ☐ | ☐ | ☐ | ☐ | ☐ |
| Obtaining and negotiating the next career step | ☐ | ☐ | ☐ | ☐ | ☐ |

| | |
|---|---|
| | 18. Have you ever been a postdoctoral fellow?<br>☐ No<br>☐ Yes<br>**[Skip logic is employed. If "No," skip new two postdoctoral pages]** |

| PhD Student | Postdoctoral Fellow | Professional in the workforce/Unemployed/Other |
|---|---|---|
| | These questions concern your experience during your during your current **postdoctoral training**. | These questions concern your experience during your during your most recent **postdoctoral training experience**. |

19. Please make an overall assessment of your relationship with your current/most recent **primary postdoctoral training advisor.**

Very negative | Negative | Neutral | Positive | Very positive

20. Please indicate your agreement with the following statements about your current/most recent **primary postdoctoral training advisor**.

| | Strongly disagree | Disagree | Neither agree nor disagree | Agree | Strongly agree |
|---|---|---|---|---|---|
| My advisor made me feel included in the lab | ☐ | ☐ | ☐ | ☐ | ☐ |
| My advisor appreciated my contributions | ☐ | ☐ | ☐ | ☐ | ☐ |

21. **During postdoctoral training**, many people can provide advice and mentorship in developing skills in a variety of areas, such as teaching, work-life balance, preparing publications and grants, and navigating institutional barriers. In the chart below, please rate the helpfulness of the support provided in these areas by the following groups **during your current/most recent postdoctoral position**. *Please check all that apply.*

| | N/A or no guidance provided | Not helpful | Somewhat helpful | Very helpful |
|---|---|---|---|---|
| Primary postdoctoral advisor | ☐ | ☐ | ☐ | ☐ |
| Faculty at primary institution | ☐ | ☐ | ☐ | ☐ |
| Faculty outside of primary institution | ☐ | ☐ | ☐ | ☐ |
| Peers (other students or postdocs) | ☐ | ☐ | ☐ | ☐ |

22. More specifically, how would you characterize the guidance you have received from the following during your current/most recent **postdoctoral position** to help you plan your **future career**?

| | N/A or no guidance provided | Not helpful | Somewhat helpful | Very helpful |
|---|---|---|---|---|
| Peers | ☐ | ☐ | ☐ | ☐ |
| Advisor | ☐ | ☐ | ☐ | ☐ |
| Department/Program | ☐ | ☐ | ☐ | ☐ |
| Institution | ☐ | ☐ | ☐ | ☐ |

| PhD Student | Postdoctoral Fellow | Professional in the workforce/Unemployed/Other |
|---|---|---|
| | 23. These statements focus on your experiences during your **current/most recent postdoctoral fellowship**. How strongly do you agree or disagree with the following statements: | |

| | Strongly disagree | Disagree | Neither agree nor disagree | Agree | Strongly agree |
|---|---|---|---|---|---|
| I belong/ed to the intellectual community of my research group. | ☐ | ☐ | ☐ | ☐ | ☐ |
| I belong/ed to the social community of my research group. | ☐ | ☐ | ☐ | ☐ | ☐ |
| I belong/ed to the intellectual community of my department/program. | ☐ | ☐ | ☐ | ☐ | ☐ |
| I belong/ed to the social community of my department/program. | ☐ | ☐ | ☐ | ☐ | ☐ |

24. My **current/most recent postdoctoral fellowship** helped me develop/continue to develop this skill:

| | Strongly disagree | Disagree | Neither agree nor disagree | Agree | Strongly agree |
|---|---|---|---|---|---|
| Leadership | ☐ | ☐ | ☐ | ☐ | ☐ |
| Oral communication | ☐ | ☐ | ☐ | ☐ | ☐ |
| Written communication | ☐ | ☐ | ☐ | ☐ | ☐ |
| Teaching | ☐ | ☐ | ☐ | ☐ | ☐ |
| Collaboration and teamwork | ☐ | ☐ | ☐ | ☐ | ☐ |
| Data analysis, interpretation, and management | ☐ | ☐ | ☐ | ☐ | ☐ |
| Problem-solving | ☐ | ☐ | ☐ | ☐ | ☐ |
| Project management | ☐ | ☐ | ☐ | ☐ | ☐ |
| Time management | ☐ | ☐ | ☐ | ☐ | ☐ |
| Creativity/innovative thinking | ☐ | ☐ | ☐ | ☐ | ☐ |
| Interpersonal relations (manage conflict; relate to others) | ☐ | ☐ | ☐ | ☐ | ☐ |
| Ethical awareness and integrity | ☐ | ☐ | ☐ | ☐ | ☐ |
| Ability to supervise others | ☐ | ☐ | ☐ | ☐ | ☐ |
| Obtaining and negotiating the next career step | ☐ | ☐ | ☐ | ☐ | ☐ |

| PhD Student | Postdoctoral Fellow | Professional in the workforce/Unemployed/Other |
|---|---|---|
| | | 25. Please choose the job sector that best describes your current position.<br>☐ Academic position, research focus (includes physician-scientist)<br>☐ Academic position, teaching focus<br>☐ Non-academic research (e.g., research in industry, biotech, or government settings)<br>☐ Science-related, non-research (e.g., science outreach, communication, policy, advocacy, or administration)<br>☐ Other, non-science-related<br><br>26. What was the primary way you found your current position?<br>☐ Job posting<br>☐ Former direct advisor or supervisor<br>☐ Professional network other than direct advisor (networking, personal referral, external collaboration)<br>☐ Previous position (internship, postdoc) at same organization<br>☐ Directly contacted by employer or professional recruiter<br>☐ Self-employed/created start-up<br>☐ N/A or unemployed<br>☐ Other (please specify)<br>________________________ |

| PhD Student | Postdoctoral Fellow | Professional in the workforce/Unemployed/Other |
| --- | --- | --- |

27. Did you have research experience(s) prior to entering a PhD program?
    - ☐ No
    - ☐ Yes (how many years, total? Round to the nearest year) _______________

28. Did you participate in a formal post-baccalaureate research program after your undergraduate degree?
    - ☐ No
    - ☐ Yes (how many years, total?  Round to the nearest year) __________________

29. Did you earn a Master's degree in a biomedical research discipline **prior** to entering a PhD program?
    - ☐ No
    - ☐ Yes

30. Please indicate your **undergraduate** degree institution from one of the following dropdown menus. (If you don't see your institution, please check under "The" or alternative names, such as "State University of New York System, Stony Brook University"; if your institution is not listed, please write it below)

- A – Fa
  [List of institutions]
- Fe – Ni
  [List of institutions]
- No – To
  [List of institutions]
- Tr – Y
  [List of institutions]

If your undergraduate institution was not listed in one of the dropdown menus above, please write it here: __________________

**[List of baccalaureate-granting institutions taken from The Carnegie Classification of Institutions of Higher Education and slightly edited]**

**PhD Student:**

31. Please indicate the institution from which you will receive your **doctoral degree** using one of the following dropdown menus (If you don't see your institution, please check under "The" or alternative names, such as "State University of New York System, Stony Brook University"; if your institution is not listed, please write it below)

- A – L
  [List of institutions]
- M – T
  [List of institutions]
- U – Y
  [List of institutions]

- If your doctoral institution was not listed in one of the three dropdown menus above, please write it here: _________________________

**Postdoctoral Fellow / Professional in the workforce/Unemployed/Other:**

31. Please indicate your **doctoral degree** institution using one of the following dropdown menus (If you don't see your institution, please check under "The" or alternative names, such as "State University of New York System, Stony Brook University"; if your institution is not listed, please write it below)

- A – L
  [List of institutions]
- M – T
  [List of institutions]
- U – Y
  [List of institutions]

32. If your doctoral institution was not listed in one of the three dropdown menus above, please write it here: ____________________

33. How much time did it take you to complete your PhD? (Round to the nearest year) _____________________

**[List of doctoral-granting institutions taken from The Carnegie Classification of Institutions of Higher Education and slightly edited]**

**PhD Student:**

34. Are you currently enrolled in an MD/PhD program?
    - ☐ No
    - ☐ Yes

| PhD Student | Postdoctoral Fellow | Professional in the workforce/Unemployed/Other |
|---|---|---|
| 35. What year of your PhD program are you currently in?<br>☐ First year<br>☐ Second year<br>☐ Third year<br>☐ Fourth year<br>☐ Fifth year<br>☐ Sixth year<br>☐ Seventh year<br>☐ Eighth year<br>☐ Ninth year or later<br><br>36. Please indicate your expected year of PhD degree completion<br>☐ 2017<br>☐ 2018<br>☐ 2019<br>☐ 2020<br>☐ 2021<br>☐ 2022<br>☐ 2023<br>☐ 2024 or later<br><br>37. In what discipline will you obtain your PhD?<br>☐ Biochemistry<br>☐ Bioengineering<br>☐ Bioinformatics<br>☐ Biological Sciences<br>☐ Biostatistics, Epidemiology, Public Health, Clinical Sciences<br>☐ Cellular/Molecular Biology<br>☐ Genetics<br>☐ Microbiology and Immunology<br>☐ Neuroscience<br>☐ Pathology<br>☐ Pharmacology/Toxicology<br>☐ Physiology<br>☐ Psychology<br>☐ Other (please specify): ___________________ | 35. Year of PhD conferral<br>☐ 2015<br>☐ 2014<br>☐ 2013<br>☐ 2012<br>☐ 2011<br>☐ 2010<br>☐ 2009<br>☐ 2008<br>☐ 2007 or earlier [Disqualification]<br><br>36. How many years did it take you to complete your PhD? (Round to the nearest year) ___________________<br><br>37. In what discipline did you obtain your PhD?<br>☐ Biochemistry<br>☐ Bioengineering<br>☐ Bioinformatics<br>☐ Biological Sciences<br>☐ Biostatistics, Epidemiology, Public Health, Clinical Sciences<br>☐ Cellular/Molecular Biology<br>☐ Genetics<br>☐ Microbiology and Immunology<br>☐ Neuroscience<br>☐ Pathology<br>☐ Pharmacology/Toxicology<br>☐ Physiology<br>☐ Psychology<br>☐ Other (please specify): ___________________ | |

| PhD Student | Postdoctoral Fellow | Professional in the workforce/Unemployed/Other |
|---|---|---|
| | 38. How many postdoctoral positions (including your current position) have been a part of your training?<br>☐ 1<br>☐ 2<br>☐ 3<br>☐ More than 3<br><br>39. Please indicate the total time you have spent in postdoctoral training, including all postdoctoral positions you've held (current and previous): (Round to the nearest year) ________<br><br>40. Please use the dropdown menus below to indicate the institution at which you are currently receiving post-doctoral training. (If you don't see your institution, please check under "The" or alternative names, such as "State University of New York System, Stony Brook University"; if your institution is not listed, please write it below)<br><br>• A – L [List of institutions] • M – T [List of institutions] • U – Y [List of institutions]<br><br>• If your postdoctoral institution was not listed in one of the three dropdown menus above, please write it here: ___________________ | 38. How many postdoctoral experiences were part of your training?<br>• 0<br>• 1<br>• 2<br>• 3<br>• More than 3<br>**[If 0, skip to next page]**<br><br>39. Please indicate the total time you spent in postdoctoral training, including all postdoctoral positions you've held: (Round to the nearest year) _______________________<br><br>40. Please use the dropdown menus below to indicate the institution at which you **most recently received post-doctoral training**. (If you don't see your institution, please check under "The" or alternative names, such as "State University of New York System, Stony Brook University"; if your institution is not listed, please write it below)<br><br>• A – L [List of institutions] • M – T [List of institutions] • U – Y [List of institutions]<br><br>• If your postdoctoral institution was not listed in one of the three dropdown menus above, please write it here: ________________ |
| | **[List of institutions taken from The Carnegie Classification of Institutions of Higher Education and slightly edited]** | |

| PhD Student | Postdoctoral Fellow | Professional in the workforce/Unemployed/Other |
|---|---|---|

41. How many **total** peer-reviewed publications are you an author on? ________________

42. How many **first author** peer-reviewed publications do you have? _______________

43. Has your research or training ever been supported by funding from NIH? *(This includes NRSA fellowships and training grant appointments, supplements, career development (K) awards, and NIH-funded diversity programs. This does not include R series or other grants awarded to your training advisor that may have supported your undergraduate, graduate, or postdoctoral work.)*
    - ☐ No
    - ☐ Yes
    - ☐ Unsure

    **[Skip logic is employed, skip next page if "No"]**

44. Please indicate each of the NIH funding types that supported your research or training. If you are unsure, give your best guess. For all grants except institutional training grants (T32), please indicate **only grants awarded to you as principal investigator.** *Indicate all that apply.*

| | Awarded/Appointed |
|---|---|
| F30 NRSA Individual Fellowship | ☐ |
| F31 NRSA Individual Fellowship | ☐ |
| F31 Diversity NRSA Individual Fellowship | ☐ |
| F32 NRSA Individual Fellowship | ☐ |
| T32 (Predoctoral) Institutional Training Grant Appointment | ☐ |
| T32 (Postdoctoral) Institutional Training Grant Appointment | ☐ |
| K22 Career Transition Award | ☐ |
| K22 Diversity Career Transition Award | ☐ |
| K99/R00 Pathway to Independence Award | ☐ |
| K01 Career Development Award | ☐ |
| K01 Diversity Career Development Award | ☐ |
| K08 Career Development Award | ☐ |
| K23 Career Development Award | ☐ |
| R01 Research Project Grant | ☐ |
| R03 Diversity Small Grants for New Investigators | ☐ |
| R15 Research Grant | ☐ |
| R21 Research Grant | ☐ |
| Other (e.g., institutional grants such as U54, P50, P20; please specify) | ☐ |

Do not indicate R series or other grants awarded to your training advisor that may have supported your undergraduate, graduate, or postdoctoral work.

| PhD Student | Postdoctoral Fellow | Professional in the workforce/Unemployed/Other |
|---|---|---|

45. Please indicate which, if any, of these NIH diversity programs you have participated in or been supported by: *Please check all that apply.*

- ☐ None
- ☐ Building Infrastructure Leading to Diversity (BUILD) Initiative
- ☐ MARC, MBRS, RISE, IMSD or other NIGMS undergraduate/graduate programs
- ☐ NIH Diversity Supplement for High School
- ☐ NIH Diversity Supplement for Undergraduate
- ☐ NIH Diversity Supplement for Postbaccalaureate
- ☐ NIH Diversity Supplement for Predoctoral Students
- ☐ NIH Diversity Supplement for Postdoctoral Trainees
- ☐ NIH Diversity Supplement for Faculty
- ☐ NIH Supplement to Promote Re-Entry
- ☐ National Research Mentoring Network (NRMN)
- ☐ NIH Blueprint ENDURE
- ☐ SfN Neuroscience Scholars Program
- ☐ Broadening the Representation of Academic Investigators in Neuroscience (BRAINS)
- ☐ Mentoring Institute for Neuroscience Diversity Scholars (MINDS)
- ☐ Summer Program in Neuroscience, Ethics, and Survival (SPINES)
- ☐ Research Enhancement Awards (SC1, SC2, SC3)
- ☐ Specialized Neuroscience Research Program (SNRP)
- ☐ Other *(please specify)* ________________________

46. Please indicate your career stage(s) while supported by NIH: *Check all that apply*

- ☐ High School Student
- ☐ Undergraduate Student
- ☐ Post-Baccalaureate
- ☐ Master's Student
- ☐ PhD Student
- ☐ Medical Student
- ☐ Postdoctoral Fellow
- ☐ Junior Faculty
- ☐ Independent Investigator
- ☐ Other (please specify)__________________________

47. How did NIH funding impact your research training? Were there effects beyond monetary contribution? _________________________________

| PhD Student | Postdoctoral Fellow | Professional in the workforce/Unemployed/Other |
| --- | --- | --- |

**Final Comments**

48. The goal of this survey is to learn more about early-stage scientists' career choices. In a few sentences, please tell us the "why" of your career choice or personal scientific journey.

49. If you have any additional comments on the usefulness and availability of career development resources; factors that were particularly helpful or obstacles you encountered in planning your career path; or if there was anything about your training that we didn't capture in this survey, please detail it here.

| PhD Student | Postdoctoral Fellow | Professional in the workforce/Unemployed/Other |
|---|---|---|

**Please tell us about yourself. (Optional)**

50. Age range
    - ☐ 20-24
    - ☐ 25-29
    - ☐ 30-34
    - ☐ 35-39
    - ☐ 40-44
    - ☐ 45-50
    - ☐ 51 or older
    - ☐ Prefer not to answer

51. What is your gender?
    - ☐ Female
    - ☐ Male
    - ☐ Other (please specify) ___________________
    - ☐ Prefer not to answer

52. Do you consider yourself to be Hispanic or Latino?
    - ☐ No
    - ☐ Yes
    - ☐ Prefer not to answer

53. What do you consider to be your race? *Select one or more*
    - ☐ American Indian or Alaska Native
    - ☐ Asian
    - ☐ Black or African American
    - ☐ Native Hawaiian or Other Pacific Islander
    - ☐ White
    - ☐ Prefer not to answer

54. Do you have a disability as defined as a physical or mental impairment that substantially limits one or more major life activities?
    - ☐ No
    - ☐ Yes
    - ☐ Prefer not to answer

| PhD Student | Postdoctoral Fellow | Professional in the workforce/Unemployed/Other |
|---|---|---|

55. Are you the first person or among the first generation in your immediate family to graduate from a 4-year college (e.g. neither of your parents/guardian(s) graduated from college)?
    - ☐ No
    - ☐ Yes
    - ☐ Prefer not to answer

56. Which of the following best represents how you think of yourself?
    - ☐ Lesbian, gay, homosexual, bisexual, or other sexual minority
    - ☐ Straight or heterosexual
    - ☐ Other (please specify)________________
    - ☐ Prefer not to answer

57. Which degrees do you currently hold? *Select one or more*
    - ☐ AS/AA (Associate's degree)
    - ☐ BA/BS (Bachelor's of Arts/Science)
    - ☐ MA/MS (Master's of Arts/Science)
    - ☐ MPH (Master's of Public Health)
    - ☐ MBA (Master's of Business Administration)
    - ☐ Other Master's degree
    - ☐ PhD (Doctor of Philosophy) or equivalent, such as DSc (Doctor of Science)
    - ☐ Other doctoral degree (EdD, Doctor of Education; PsyD, Doctor of Psychology, etc.)
    - ☐ DVM (Doctor of Veterinary Medicine)
    - ☐ MD (Doctor of Medicine)
    - ☐ DO (Doctor of Osteopathic Medicine)
    - ☐ JD (Juris Doctor)
    - ☐ Other professional degree (DDS, Doctor of Dental Surgery; OD, Doctor of Optometry etc.)
    - ☐ Other (please specify) _____________________________

### Disqualification Text

Thank you for your interest in completing our survey.

This survey is only open to current PhD trainees or those who finished their PhD in 2008 onward, who are US citizens or permanent residents, and who have applied for funding from NINDS.
